# Quantitative evaluation of internal clustering validation indices using binary datasets

**DOI:** 10.1101/2023.08.09.552566

**Authors:** Naghmeh Pakgohar, Attila Lengyel, Zoltán Botta-Dukát

**Affiliations:** Centre for Ecological Research, Institute of Ecology and Botany, Alkotmány u- 2-4, Vácrátót, H- 2163, Hungary

**Keywords:** Geometric indices, Non-geometric indices, Internal indices, Clustering

## Abstract

Different clustering methods often classify the same dataset differently. Selecting the ‘best’ clustering solution out of a multitude of alternatives is possible with cluster validation indices. The behavior of validity indices changes with the structure of the sample and the properties of the clustering algorithm. Unique properties of each index cause increasing or decreasing performance in some conditions. Due to the large variety of cluster validation indices, choosing the most suitable index concerning the dataset and clustering algorithms is challenging. We aim to assess different internal clustering validation indices. In the present paper, the validity indices consist of geometric and non-geometric methods. For this purpose, we applied simulated datasets with different noise levels. Each dataset was repeated 20 times. Three clustering algorithms with Jaccard dissimilarity are used, and 27 clustering validation indices are evaluated. The results provide a reliability guideline for the selection cluster validity indices.

## Introduction

Cluster analysis is a standard procedure in multivariate data analysis that discovers useful dataset partitions in a purely unsupervised manner (Hennig, 2015). It is designed to group heterogeneous data objects into more homogenous subgroups (Tarekegn et al., 2020). Numerical clustering algorithms have been used for decades in vegetation science (Peet & Roberts, 2013). It is often assumed (although sometimes only implicitly) that there is only a single “true” clustering for a given dataset, and cluster analysis aims to find that clustering (Akhanli & Hennig, 2020). On the other hand, applying different clustering methods to the same dataset results in various classifications. Even if the clustering algorithm is fixed, the grouping depends on the number of clusters. According to this logic, two fundamental questions must be addressed: 1) Which algorithm can achieve the best classification? 2) What is the optimal number of clusters? (Xu & Wunsch, 2009) The role of cluster validation is to answer these questions. Cluster validation is the procedure of assessing how good a classification is, both in absolute terms and in comparison with other classifications (Akhanli & Hennig, 2020). Cluster validation is typically done by calculating a cluster validation index (hereafter denoted by CVI) that measures the “goodness” of the classification.

What do we mean when we talk about the “goodness” of a classification? The answer partly depends on whether we want to measure the “goodness” of a group, a partition, or a hierarchical classification (Sousa & Tendeiro, 2005). Still, most CVIs quantify the goodness of partitions, and we also focus on this problem. Hennig (2015) concluded that it depends on the clustering goal which partition seems the best one. That is, in the assessment of partitions, different properties of clusters can be considered, including preserving distance information in the partition, compactness, connectedness, and separation of groups, the robustness of results, stability against data perturbation, replicability, interpretability, and predictive power for external variables (Handl et al., 2005). Each CVI focuses on one or some of these aspects. Therefore, users are advised to look at several CVIs to compare partitions (Akhanli & Hennig, 2020).

CVIs are often divided into two types, internal and external indices. The internal validation indices measure the quality of partitions based on the variables used for clustering (Gagolewski et al., 2021). Internal validation techniques can be subdivided into two subgroups. Geometric indices rely on the distances between sample units in the space defined by the internal variables. In contrast, non-geometric indices measure classification effectiveness with respect to the distribution of the variables within and across groups (Aho et al., 2008). In contrast to internal indices, external validation methods assess partitions based on information not used for creating the classification. Often, but not exclusively, this information is the true grouping of the same dataset (called ‘ground truth’ or ‘gold standard’), and its similarity to the actual classification serves as an external CVI (Arbelaitz et al., 2013).

There were several attempts to evaluate cluster validation indices (Bezdek & Pal, 1998; Dubes, 1987; Günter & Bunke, 2003; Gurrutxaga et al., 2011; Maulik & Bandyopadhyay, 2002; Milligan & Cooper, 1985; Vendramin et al., 2010). Almost all of them used datasets containing points spreading in Euclidean space. However, in real ecological studies, the space of measured variables is often not Euclidean. Thus the Euclidean distance between communities, as well as the centroids, the sum of squares (SQ), and variances of the groups are not meaningful measures. A plethora of indices is available for expressing dissimilarity between sample units in non-Euclidean space (Faith et al., 1987; Podani, 2000). An often-used example where we can encounter non-Euclidean space is the application of binary data (e.g., species presence/absence). The only effort to evaluate CVIs on binary data has been done by Dimitriadou et al. (2002) who used datasets with twelve variables. However, the number of variables (species) is typically much higher in ecology, and this difference may influence the behavior of CVIs. The main point of their evaluation of CVIs was comparing the number of clusters in the best partition according to the CVI with the true number of clusters. In contrast, Vendramin et al. (2010) pointed out that the selected partition may considerably differ from the true partition, even if the number of clusters is the same. To override this shortcoming, different alternative approaches were proposed for evaluating CVIs (Botta-Dukát, 2023; Gurrutxaga et al., 2011; Vendramin et al., 2010). However, to the best of our knowledge, these have not yet been applied to binary datasets. In vegetation science, several non-geometric CVIs have been developed for binary data (Aho et al., 2008; Tichý et al., 2010) but have not yet been evaluated. This paper aims to fill these knowledge gaps by comparing twenty-seven internal CVIs (seventeen geometric and ten non-geometric indices) that provide a multivariate assessment covering different aspects of cluster validity. We choose only internal CVIs that may be useful for binary data. Thus, we excluded CVIs that apply concepts related to Euclidean space, like centroids or variance. We also did not consider validation approaches based on the stability or robustness of the results. Since occurrence of species in the communities is the most typical example of binary data in ecology, we call variables to species and objects (points) to communities. It does not mean that our results are restricted to these type of data; they are valid to any type of binary data.

## Methods

### Overview of the analysis

In evaluating CVIs, we followed the approach proposed by Botta-Dukát (Botta-Dukát, 2023). The main steps were the following:

1. Artificial datasets were created with predefined grouping. Hereafter, this grouping is referred to as ground truth. The datasets were initially generated to be perfectly clustered according to the ground truth, then different amounts of noise were added to them in order to increase within- and decrease between-group variation.
2. Each dataset was classified by different algorithms, resulting in partitions where the number of groups ranged from 2 to twice the number of groups in the ground truth.
3. Each CVI was calculated for each partition, and the optimal partition according to each CVI (i.e., the partition with the highest or lowest value of the CVI) was selected.
4. Similarity to ground truth was calculated for each partition using the adjusted Rand index (L. Hubert & Arabie, 1985).
5. Performance of a CVI on a dataset was characterized by the similarity of the optimal partition based on the CVI to the ground truth divided by its observed maximum.
6. Performances of CVIs were compared by the Friedman test with Shaffer familywise error correction (Arbelaitz et al., 2013).

First, the last step was done by including all partitions irrespectively to noise level and clustering algorithms. Then it was repeated for datasets with each noise level and partitions generated by each clustering algorithm separately. See the following sections for details on generating datasets, the applied clustering algorithms, and the evaluated CVIs.

All analyses were done in R environment (version 4.2.2; R Core Team, 2022), using cluster (Maechler et al., 2022), exreport (Arias & Cozar, 2016), ggplot2 (Wickham, 2016), ggpubr (Kassambara, 2022), mclust (Scrucca et al., 2016), multcompView (Graves et al., 2019), optpart (D. W. Roberts, 2020), and vegan (Oksanen et al., 2022) packages.

### Datasets and clustering algorithm

For creating datasets with different noise levels, we applied an algorithm similar to what was used by Tichy et al. (2011) and Lengyel et al. (2018). In the first step, noise-free datasets were generated. Each simulated dataset consisted of ten a priori groups. In the case of equal-sized clusters, each group consisted of 20 communities; thus, the dataset consisted of 200 communities. When cluster sizes were set to be unequal, the first group contained 100 communities, and the other groups had 20 communities, with a total of 280 communities. The number of species was 100 in each dataset. Each group had ten characteristic species that occurred in all communities of one specific group and none of the communities outside that group. Since there were ten groups, each species was a characteristic species of one and only one group. It resulted in a perfectly clustered data structure. Then, random noise was added to the data by the swapping algorithm (Gotelli & Entsminger, 2001). This algorithm mixes zeros and ones in the matrix without changing the row and column totals (i.e., species richness of communities and species’ frequencies). Preliminary experiments showed that clustering algorithms still almost completely recovered the original group structure after 20 000 swaps. After 60 000 swaps, the group structure was hardly detectable, but the occurrence pattern was not entirely random. Therefore, we applied three noise levels to evaluate CVIs; 20 000, 40 000, and 60 000 swaps. Combining two types of cluster sizes (equal and unequal) and three noise levels resulted in six datasets. All types were replicated 20 times.

Each dataset was classified using three algorithms: the partitioning around medoids method (PAM), beta-flexible clustering with β = −0.25, and an unweighted pair group method (UPGMA) (Kaufman & Rousseeuw, 1990). In all algorithms, the complement of Jaccard similarity, a commonly used dissimilarity measure for presence-absence data in vegetation science (Podani, 2000), was used to measure dissimilarity. PAM was run independently for each number of groups in the same range. Dendrograms created by beta-flexible and UPGMA clustering were cut at different heights to result in partitions with the number of groups ranging from 2 to 20.

### Clustering validity indices (CVI)

This section describes the twenty-seven CVIs considered in this study, starting with geometric indices and following with non-geometric ones. Most of them are “optimization-like criteria” in the sense that when plotting their values against the number of clusters, the maximum/minimum indicates the optimal number of clusters (Vendramin et al., 2010). Some CVIs monotonically change with an increasing number of clusters in a hierarchical classification. Although Vendramin et al. (2010) proposed a method for converting these “difference-like criteria” to “optimization-like” ones, it cannot be applied if there are two clusters. Therefore, in our opinion, difference-like criteria should be used only for partitions with the same (a priori known) number of clusters. Thus, they are mentioned here but not included in the comparison.

The first group of geometric indices is based on the concept that a classification is good if it maps the information in the distance matrix as accurately as possible. This is achieved if the original distances and the binary matrix in which each entry indicates whether a given pair of objects belongs to the same cluster (0) or not (1) show maximal correlation.

Let us denote the distance between communities *i* and *j* by *d_ij_*. Its complement *s_ij_*=1-*d_ij_/d_max_* is the similarity between communities (*d_max_* is the theoretical maximum of the distance). Let us define a distance based on a partition by the following:

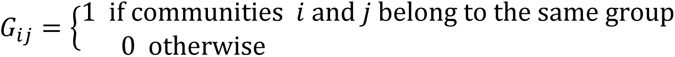

Upper (or lower) half-matrices (without the diagonal) of *d*, *G*, and *s* matrices can be treated as vectors denoted by δ, Δ and σ, respectively, ^δ̅^ and ^Δ̅^ are the means of the two vectors.

*Point-biserial correlation* (Milligan & Cooper, 1985) is the linear correlation between δ and Δ:

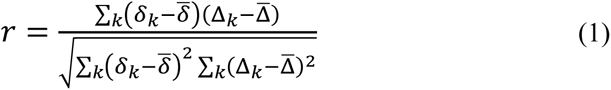

Since the vector Δ contains only 0 and 1 values, coding within-cluster (Δ_*k*_ = 0) and between-cluster distances (Δ_*k*_ = 1), point-biserial correlation can be expressed in another, more meaningful form. Let us denote the number of within- and between-cluster distances by n_W_ and n_B_, respectively. Note, *n*_*B*_ = ∑_*k*_ Δ_*k*_ and *n*_*w*_ = ∑_*k*_(1 − Δ_*k*_). Let us denote the mean within- and between-group distances by d_W_ and d_B_, respectively. Then point-biserial correlation can be expressed as

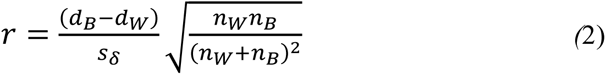

Where *s*_δ_ is the standard deviation of distances:

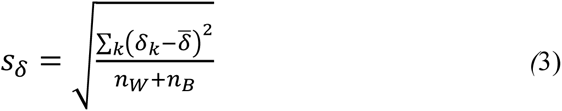

Thus, we can say that the point-biserial correlation is the difference between the means of between-cluster and within-cluster distances relative to the standard deviation of distances multiplied by the geometric mean of the proportion of within- and between-cluster distances.

Instead of linear correlation, rank correlations can also be applied. They are based on the number of concordant and discordant pairs of the elements of compared vectors. Two elements (*k* and *l*) of these vectors are concordant when δ_k_> δ_l_ and Δ_k_> Δ_l_, or δ_k_< δ_l_ and Δ_k_<Δ_l_. They are discordant when δ_k_> δ_l_ and Δ_k_< Δ_l_, or δ_k_< δ_l_ and Δ_k_>Δ_l_. *C* and *D* denote the number of concordant and discordant pairs, respectively.

*Baker & Hubert* (1975) proposed using Goodman-Kruskal’s gamma index. It can be calculated from the number of concordant and discordant pairs:

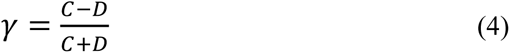

It ranges from -1 (the lowest within cluster distance is higher than the highest between cluster index) to 1 (the highest within cluster distance is lower than the lowest between cluster index).

The *G(+)* index (Rohlf, 1974) is a criterion based on the relationships between discordant pairs of objects. Specifically, it is defined as the proportion of discordant pairs with respect to the total number of pairs. It is defined as:

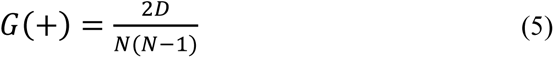

where *N* is the number of objects in the classification. A good partition has a small value for *D* and, consequently, small values for *G(+)*.

The *tau* index is Kendall’s rank correlation (Kendall, 1945) between δ and Δ vectors. It is another criterion that can be written in terms of the number of concordant (*C*) and discordant (*D*) pairs of objects (Rohlf, 1974). Contrary to Goodman-Kruskal’s gamma, it considers the number of ties (i.e., pairs with equal values):

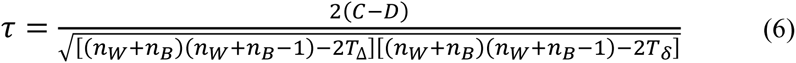

where:

*T*_δ_ and *T*_Δ_ are the numbers of tied pairs in vector δ and Δ, respectively, and

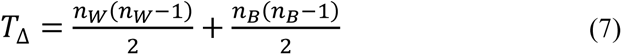

If the rank correlation is maximal, any within-group distance is lower than the lowest between-group distance.

We have already seen that the point-biserial correlation is high if the mean of between-cluster distances is much higher than that of within-cluster distances. Following this logic, the mean or sum of within-cluster and between-cluster distances can be compared. The *McClain & Rao* index is the ratio of the means of within-cluster and between-cluster distances (Good, 1982; McClain & Rao, 1975):

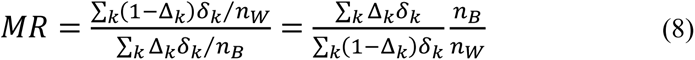

The *PARTANA* statistic (D. Roberts, 2005) calculates the same ratio for similarities:

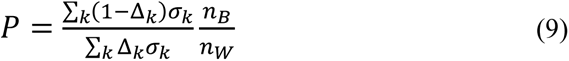

Both indices range from zero to infinity. The low value of the McClain & Rao index and the high value of PARTANA indicate a good partition. They are “difference-like” criteria; therefore, they were not involved in our comparison.

Hubert and Levine (1976) proposed using the sum of within-cluster distances after re-scaling to the zero-one interval. For rescaling, first, its possible minimum is subtracted from the sum of within-cluster distances, and then this difference is divided by the possible range:

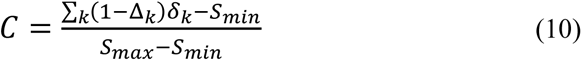

where *S*_*min*_ and *S*_*max*_ are the possible minimum and maximum of the sum of the within-cluster distances

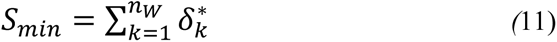

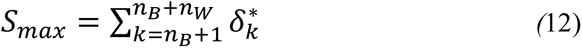

and δ^∗^ is the vector δ sorted in ascending order.

Another group of geometric indices combines the separation of groups with their compactness or connectedness. High compactness means small within-group dissimilarities, while high connectedness means that each object is close to some other (but not necessarily all) objects from the same group. The separation is maximal when the distances between the groups are maximal. Hereafter compactness and connectedness together will be called homogeneity of the cluster. The classification algorithms also optimize these properties (for example, the k-means and the average linkage methods maximize the compactness of the groups, while the single linkage is based on the connectedness). Still, most of them focus on only one property. An exception is Podani’s (1989) global optimization algorithm, which considers compactness and separation. As the number of groups increases, even in the case of random grouping, the compactness of the groups increases, but their separation decreases. Therefore, the majority of methods combine these aspects to obtain a curve that passes through a maximum (or minimum) by increasing the number of groups.

Let us denote group *k* by set ℂ_*k*_ (i.e., each group is a set of communities). The heterogeneity of this cluster is *H_k_*, its size (i.e., number of communities belonging to the groups) is *n_k_*, while the separation of cluster *k* and *l* is *S_kl_*.

The family of *Dunn index* (Dunn, 1974) measures goodness by the ratio of separation between the two closest groups and heterogeneity of the less compact (or less connected) group.

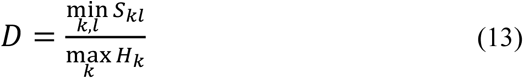

It has several versions depending on the actual measure of separation and heterogeneity (Bezdek & Pal, 1998; Gagolewski et al., 2021; Pal & Biswas, 1997). Dunn’s original formulas for heterogeneity and separation was

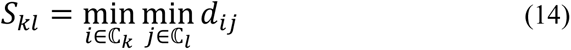

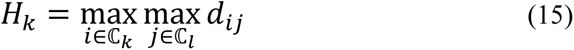

Since it is sensitive to the outlier values, alternative definitions can also be used (Bezdek & Pal, 1998; Gagolewski et al., 2021; Pal & Biswas, 1997), for example:

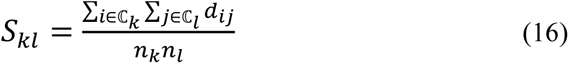

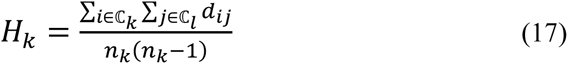

Hereafter, we refer to the Dunn index using the original definition (equations **Hiba! A hivatkozási forrás nem található.**) and **Hiba! A hivatkozási forrás nem található.**)) as Dunn_min_max_ while using the alternative definition (equations **Hiba! A hivatkozási forrás nem található.**) and **Hiba! A hivatkozási forrás nem található.**)) as *Dunn_mean_mean_*.

The Dunn index is based on the highest separation and lowest heterogeneity. Davies and Bouldin (1979) and Popma et al. (1983) proposed indices that first calculate the goodness of each cluster, and then the classification is characterized by the mean of these values.

In the *Popma* index, the goodness of cluster *k* (*Q_k_*) is the ratio of its heterogeneity and separation from the nearest clusters:

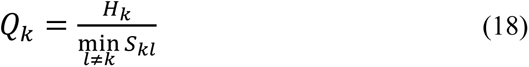

using mean distances as the measure of heterogeneity and separation, as in equations **Hiba! A hivatkozási forrás nem található.**) and **Hiba! A hivatkozási forrás nem található.**). The unweighted Popma index is the arithmetic mean of cluster goodness values, while its weighted version uses the size of clusters for weighting.

*Davies & Bouldin* (1979) suggested that when the separation of two clusters is evaluated, the heterogeneity of both clusters should be considered. Therefore, they measured the goodness of a cluster by

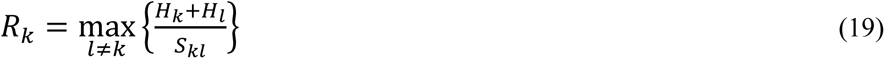

In their paper, to demonstrate the properties of this new index, they measured heterogeneity and separation by mean Minkowsky distance (a generalization of the Euclidean distance) of points from cluster centroids and between centroids. However, the general definition allows measuring them in other ways. In this paper, we used formulas in equations **Hiba! A hivatkozási forrás nem található.**) and **Hiba! A hivatkozási forrás nem található.**) to define separation and heterogeneity.

The *silhouette width* index (Kaufman & Rousseeuw, 1990; Lengyel & Botta-Dukát, 2019; Rousseeuw, 1987) is calculated for each communities, and then it can be averaged for each group or the whole partition. The silhouette width of a community is the difference between the distance of the focal community to the closest outgroup (separation) and its distance to the rest of its own group (compactness, connectedness) rescaled between -1 and 1:

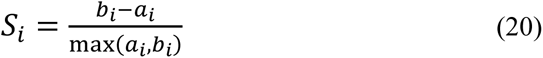

where:

*a_i_* = mean distance of community i to other communities belonging to the same group
*b_i_* = mean distance of community i to communities belonging to the closest group (i.e., a group with the lowest mean distance)

In the original definition of Rousseeuw (1987), the “mean” is an arithmetic mean that prefers spherical clusters. To relax this preference for this specific cluster shape, Lengyel & Botta-Dukát (2019) proposed using the generalized mean instead of the arithmetic means in calculating *a_i_* and *b_i_*. The generalized mean has a parameter (*q*, the degree of mean) that influences its sensitivity to small and large values. The original silhouette width using the arithmetic mean is identical to the *q* = 1 case of the generalized silhouette width. Lower *q* values increase the sensitivity to compactness, while higher values increase the sensitivity to connectedness. In this study, beyond the original silhouette width (which corresponds to *q* = 1), we applied two values of degree (-1 and 0) and referred to them as *gensil(-1)* and *gensil(0)*, respectively. A community is correctly classified if it is, on average, closer to other communities in its own group than communities in any other group. The silhouette of such communities is positive. Feoli et al. (2006) proposed using the proportion of these communities, the *Proportion of correct classifications*, as a measure of goodness.

From the similarity between communities, we can calculate the similarity between groups. If the groups are well-separated, their similarities to each other are close to zero. In the ideal case, the non-diagonal elements of their similarity matrix are zero (i.e., any group is maximally different from any other). In this case, all eigenvalues of the similarity matrix are equal, while eigenvalues become different when the groups are not well-separated. Therefore, Feoli et al. (2009) proposed using the evenness of eigenvalues as a measure of partition quality.

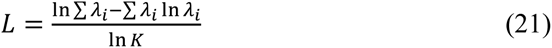

where:

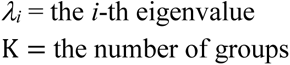

The non-geometric indices do not use the distances between objects but the characteristics of the groups used for interpreting them. In a good classification, each group has its own distinctive features. For binary data, such a feature is the proportion of presence (relative frequency) of each variable (species).

Before showing the non-geometric indices, let us introduce some notations:

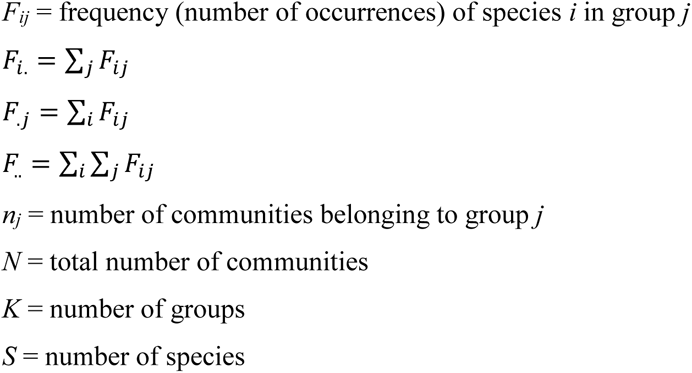

The interpretation of the group is more straightforward if each species occurs in it with either high or very low frequency. *ISAMIC* (Aho et al., 2008) quantifies this aspect of goodness:

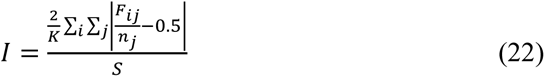

ISAMIC ranges from zero to one. A higher value indicates better classification. It is a “difference-like” criterion: ISAMIC increases or remains unchanged but cannot decrease when a group is divided into two parts. Therefore, it is maximal if each group contains only a single community.

Aho et al. (2008) suggested calculating the similarity between pairs of groups based on frequencies by the Morisita index,

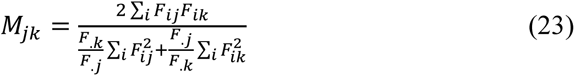

and using mean similarity as a cluster validation index. It is also a “difference-like” criterion: its value tends to increase with the number of clusters.

Species by group table of frequencies can be treated as a contingency table which allows calculating information theory measures of partition’s goodness (Feoli & Lausi, 1980). Mutual information of this contingency table is:

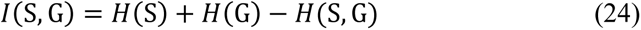

where *H*(S) is the entropy of the vector of species frequencies, *H*(G) is the entropy of group sizes, and *H*(S, G) is the joint entropy of the contingency table. Using Shannon entropy, they can be calculated as

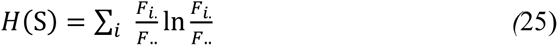

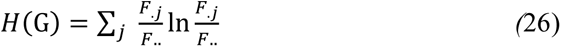

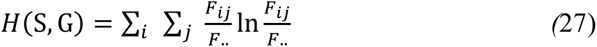

Mutual information is zero if knowledge of group membership does not decrease the uncertainty of species composition (i.e., we cannot better predict the list of occurring species when we know which group the community belongs to than in lack of this information). On the other hand, it is maximal if knowledge of the group membership allows predicting species composition without any uncertainty. The maximum mutual information increases with the increasing number of groups; therefore, only its standardized version could be used as CVI. Using the fact that *I*(S, G) ≤ *H*(S, G), Feoli and Lausi (1980) proposed using the relative divergence:

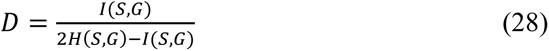

that ranges from zero to one, and a higher value indicates better classification.

The predictive power of the partition information can be measured not only globally but also for each species separately. We could call these measures species’ distinctive ability, and the means of these distinctive abilities can be used as partition goodness. Botta-Dukát et al. (2005) proposed using effect size (Gotelli & McCabe, 2002) calculated from G statistic as a measure of distinctive ability. The sum of effect sizes they called *crispness*. The effect size (i.e., departure from random expectation) is more straightforward than the raw test statistic because G statistics is an increasing function of the number of clusters, even if the partition is random. They applied the theoretical mean and standard deviation deduced from the Chi-square distribution to calculate effect size. Since the calculated G statistic only approximately follows this distribution, rare species, where the approximation is poor, should be excluded from the calculation.

Instead of the G statistic, the Chi-square statistic can also be used. Its maximum is the number of communities. The Chi-square statistic divided by its maximum equals the R^2^ of a linear model where the presence/absence data is the dependent variable, and grouping is the independent variable. Grouping is a categorical variable, but it can be replaced by dummy binary variables that can be treated as continuous predictors. The number of these dummy variables is the number of groups minus one. It is well-known that adding a new variable to the model always increases the value of R^2^, even if it has no relationship with the dependent variables. Therefore, the R^2^ of models with a different number of independent variables cannot be compared. To overcome this problem, Fischer (1925) introduced the adjusted R^2^ that could be used to measure species’ distinctive ability.

From distinctive ability, we can calculate an overall statistic (G statistic or adjusted R-squared). It is useful for evaluating the whole partition but does not inform about the goodness of each group. For this purpose, we can calculate fidelity measures that assess a species’ preference by comparing its frequency (or abundance) in the focal group vs. the rest of the dataset (Bruelheide, 2000). Fidelity measures the goodness of the focal cluster with respect to the occurrence of the focal species. There are two main groups of approaches proposed to summarize this information over species and groups for evaluating the whole partition with respect to all species.

In the first group, the species-by-group matrix of fidelity values is transformed into indicator values of species. The indicator value of each species with respect to the given partition is the highest fidelity over all groups of that partition (Dufrene & Legendre, 1997; Podani & Csányi, 2010). Then, these indicator values are summarized into a measure of the goodness of partition. Dufrene & Legendre (1997) proposed using the sum of significant indicator values. According to Podani and Csányi (2010), the fidelity of all species is worth considering; however, instead of the raw values, their effect sizes (Gotelli & McCabe, 2002) should be summed. Aho et al. (2008) suggested averaging Type I errors instead of indicator values. For representing this group, the last approach was included in the comparison. Each species was characterized by the lowest p-value from Fisher’s exact test over groups, and the mean of these p-values (mean p-values) is used as a goodness measure.

Another group of indices, called OptimClass, is proposed by Tichy et al. (2010). OptimClass first calculates the quality of each group, and then the goodness of partition is the sum of group-level values. In OptimClass1, the cluster’s quality is measured by the number of species with significant positive fidelity (i.e., species with significantly more occurrence in the cluster than expected based on random occurrences across all clusters) according to Fisher’s exact test. The number of significant fidelities depends on the applied significance level. Thus changing the significance level may influence the ranking of partitions. Therefore, it is worthwhile to try several significant levels. In this paper, we applied three significant levels: 10^-3^, 10^-6^, and 10^-9^, abbreviated by OC(3), OC(6), and OC(9), respectively. In the other member of the OptimClass family (OptimClass2), the partitions are characterized by the number of groups with at least *k* significant character species, where *k* is a parameter to be set by the user. The idea behind OptimClass2 is that if there are already enough character species in a group, new ones no longer increase the goodness of the grouping. In other words, it is more important for each group to have a sufficient number of character species than to have many character species in total (which can also be achieved by only one group having many character species). Optimclass1 became more popular than OptimClass2, probably because the latter needs an extra parameter, the number of necessary faithful species. In our experience, it is hard to find the optimal value of this parameter. However, repeated testing with different values may give valuable insight into the classification quality. Since this iterative process cannot be automated, OptimClass2 was not involved in the comparison.

The groups obtained during the classification can be considered as different habitats, and based on this, the niche width of the species can be calculated. If the groups correspond to well-separated habitats, the niche width of the species will be small. The main difference between niche width and measures based on the species’ distinctive ability is that the former does not consider the group sizes. Casado et al. (1997) measured the niche width of the species using the Shannon entropy divided by its possible maximum for the given number of clusters:

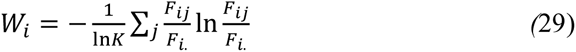

If the species are equally distributed among communities, *W_i_* equals 1, and if a species is present only in one cluster, it equals 0. The cluster validation index is the average of niche widths weighted by the number of occurrences:

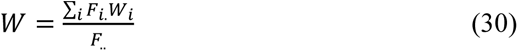

The index ranges between 0 and 1, and a lower value indicates a better partition. It is the “difference-like” criterion. When species *i* occurs, only one group, *W_i_*=0, and further merging will not change its value. Therefore, W tends to decrease with a decreasing number of clusters in a hierarchical classification.

## Results

Figure 1 shows the performance of geometric indices with equal-sized clusters by considering all noise levels and clustering algorithms together. Five CVIs –Tau, Dunn_mean_mean_, and three versions of silhouette width – form the best subset; they were not significantly different from the best partition. These indices performed well irrespectively to noise level and the clustering algorithm. Some indices showed various performances in the three noise levels. The G(+), Dunn_min_max_, and the proportion of correct classification indices were good only at the low noise level. Hubert & Levine, and Feoli performed well in low and medium but not at high levels of noise (Figure S1.1). The clustering algorithm also influences the result of the evaluation. For example, Dunn_min_max_ and the proportion of correct classifications performed well for the classifications created by pam, but their performance was worse when classifications were created by the other two algorithms. Similarly, the two versions of the Popma index behaved according to the expectations only when the classifications were created by the beta-flexible algorithm. Note that differences between the goodness of CVIs were the lowest when classifications were created by the UPGMA algorithm (Figure S1.2). For better understanding differences in performance of indices, it is worthy to see what is the optimal number of clusters (within range from 2 to 20 clusters) according to the different CVIs (Figure 2). CVIs belonging to the best subset correctly indicated ten clusters as optimal solution. Poorly performing indices often selected the partition with the lowest or largest possible number of clusters (i.e., two or twenty clusters) as optimal.

**Figure 1.**
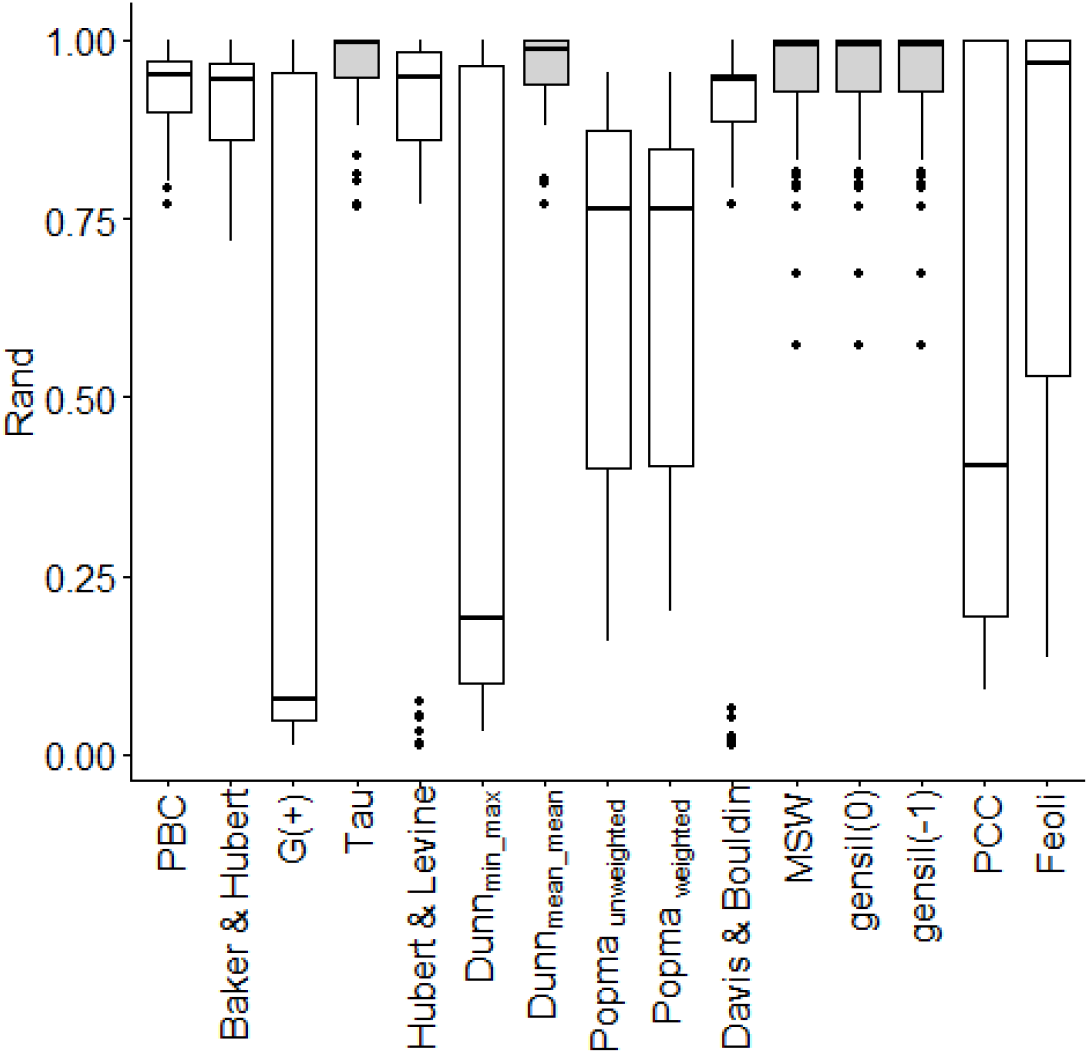
Overall results for the internal geometric indices with equal size clusters (Gray bars indicate indices not significantly different from the best). Abbreviations: PBC = point-biserial correlation; MSW = mean silhouette widths; PCC = proportion of correct classification.

**Figure 2.**
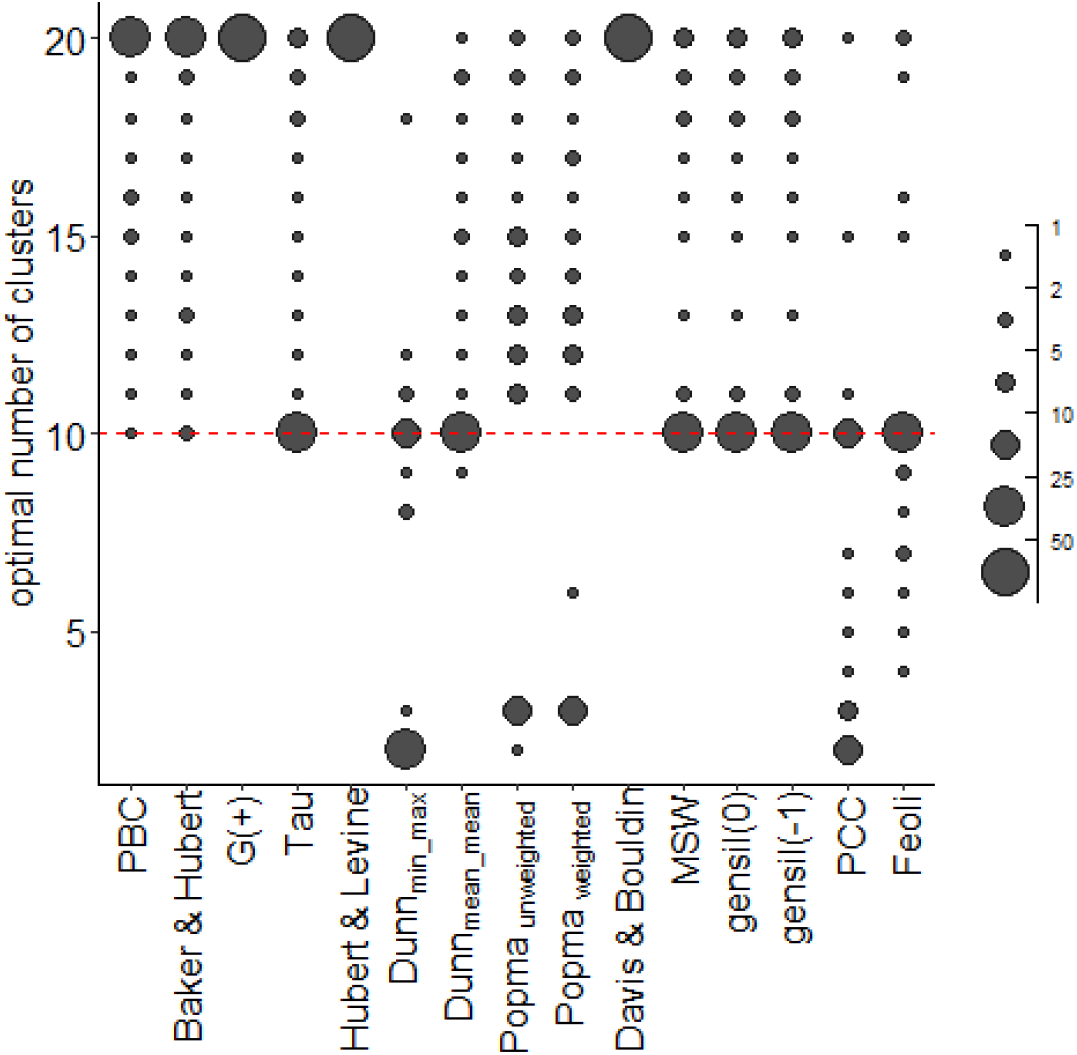
The optimal cluster number selected by the indices with internal geometric indices equal size of clusters (point size indicates the number of times for cluster number, dotted line indicates optimal number of clusters). Abbreviations: PBC = point-biserial correlation; MSW = mean silhouette widths; PCC = proportion of correct classification.

When cluster sizes were unequal, the best subset of CVIs included nine indices (Figure 3). Four of them (Tau and the three versions of silhouette width) were members of the best subset in the previous experiment too. Four CVIs from this best subset – point-biserial correlation, Baker & Hubert, Hubert & Levine, and Davies & Bouldin – also had high goodness in the previous experiment with equal cluster sizes. However, they did not belong to the best subset there. More noteworthy is the difference in the behavior of Dunn_mean_mean_ and the proportion of correct classification indices. Dunn_mean_mean_ was among the best when cluster sizes were equal, but it had only medium goodness in the second experiment with unequal cluster sizes. The proportion of correct classification had a high variation between replications in both experiments. However, its median performance was high when cluster sizes were unequal. G(+) and the proportion of correct classification indices proved to be somewhat sensitive to the level of noise (Figure S2.1). Differences in goodness between clustering algorithms were not consistent with the results of the experiment with equal group sizes. For example, two versions of Popma were among the best ones for beta flexible classification when cluster sizes were equal, and they seemed to be unsuitable only for this algorithm when the cluster sizes were unequal (Figure S2.2).

**Figure 3.**
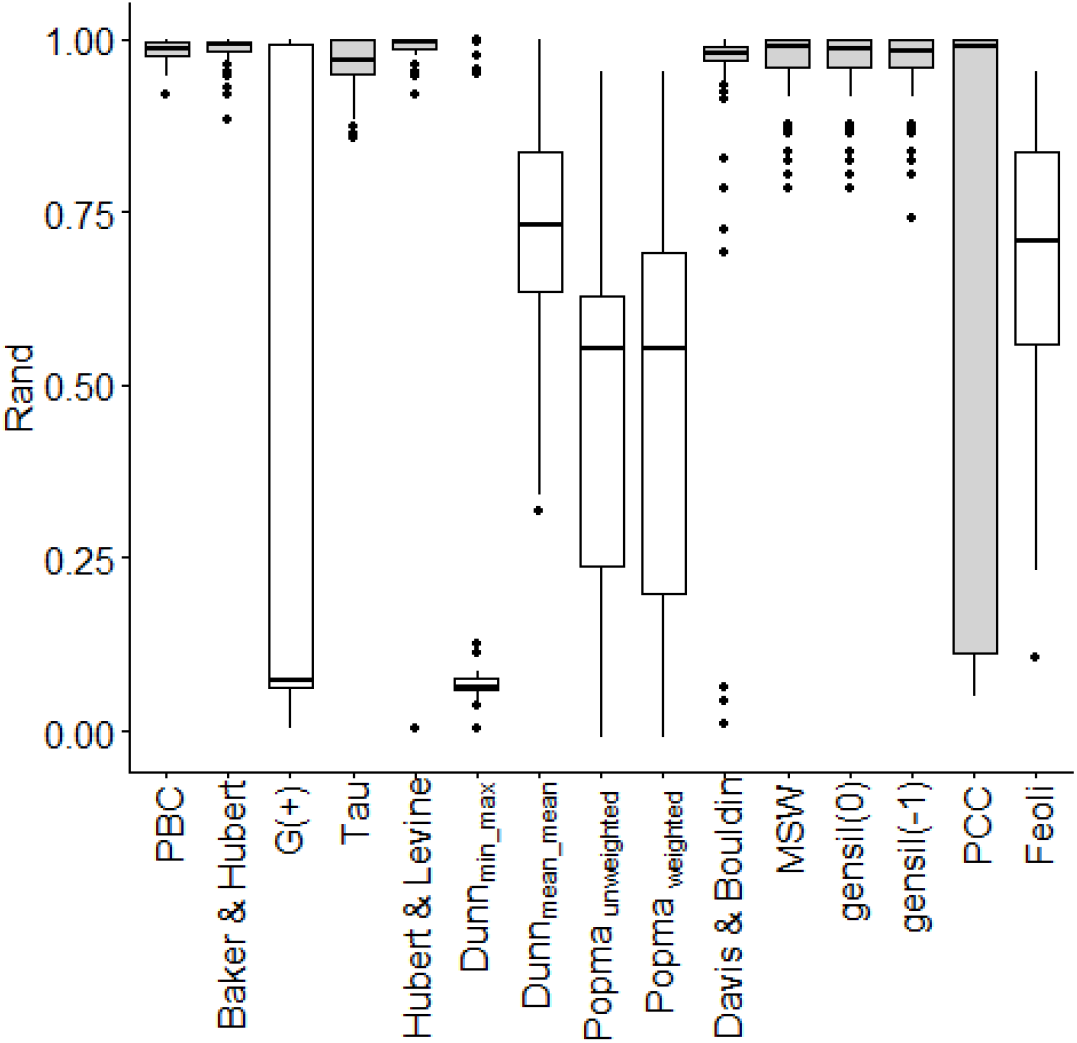
Overall results for the internal geometric indices with unequal size clusters (Gray bars indicate indices not significantly different from the best). Abbreviations: PBC = point-biserial correlation; MSW = mean silhouette widths; PCC = proportion of correct classification.

**Figure 4.**
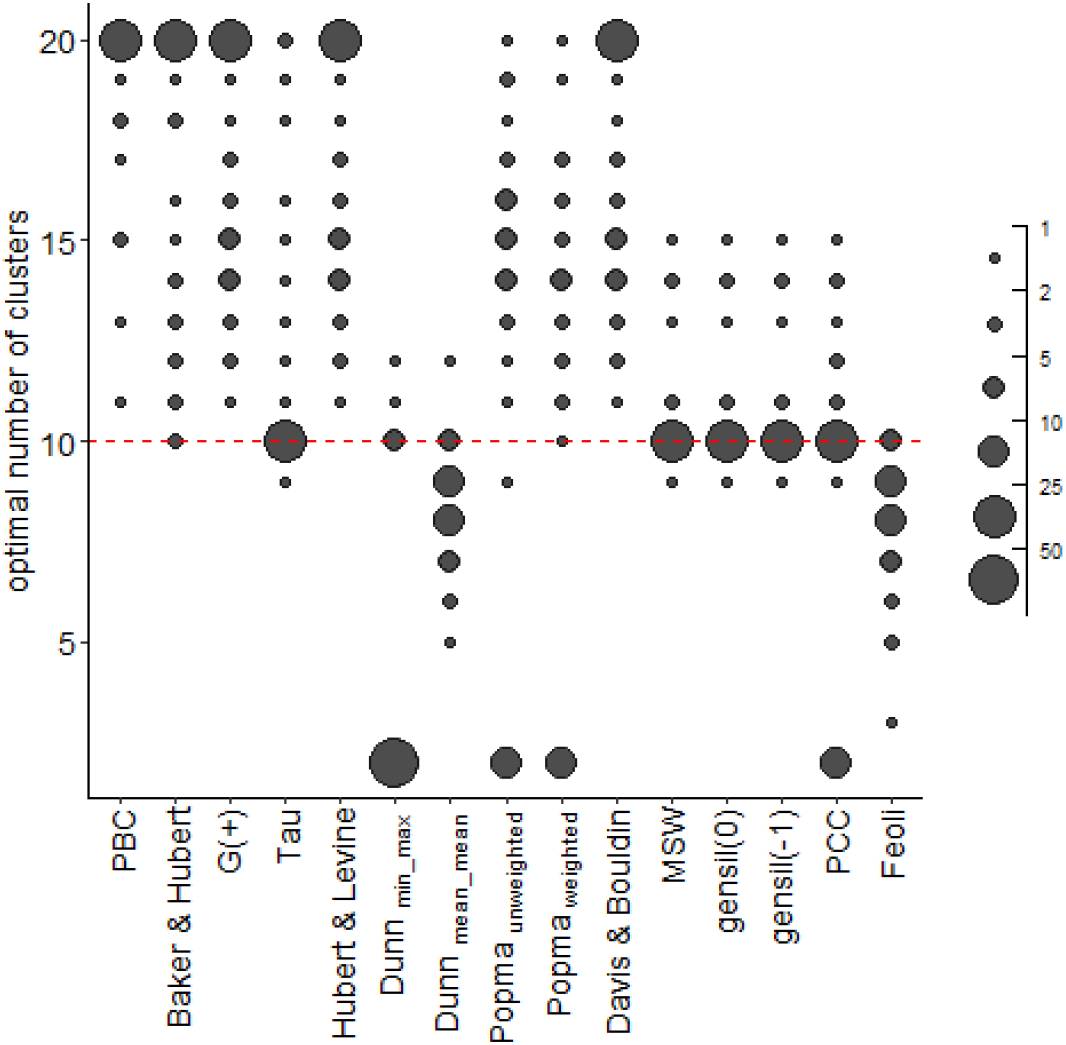
The optimal cluster number selected by the indices with internal geometric indices unequal size of clusters (point size indicates the number of times for cluster number, dotted line indicates optimal number of clusters) Abbreviations: PBC = point-biserial correlation; MSW = mean silhouette widths; PCC = proportion of correct classification.

In the case of equal-sized clusters, Cripness and OptimClass formed the best subset of non-geometric indices in the overall evaluation (Figure 5). The noise level strongly influenced the results: at low noise level, relative divergence, crispness, and mean p-values formed the best subset, while at high noise level, mean p-values and OptimClass (Figure S3.1). Using UPGMA and pam algorithms, there was no significant difference in performance between indices. Only the flexible beta algorithm creates cluster structures in which the performance of indices differs (Figure S3.2). Sometimes even the well-performing indices overestimated the optimal number of clusters (Figure 6). The overall performance of non-geometric indices was similar in the experiment with unequal-sized clusters (Figure 7, 8, S4.1, S4.2), where mean p-values also belonged to the best subset.

**Figure 5.**
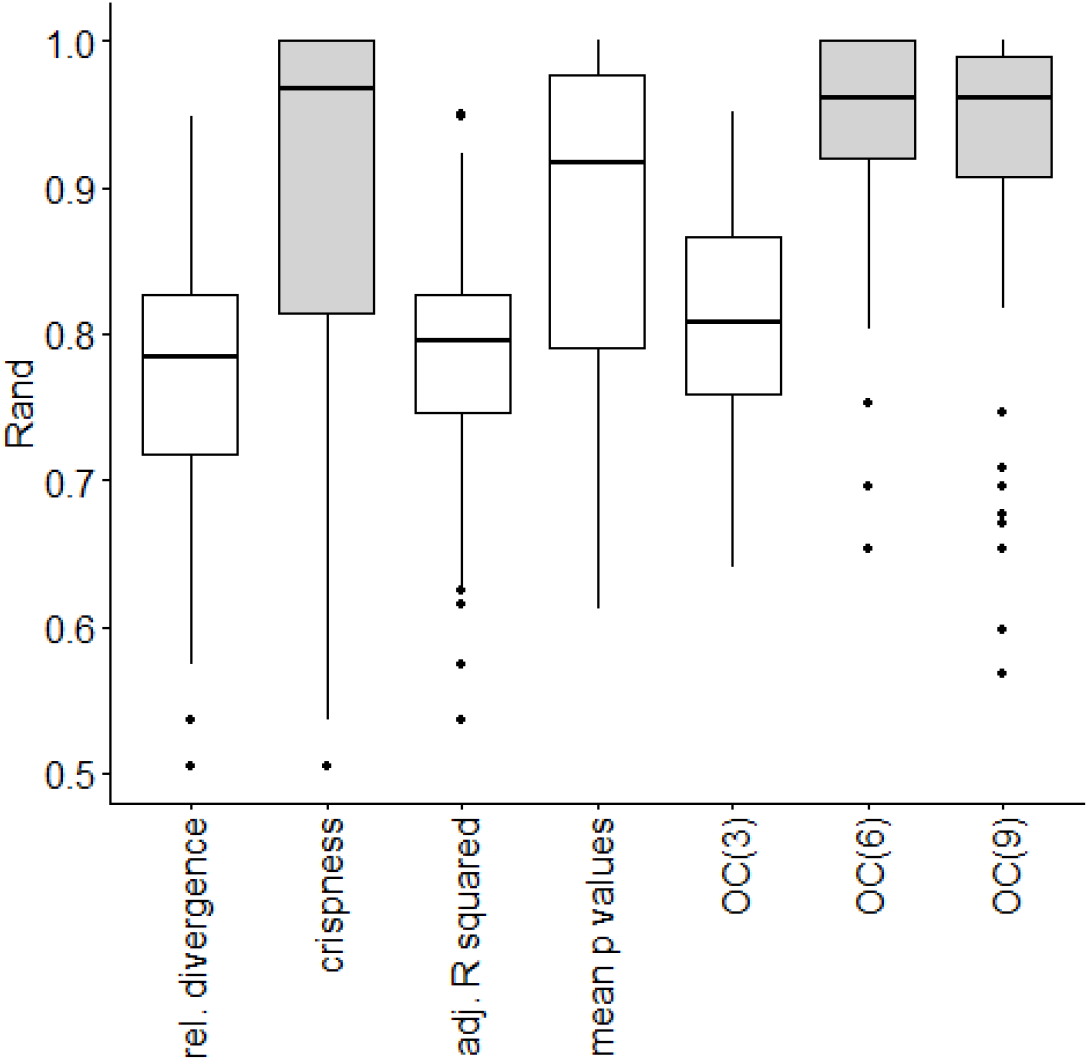
Overall results for internal non-geometric indices with equal size clusters (Gray bars indicate indices not significantly different from the best). Abbreviations: OC(x) = OptimClass with significance level 10^-x^

**Figure 6.**
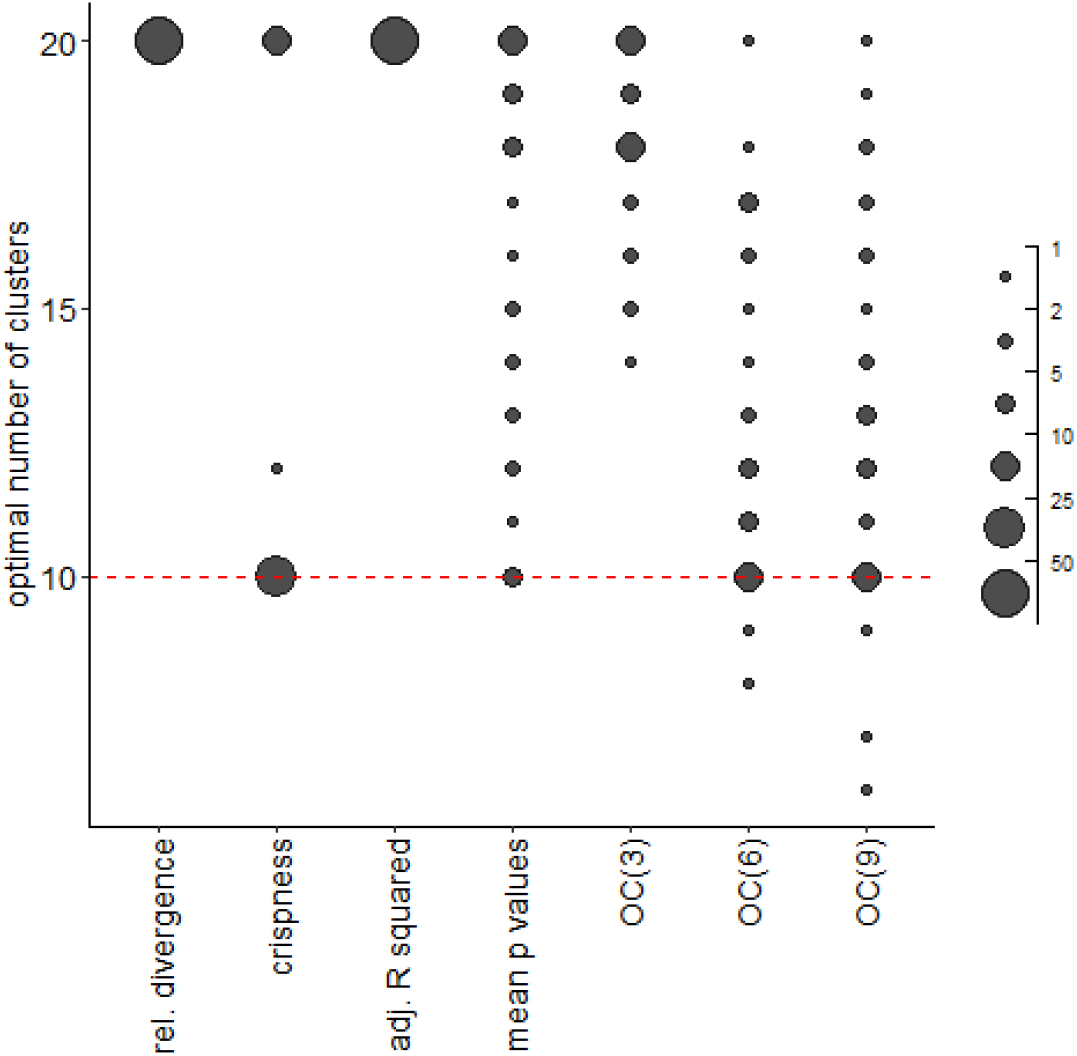
The optimal cluster number selected by the indices with internal non-geometric indices equal size of clusters (point size indicates the number of times for cluster number, dotted line indicates optimal number of clusters). Abbreviations: OC(x) = OptimClass with significance level 10^-x^

**Figure 7.**
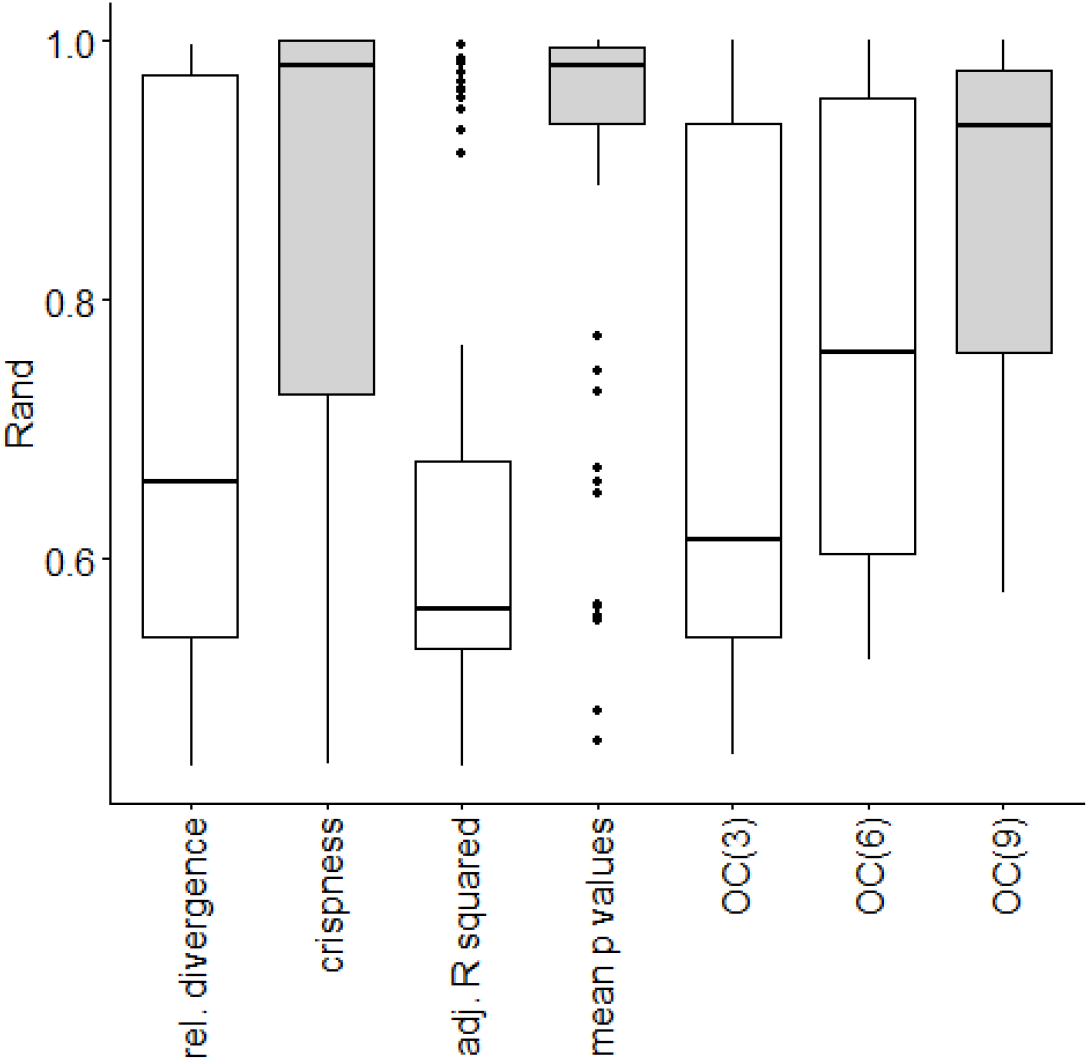
Overall results for internal non-geometric indices with unequal size clusters (Gray bars indicate indices not significantly different from the best). Abbreviations: OC(x) = OptimClass with significance level 10^-x^

**Figure 8.**
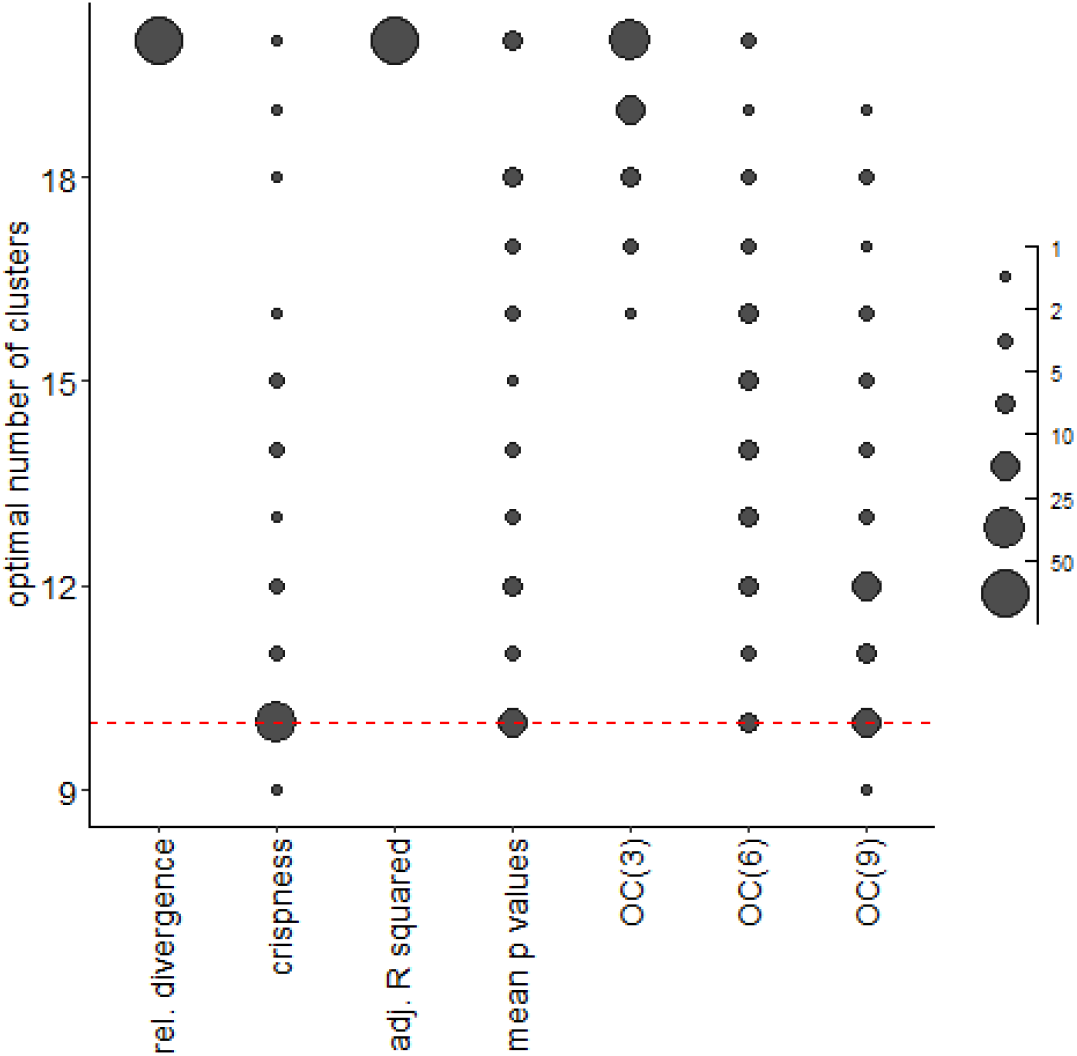
The optimal cluster number selected by the indices with internal non-geometric indices unequal size of clusters (point size indicates the number of times for cluster number, dotted line indicates optimal number of clusters). Abbreviations: OC(x) = OptimClass with significance level 10^-x^

## Discussion

Our results show that *Tau* and *silhouette widths* were the best geometric CVIs both for equal and unequal cluster sizes. In line with our study, *mean silhouette width* had shown robust performance in previous comparisons (Arbelaitz et al., 2013; Niemelä et al., 2018; 2010), including classification tasks complicated due to high dimensionalities, noise, overlapping clusters or missing data. Our results emphasize the usefulness of the silhouette indices, due to the fact that they are unaffected by noise levels or clustering algorithms. The silhouette width combines two clustering criteria, compactness, and separation (Rousseeuw, 1987). Mean silhouette width tends to prefer spherical clusters; the mean generalized silhouette indices avoid this limit (Lengyel & Botta-Dukát, 2019). In our experiments, there were no considerable differences in performance between different versions of silhouette width; however, we tested only a narrow range of parameter values (*q*=-1, *q* = 0 and *q*=1). However, we cannot rule out that the similar performance by silhouette indices was a consequence of the algorithm for generating data that created spherical clusters. Elongated clusters are probably rarer when binary variables are analyzed, since in this case all variables have the same range, contrary to continuous variables. More detailed research is needed to explore if and when different parameterizations of generalized silhouette width outperform the original definition.

Tau is closely related to point-biserial correlation; the only difference is that Tau is a rank correlation, while point-biserial is a linear correlation. In our evaluation, both reached high goodness, but the point-biserial correlation overestimated the optimal number of clusters. Vendramin et al. (2010) also observed that the number of clusters in the best partition according to point-biserial correlation often differs from the real number of clusters, but in their experiment, Tau proved to be even worse in this respect. In the study of Milligan and Cooper (1985), point-biserial correlation belonged to the best indices; however, its performance slightly worsened with the increasing number of clusters. In that study, Tau often overestimated the number of clusters. On the other hand, Vendramin et al. (2010) have found strong correlations with external indices for both point-biserial correlation and Tau. They concluded that these CVIs are suitable for choosing the optimal algorithm but unsuitable for selecting the optimal number of clusters. Inspecting the changes of the two indices with an increasing number of clusters (see figures in Appendices), it can be seen that both Tau and point-biserial correlation steeply increased until reaching the true number of clusters. Over the true number of clusters, Tau slightly decreased, thus often having a maximum at the true number of clusters. Point-biserial correlation did not decrease or even slightly increase, leading to overestimating the number of clusters. It also suggests that for choosing the optimal number of clusters, the plot of CVI values against the number of groups should be visually evaluated instead of automatically selecting the position of maximum as the optimal number. It helps to recognize situations when the decision based on the applied CVI is doubtful; thus, involving other CVI is necessary. Using departure from the random expectation (e.g., Measure of Relative Improvement; Vargha et al., 2016) instead of raw CVI values may also help in such cases. Finally, a possible drawback of Tau compared to point-biserial correlation is its much longer runtime (Aschenbruck & Szepannek, 2020).

Another CVI based on rank correlation is the Baker & Hubert index, but contrary to Tau, it does not consider the number of ties. In previous studies (Milligan & Cooper, 1985; Vendramin et al., 2010), this index proved to be good for selecting the true cluster number, but its values were only weakly correlated with the external validity measures. In our experiments, its goodness was high, but it tended to choose the highest possible number of clusters as optimal. Note that ties are more frequent in distances calculated from binary than abundance data. This fact could explain the better performance of Tau in our experiment.

Several CVIs that combine compactness and separation of groups reached high goodness, but they, except for silhouette width, often overestimated the optimal number of clusters. Thus, they may be useful for choosing the optimal clustering algorithm rather than deciding the number of clusters. The reason for the overestimation is that their values steeply change with the number of clusters below the true value but only slightly change above that, where the fluctuation may override this weak trend.

Two studied versions of the Dunn index considerably differed in their goodness. Our results confirmed that Dunn’s original index (Dunn_min_max_) is unsuitable for cluster validation because of its high sensitivity to noise (Davies & Bouldin, 1979). But it provides a rich and very general structure for defining cluster validity indexes for different types of clusters with suitable distances and set diameter generalizations (Bezdek & Pal, 1998). Dunn_mean_mean_ index behaved well, especially with low noise and equal cluster size conditions. It would be worth investigating which definition of separation and compactness could result in the best Dunn-type index for binary data.

Geometric cluster validation indices can be used for choosing the optimal algorithm, the optimal number of clusters, or both. If they are used for selecting the optimal cluster number, they could be called “stopping rules” (Milligan & Cooper, 1985). Different stopping rules could be the best for different clustering algorithms. Although we have found some differences between algorithms in CVIs’ behavior, these differences are not robust enough to draw any general conclusion.

Among non-geometric indices, Crispness and OptimClass proved to be the best. Note that Crispness applies an approximation that is valid only for frequent species. Therefore, it was suggested to exclude rare species from the calculation. In our simulated datasets, species’ frequencies were the same; thus, all species were used for calculating crispness. Excluding a lot of rare species may distort the results of Crispness, especially when these species are all concentrated in the same cluster. OptimClass is based on an exact test; thus, it is less sensitive to the presence of rare species. OptimClass has a parameter (the significance level) that could be changed, and this setting influences the performance of this validity measure. There is no generally applicable significance level that would be suitable for all datasets; hence, Tichy et al. (2010) suggested trying several different parameterizations. At first glance, it seems to be a disadvantage due to the lack of a universal outcome, but if there is a hierarchical group structure (e.g., communities can be grouped into associations and associations into alliances), applying different significance levels can reveal this hierarchy instead of choosing only one level as optimal.

## Conclusions

Although, in theory, all CVIs could differentiate between good and wrong classifications, only a few of them perform as expected with noisy data. Since the level of noise is not known in real situations, we recommend using four CVIs that are proved to be most robust against noise: Tau, (generalized) mean silhouette width, crispness, and OptimClass. The first two CVIs are geometric indices. Therefore, they are unsuitable for comparing classifications obtained using different dissimilarities.

We suggest plotting CVI against the number of clusters because the lack of a sharp peak means that the position of the maximum is uncertain. Moreover, the presence of local peaks may indicate a hierarchy of groups. For example, in Botta-Dukát et al. (2005), the first peak of crispness (at three clusters) indicated syntaxonomical alliances, while the third peak (at nine clusters) corresponded to associations.

We strongly recommend using several CVIs and different parameters of generalized silhouette width or OptimClass. Congruent results of different CVIs give stronger evidence for selecting the optimal classification than a decision based on a single CVI. Nevertheless, we should keep in mind that alternative classifications of the same dataset may be similarly meaningful, and in this case, different CVIs may choose different partitions as the best.

CVIs included in this comparison are sensitive to preserving distance information (e.g., Tau), or compactness and separation of groups (e.g., mean silhouette width), or interpretability of clusters (e.g., OptimClass). Other aspects of classification goodness not tested here – e.g., stability (Tichý et al., 2011), robustness (Datta & Datta, 2003), or repetitiveness (Breckenridge, 1989; Illyés et al., 2007) – may be important as well; thus, they should not be neglected when selecting CVIs for real case studies.

