## Appendix 3 for "Quantitative evaluation of internal clustering validation indices using binary datasets"

**Appendix 3.** Comparison of performance of the internal non-geometric indices for equal cluster sizes

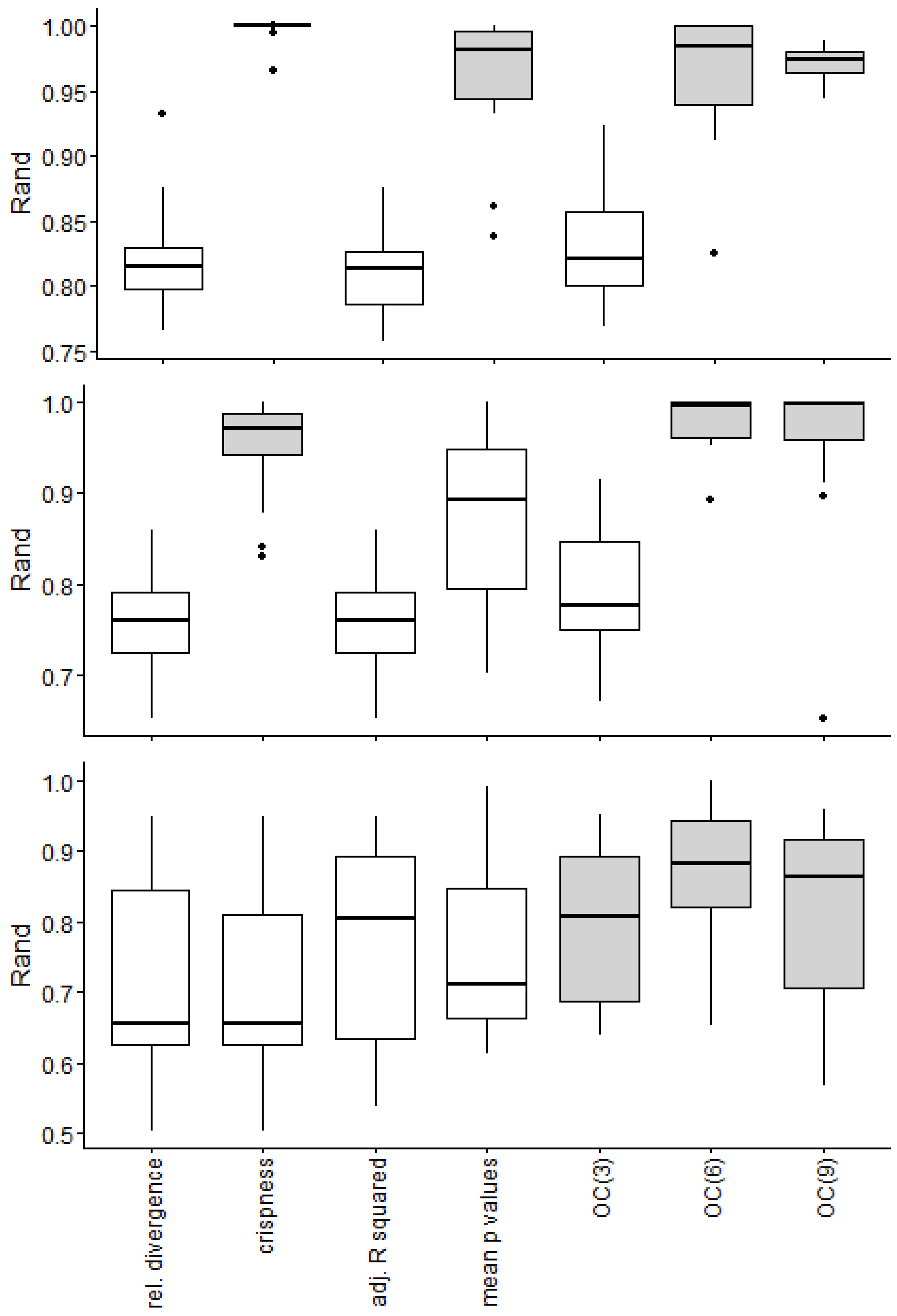


**Figure S3.1** Performance of different geometric indices for each noise level (upper = low noise, bottom=high noise) (Gray bars indicate indices not significantly different from the best). Abbreviations: OC(x) = OptimClass with significance level $10^{-x}$

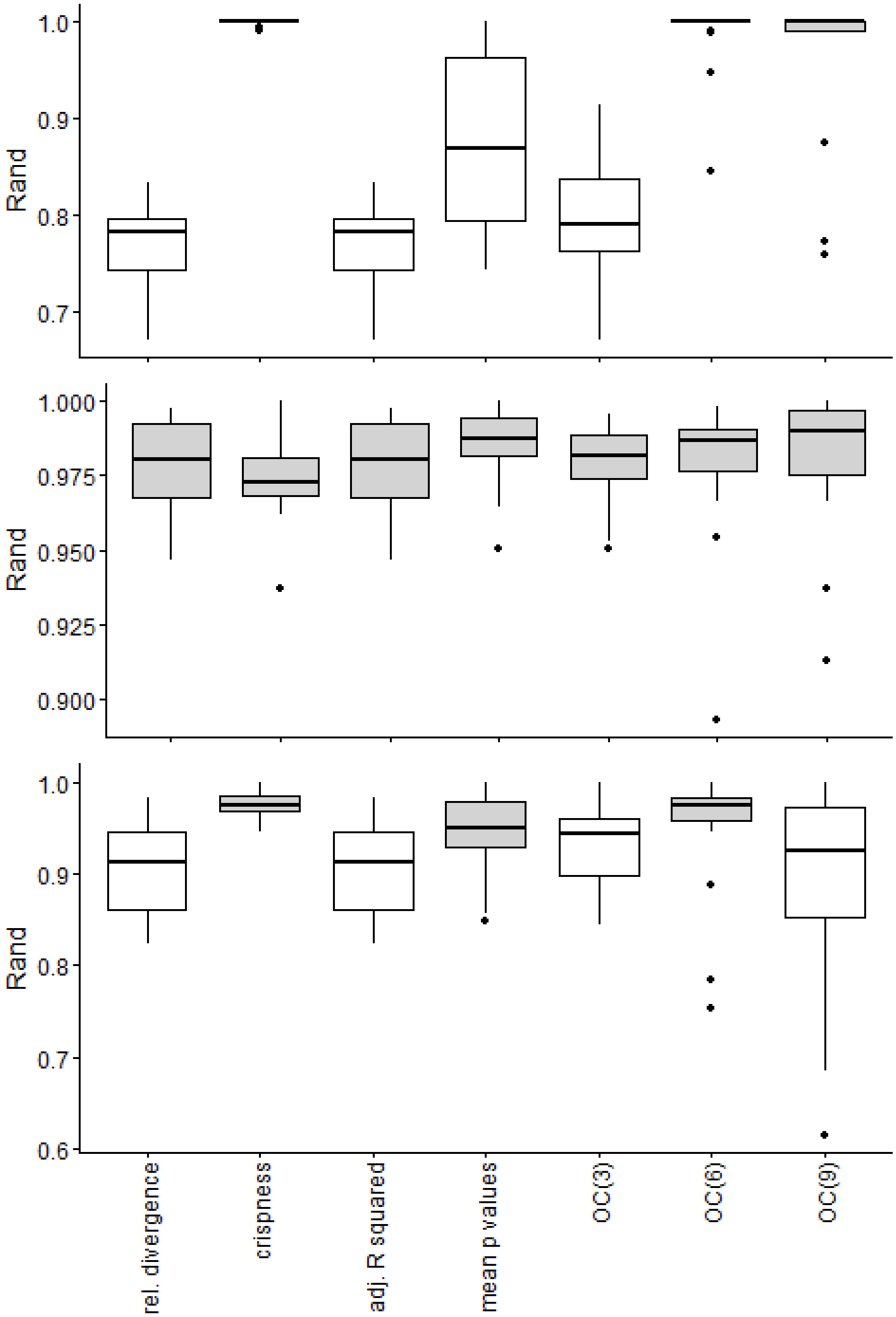


**Figure S3.2** Performance of each clustering algorithm separately at the medium level of noise (upper=pam, medium = UPGMA, lower = flexible). (Gray bars indicate indices not significantly different from the best). Abbreviations: OC(x) = OptimClass with significance level $10^{-x}$

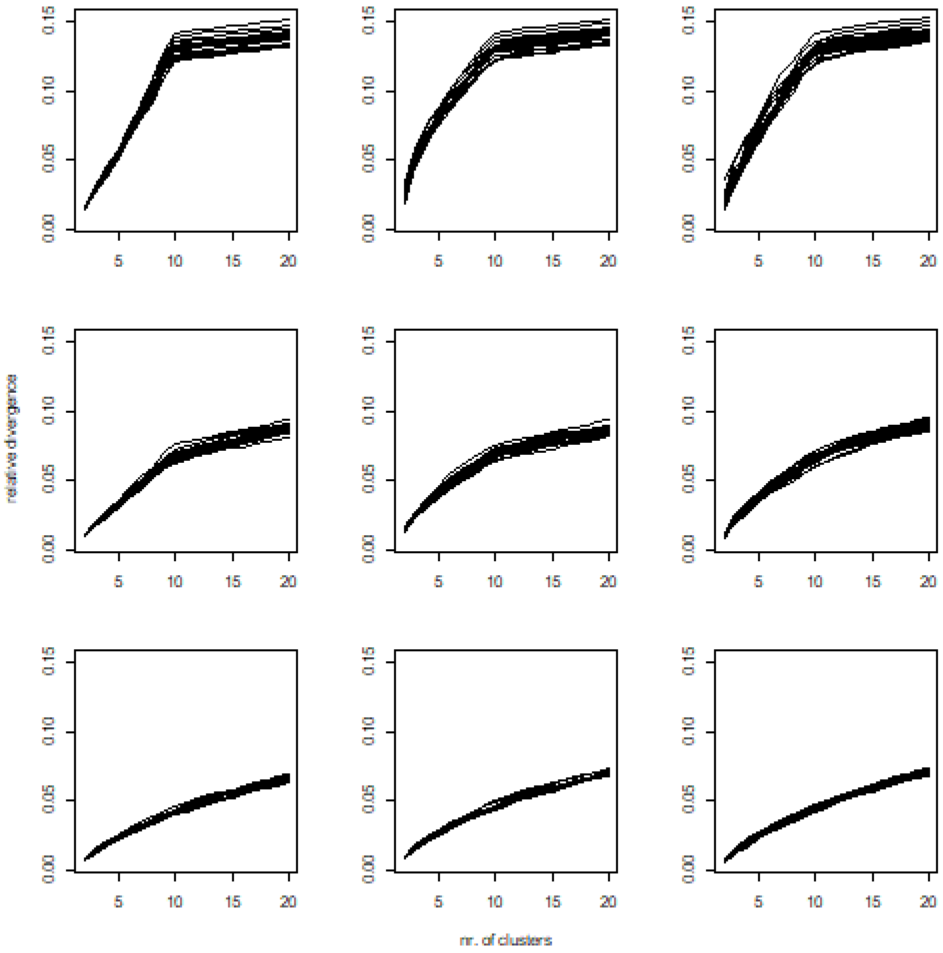


**Figure S4.3**. Relative divergence index versus the number of clusters. Noise levels are in rows (upper is the low noise), and algorithms are in columns (left = pam, medium = UPGMA, right = flexible).

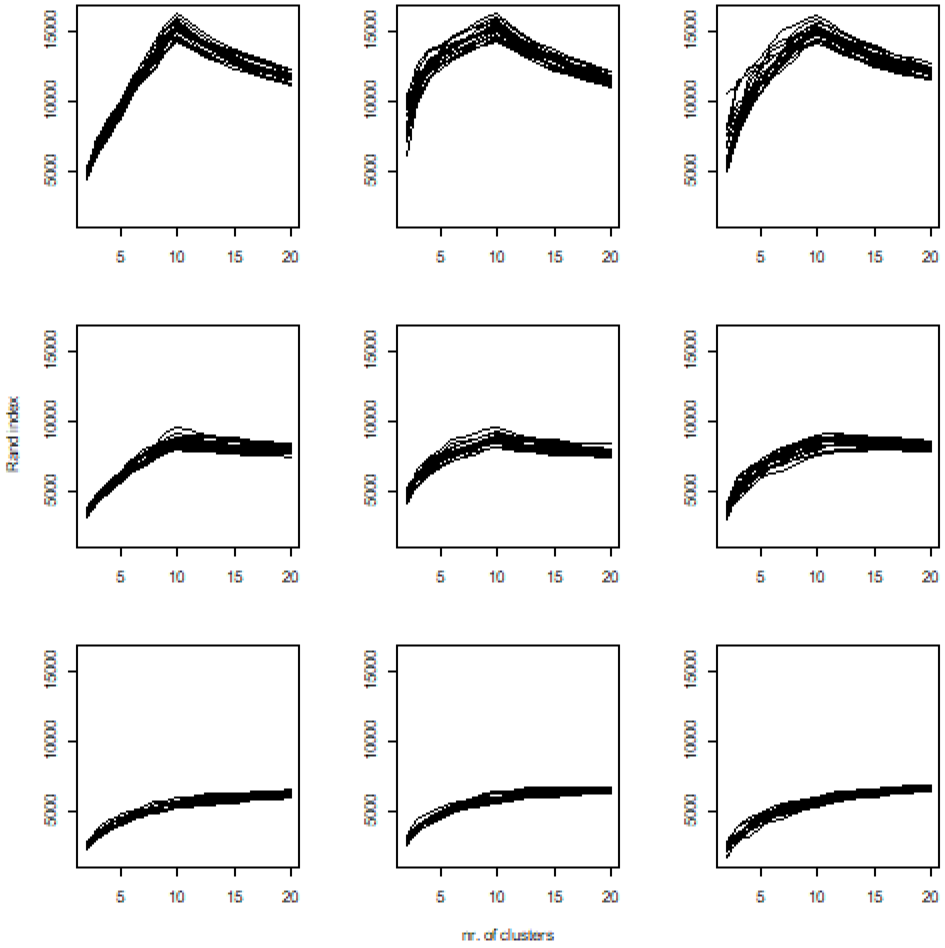


**Figure S4.4**. Crispness index versus the number of clusters. Noise levels are in rows (upper is the low noise), and algorithms are in columns (left = pam, medium = UPGMA, right = flexible).

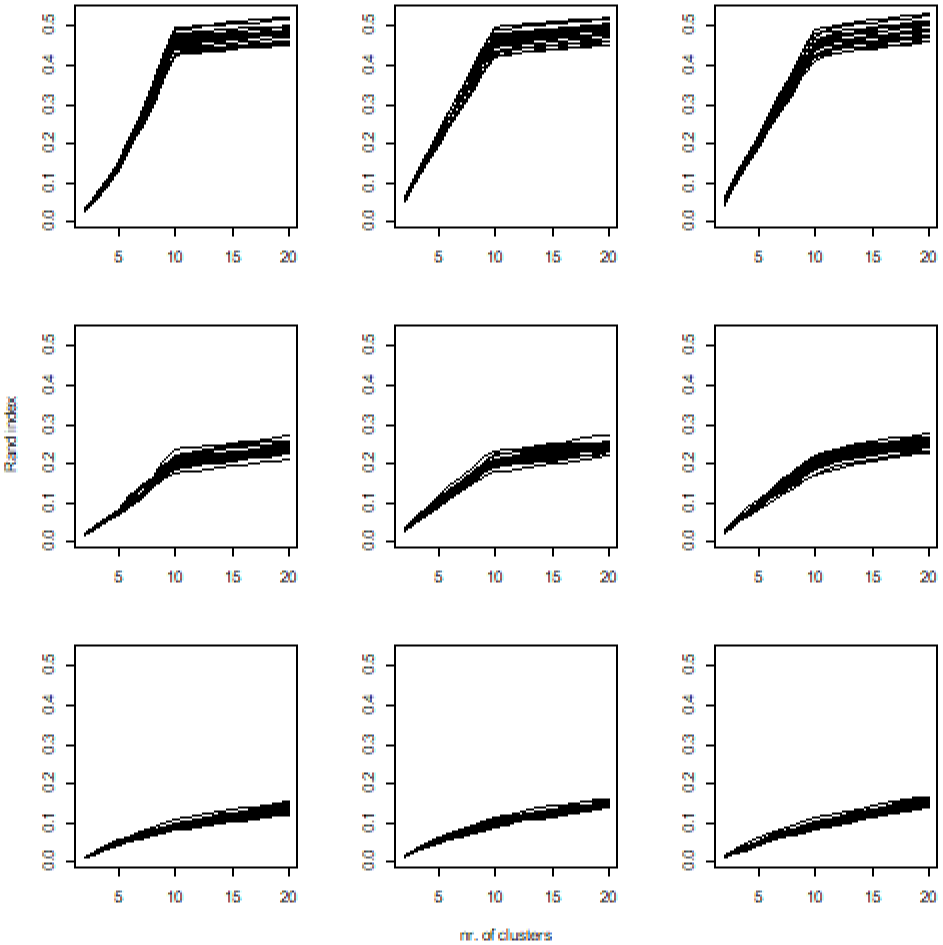


**Figure S4.5.** Adjusted R squared versus the number of clusters. Noise levels are in rows (upper is the low noise), and algorithms are in columns (left = pam, medium = UPGMA, right = flexible).

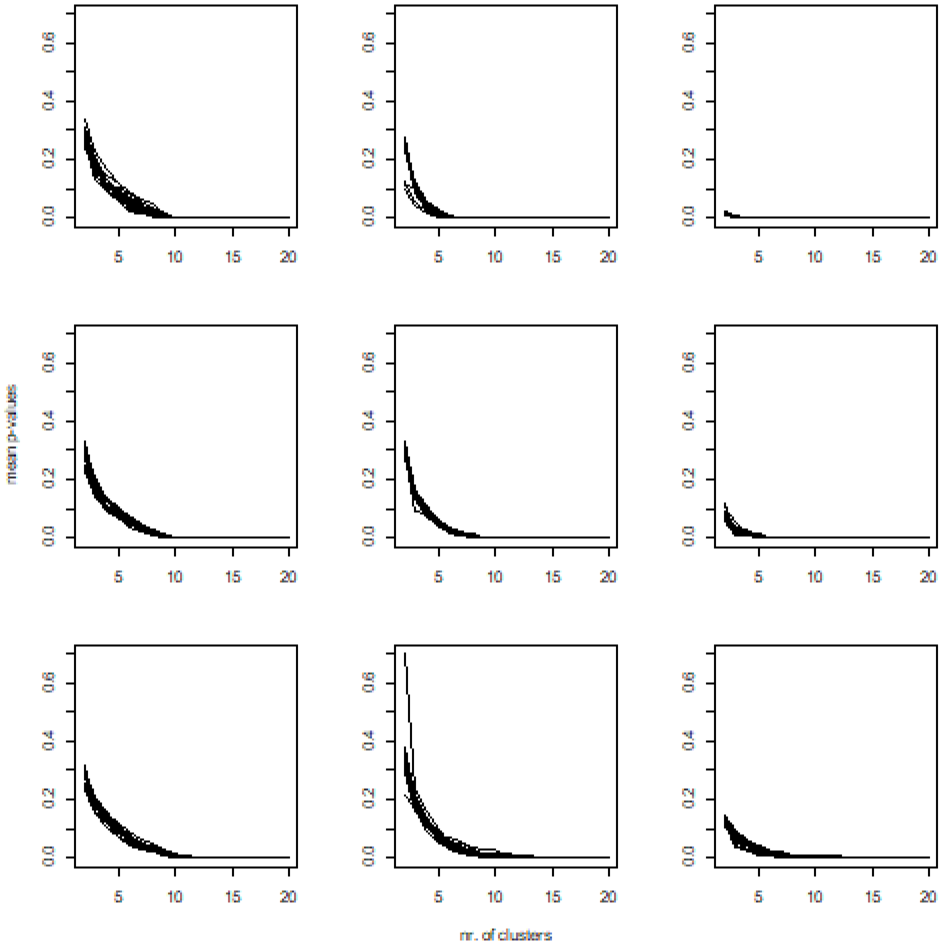


**Figure S4.6**. Mean p values index versus the number of clusters. Noise levels are in rows (upper is the low noise), and algorithms are in columns (left = pam, medium = UPGMA, right = flexible).

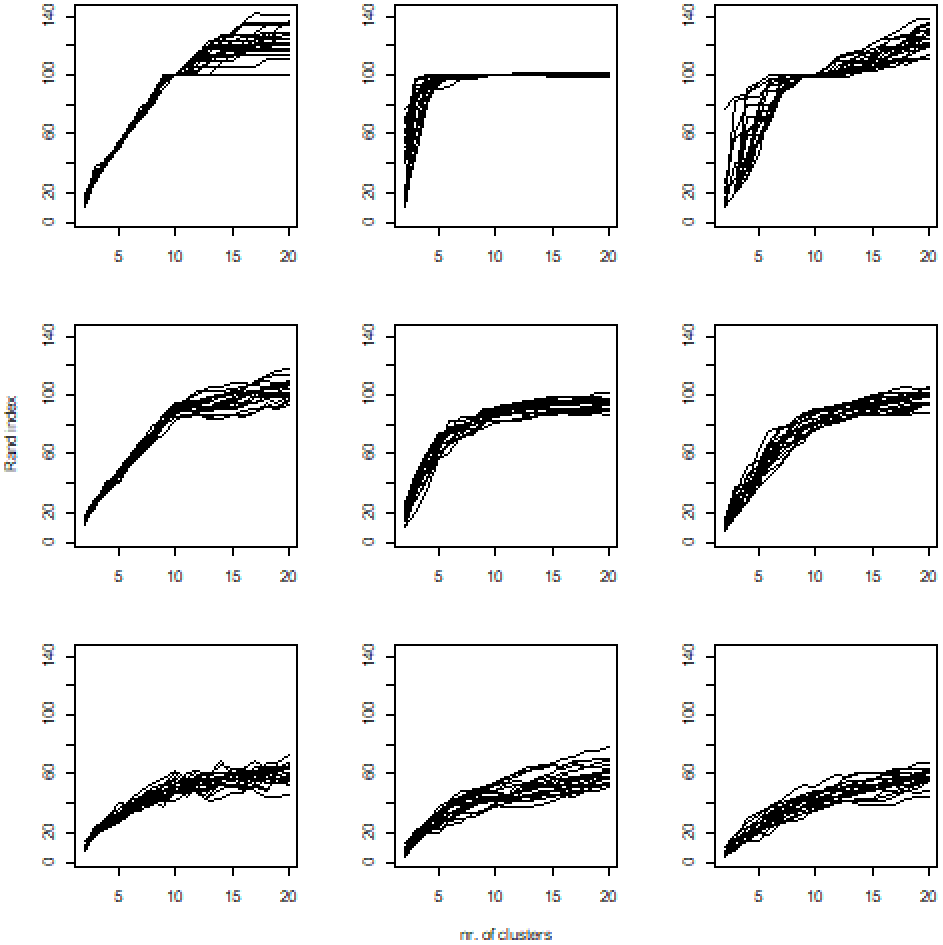


**Figure S4.7.** OptimClass p=0.001 index versus the number of clusters. Noise levels are in rows (upper is the low noise), and algorithms are in columns (left = pam, medium = UPGMA, right = flexible).

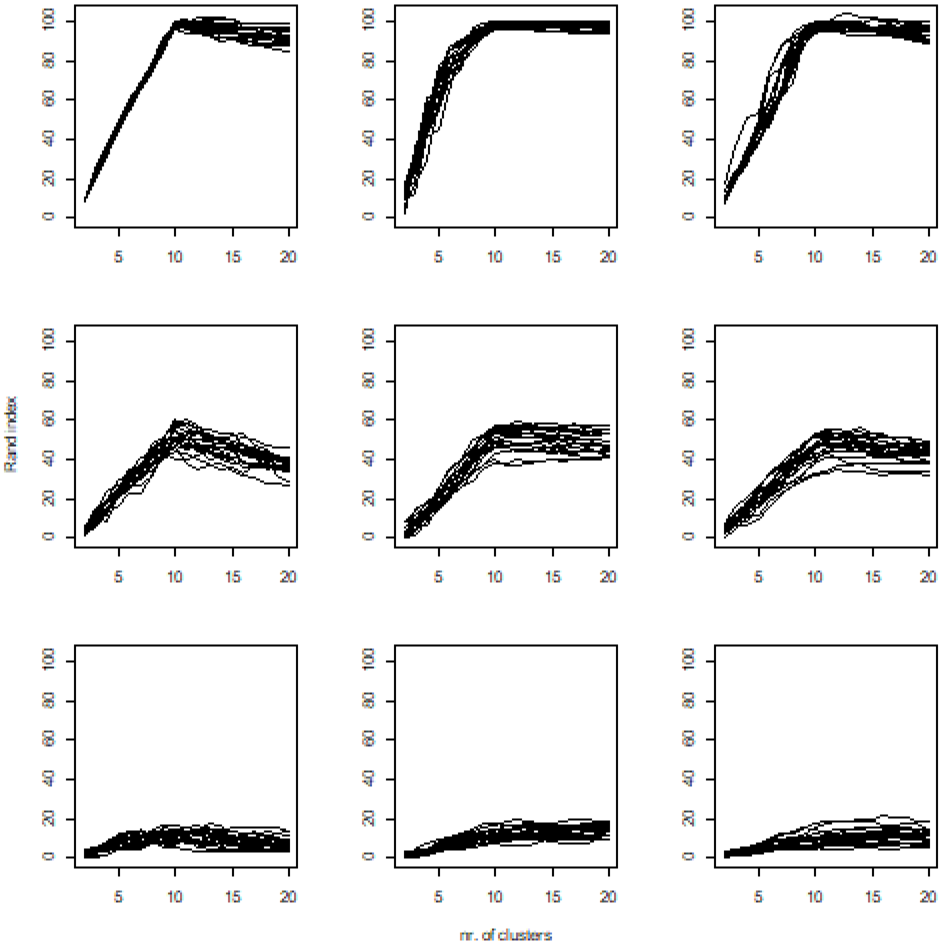


**Figure S3.8.** OptimClass p=1e_6 index versus the number of clusters at different noise levels. Noise levels are in rows (upper is the low noise), and algorithms are in columns (left = pam, medium = UPGMA, right = flexible).

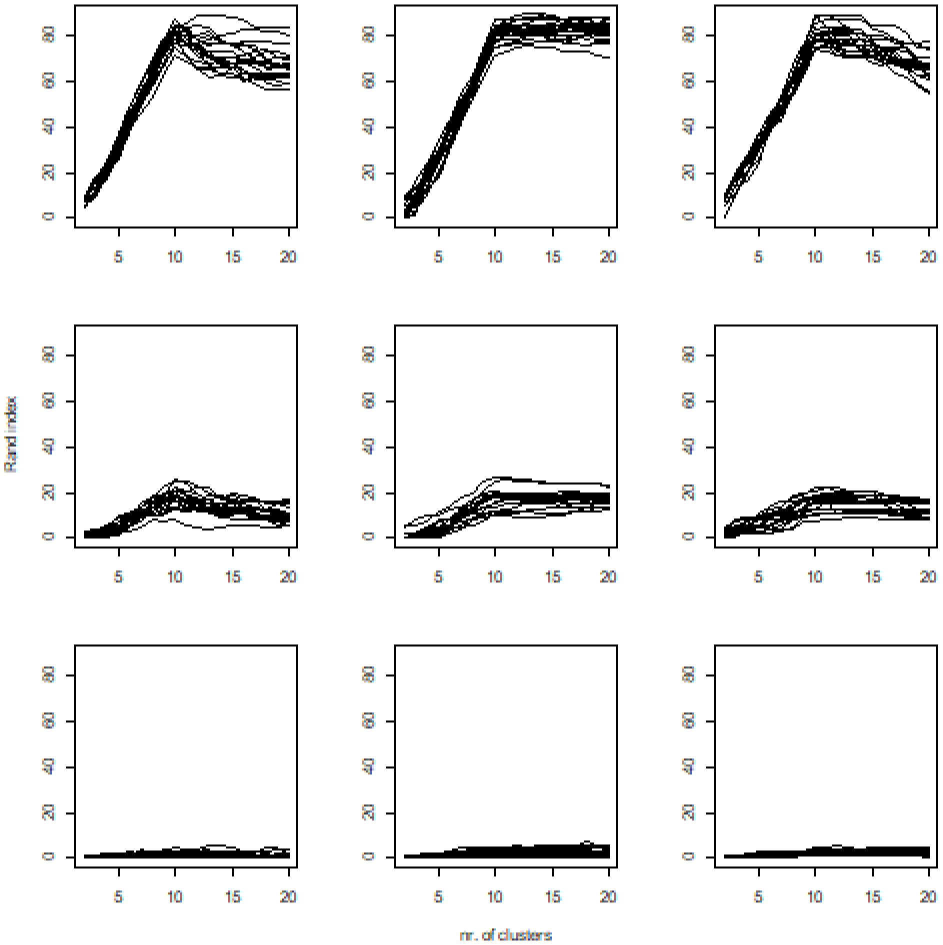


**Figure S3.9.** OptimClass p=1e_9 index versus the number of clusters at different noise levels. Noise levels are in rows (upper is the low noise), and algorithms are in columns (left = pam, medium = UPGMA, right = flexible).
