## Appendix 2 for "Quantitative evaluation of internal clustering validation indices using binary datasets"

**Appendix 2.** Comparison of performance of the internal geometric indices for unequal cluster sizes

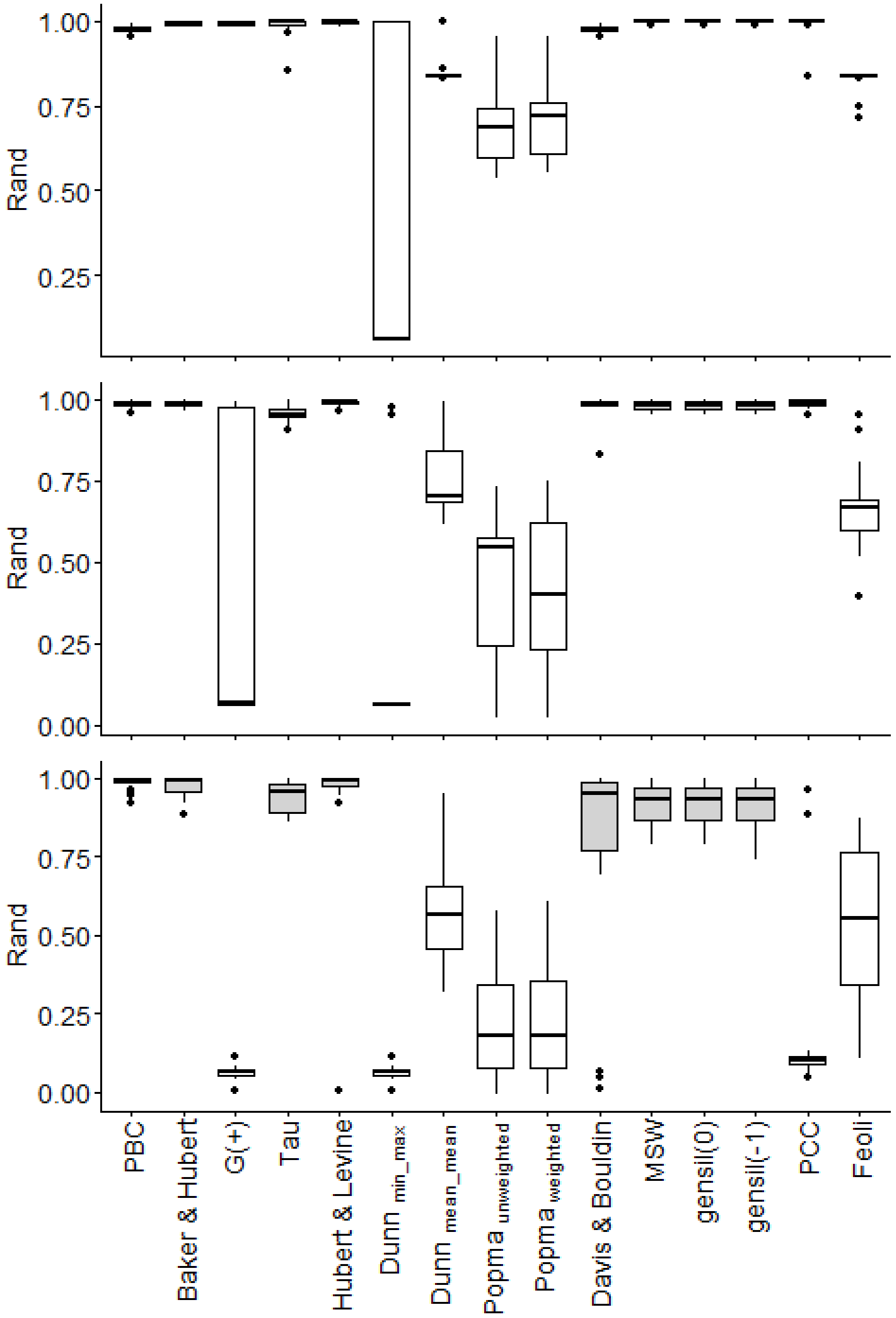


**Figure S2.1**-Performance of different geometric indices for each noise level (upper = low noise, bottom=high noise) (Gray bars indicate indices not significantly different from the best) Abbreviations: PBC = point-biserial correlation; MSW = mean silhouette widths; PCC = proportion of correct classification.

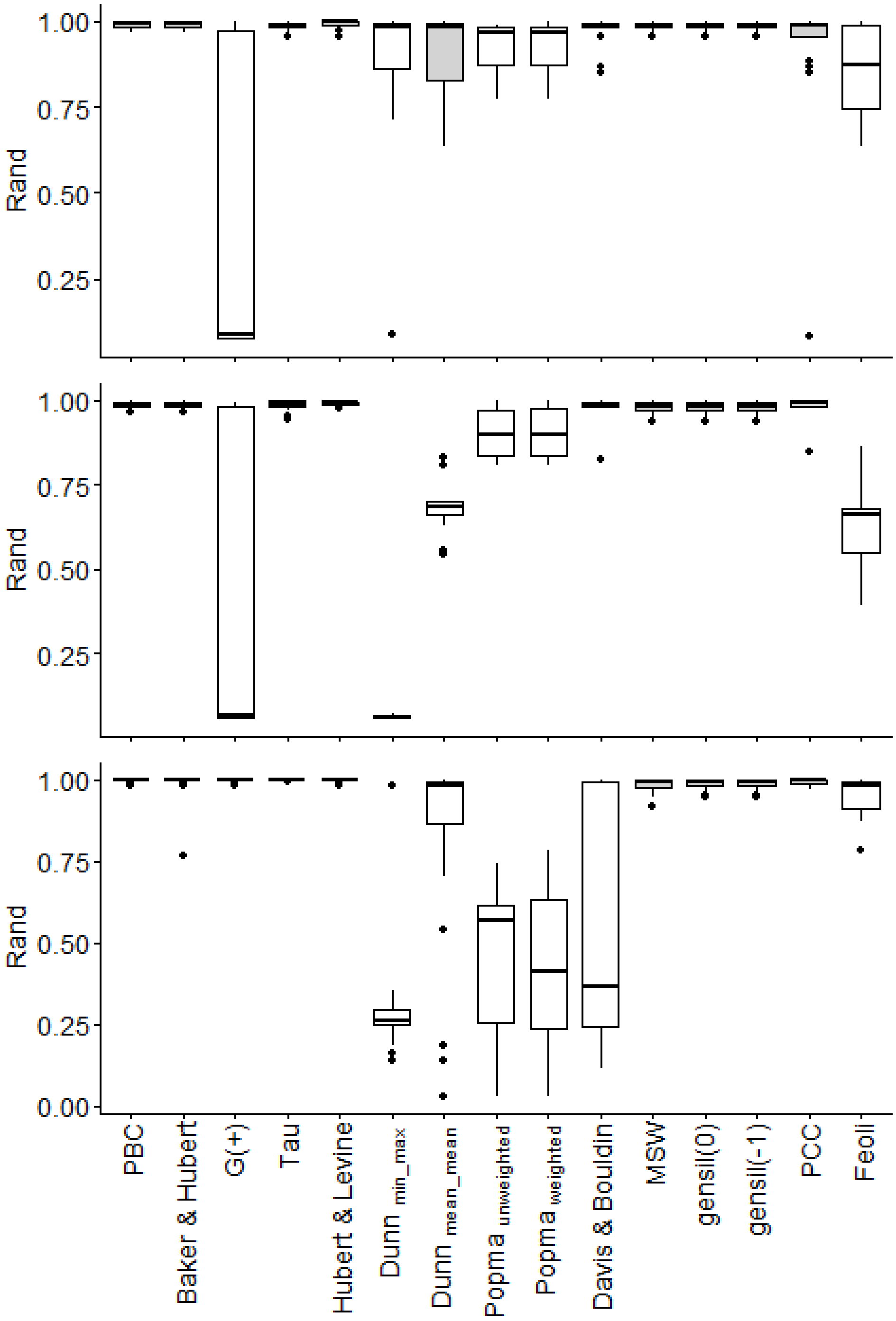


**Figure S1.2-**Performance of each clustering algorithm separately at the medium level of noise (upper=pam, medium = UPGMA, lower = flexible). (Gray bars indicate indices not significantly different from the best) Abbreviations: PBC = point-biserial correlation; MSW = mean silhouette widths; PCC = proportion of correct classification.

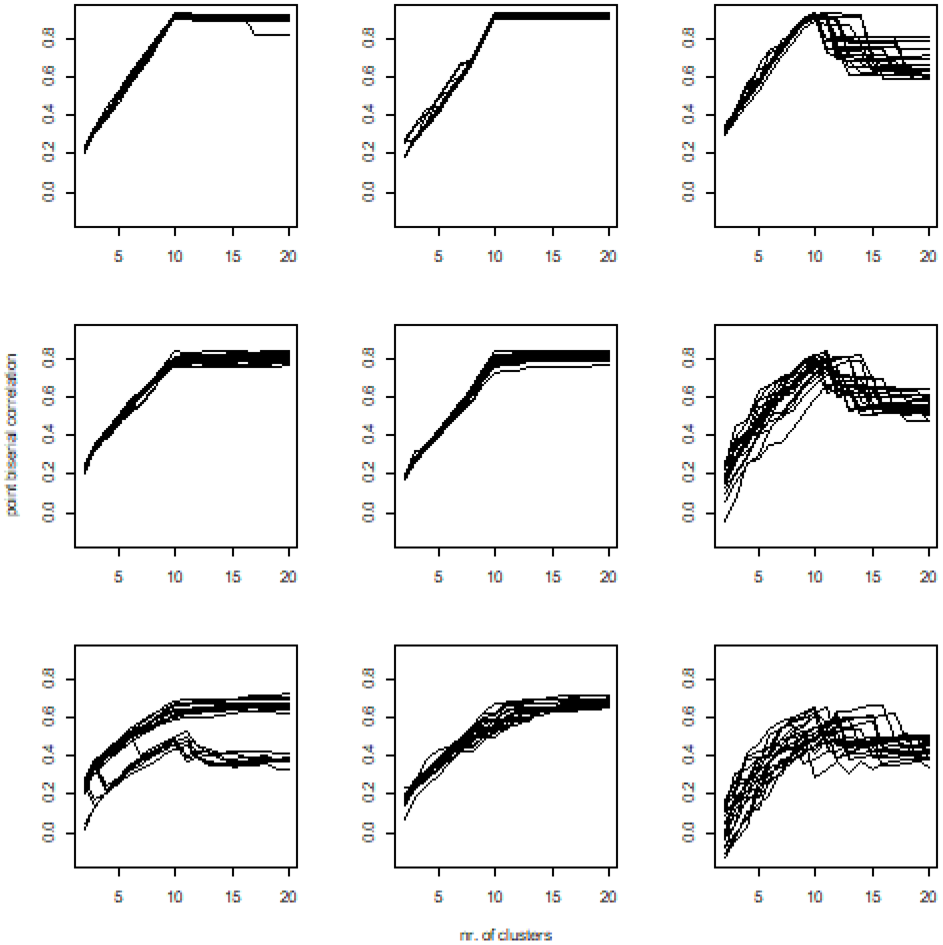


**Figure S2.3** Performance of point. biserial index versus the number of clusters at different noise levels. Rows indicate noise level (upper is the low noise), and columns are algorithms (left = pam, medium = UPGMA, right = flexible).

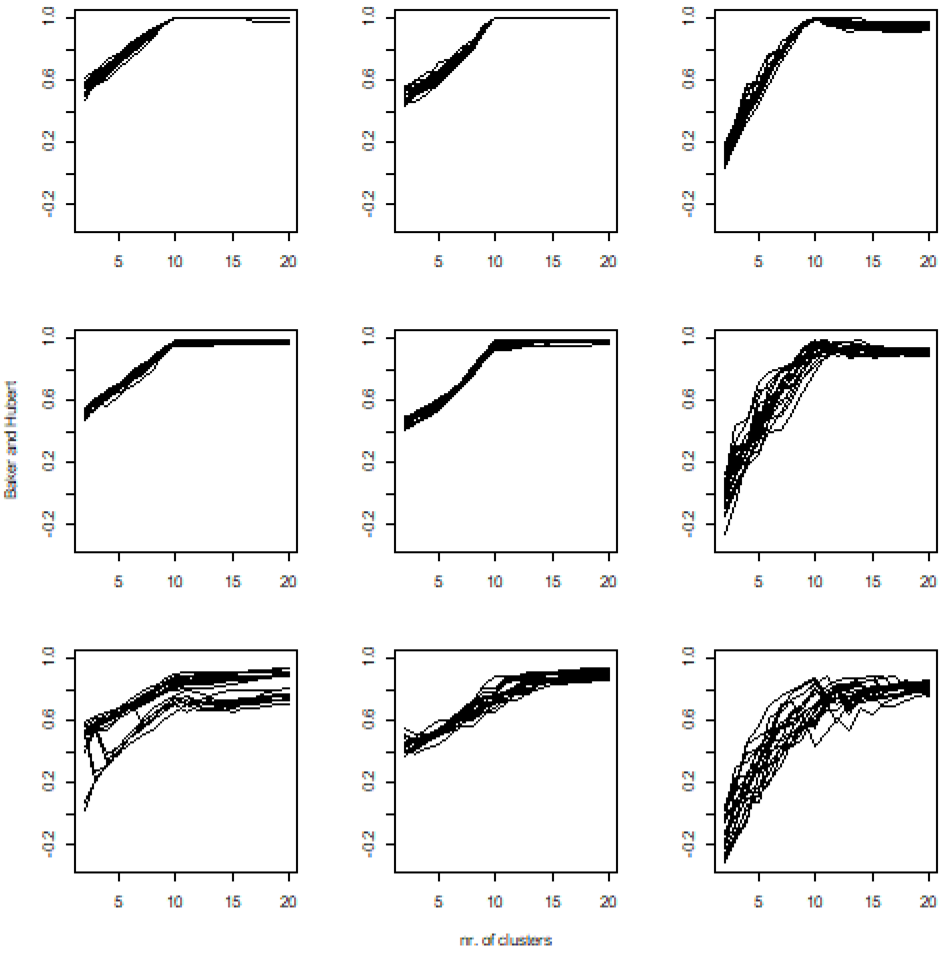


**Figure S2.4** Performance of Baker & Hubert index versus the number of clusters at different noise levels. Rows indicate noise level (upper is the low noise), and columns are algorithms (left = pam, medium = UPGMA, right = flexible).

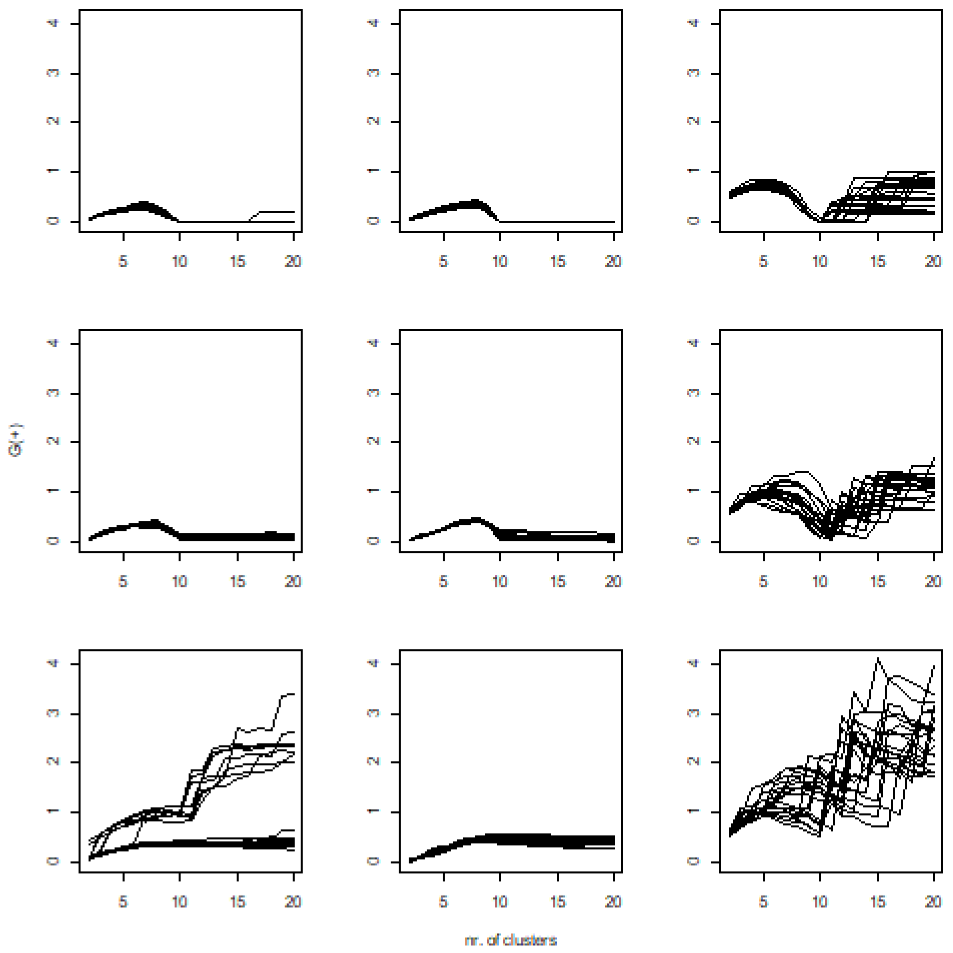


**Figure S2.5** Performance of G (+) index versus the number of clusters at different noise levels. Rows indicate noise levels (upper is the low noise), and columns are algorithms (left = pam, medium = UPGMA, right = flexible).

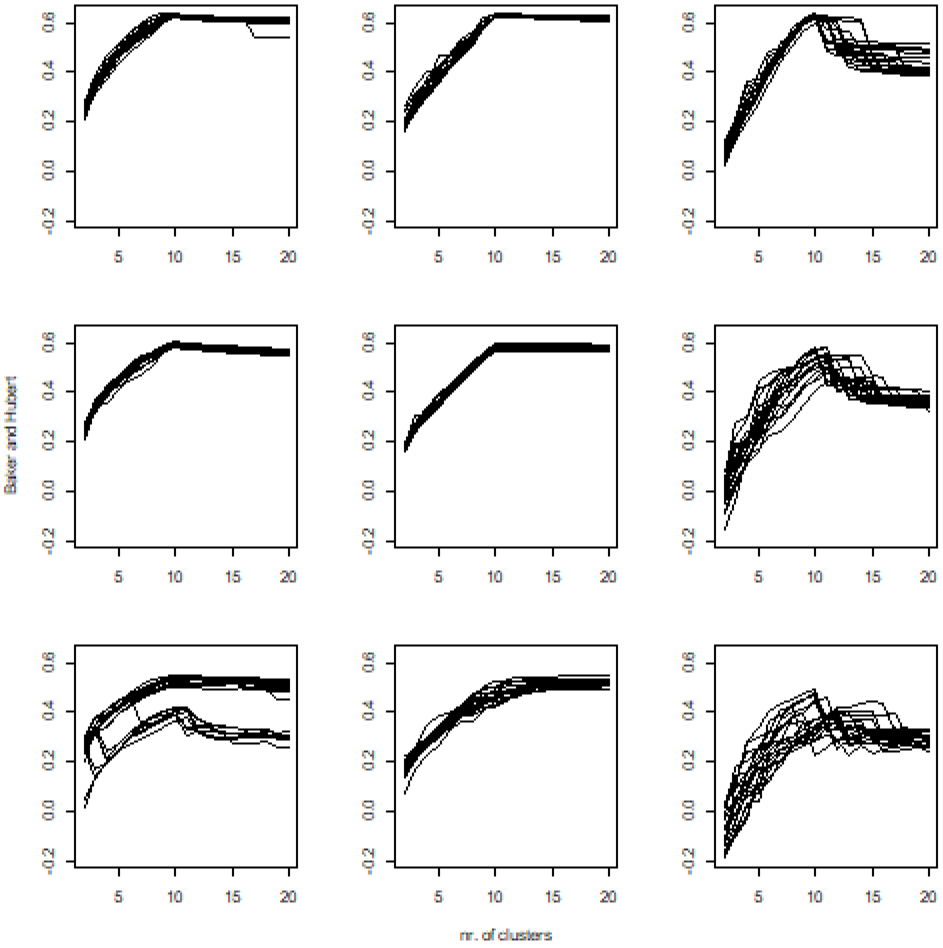


**Figure S2.6** Performance of Tau index versus the number of clusters at different noise levels. Rows indicate noise level (upper is the low noise), and columns are algorithms (left = pam, medium = UPGMA, right = flexible).

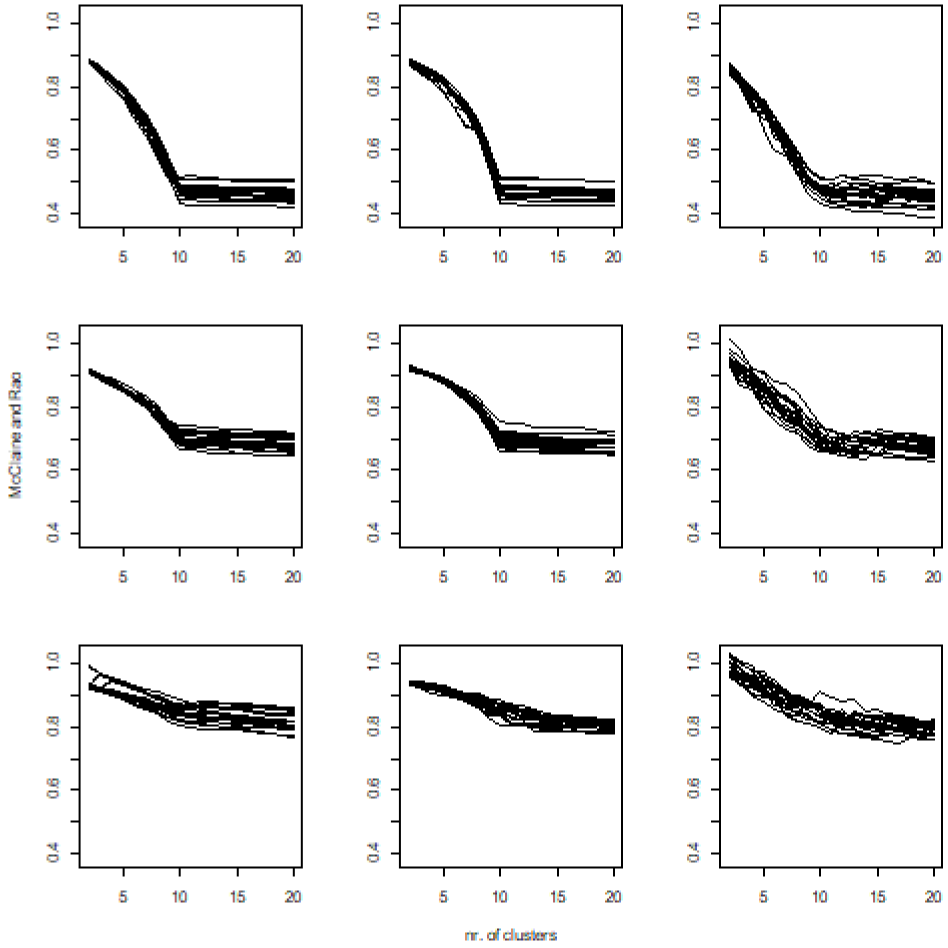


**Figure S2.7** Performance of McClaine & Rao index versus the number of clusters at different noise levels. Rows indicate noise level (upper is the low noise), and columns are algorithms (left = pam, medium = UPGMA, right = flexible).

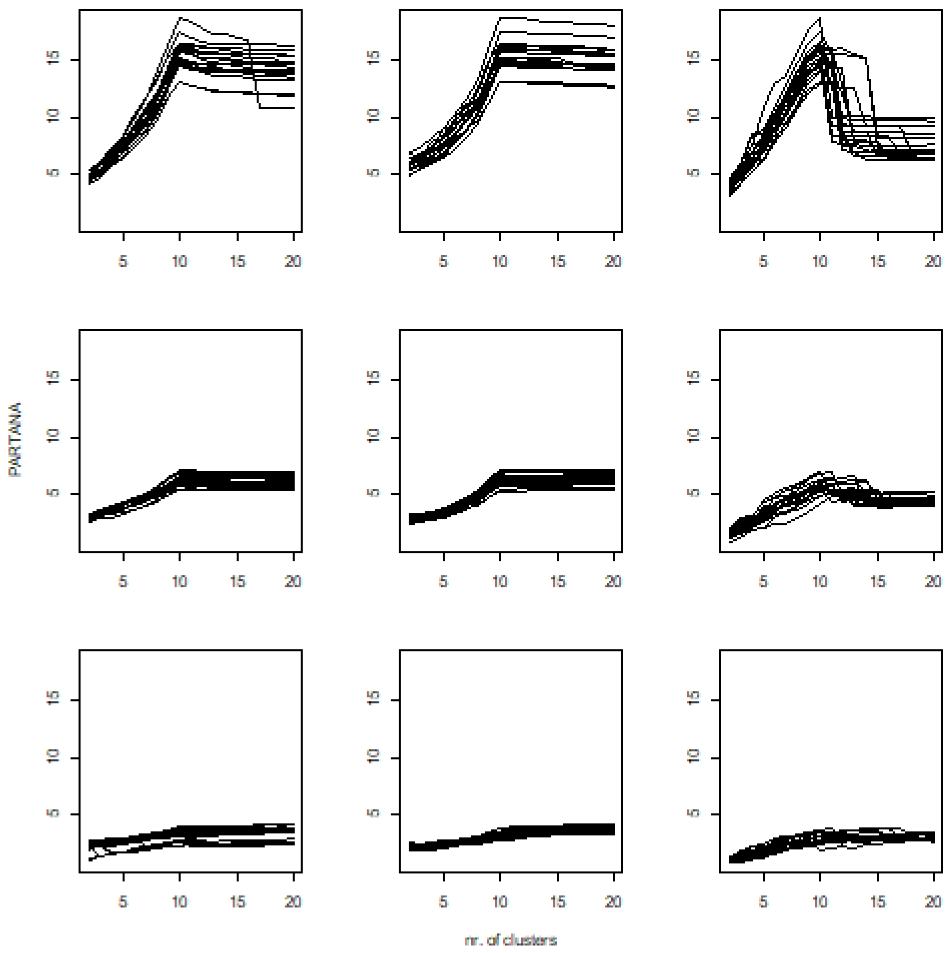


**Figure S2.8** Performance of PARTANA index versus the number of clusters at different noise levels. Rows indicate noise level (upper is the low noise), and columns are algorithms (left = pam, medium = UPGMA, right = flexible).

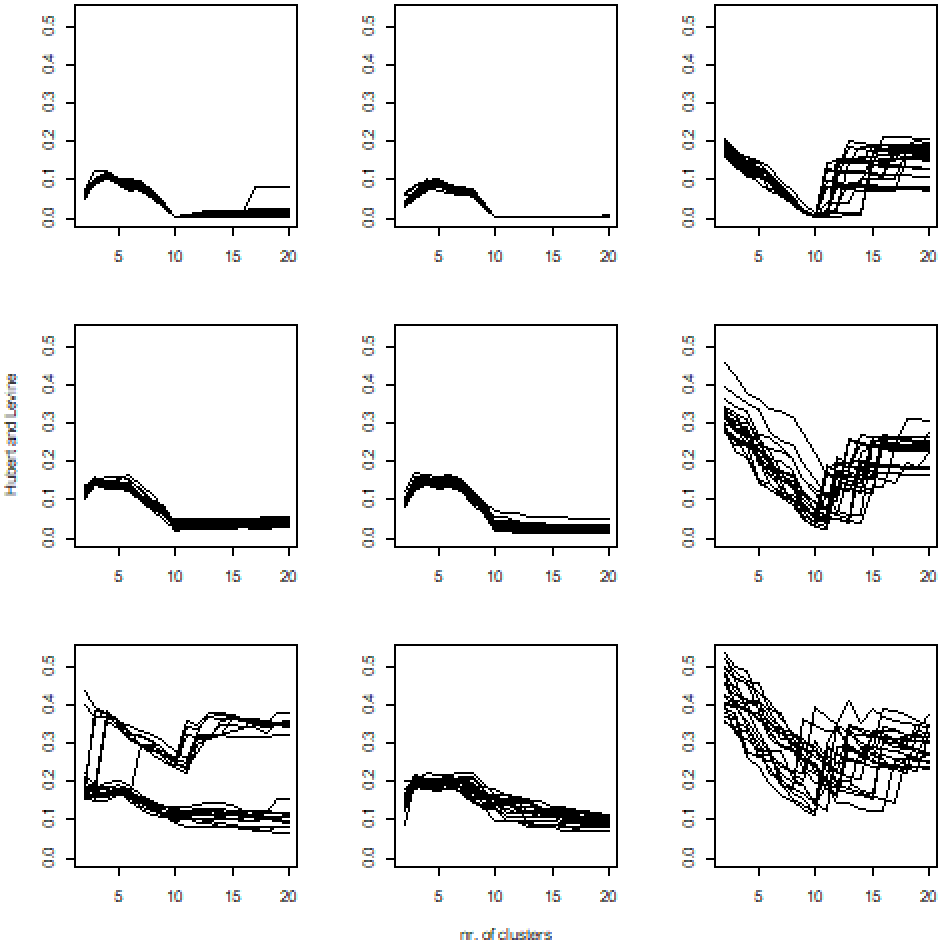


**Figure S2.9** Performance of Hubert & Levine index versus the number of clusters at different noise levels. Rows indicate noise level (upper is the low noise), and columns are algorithms (left = pam, medium = UPGMA, right = flexible).

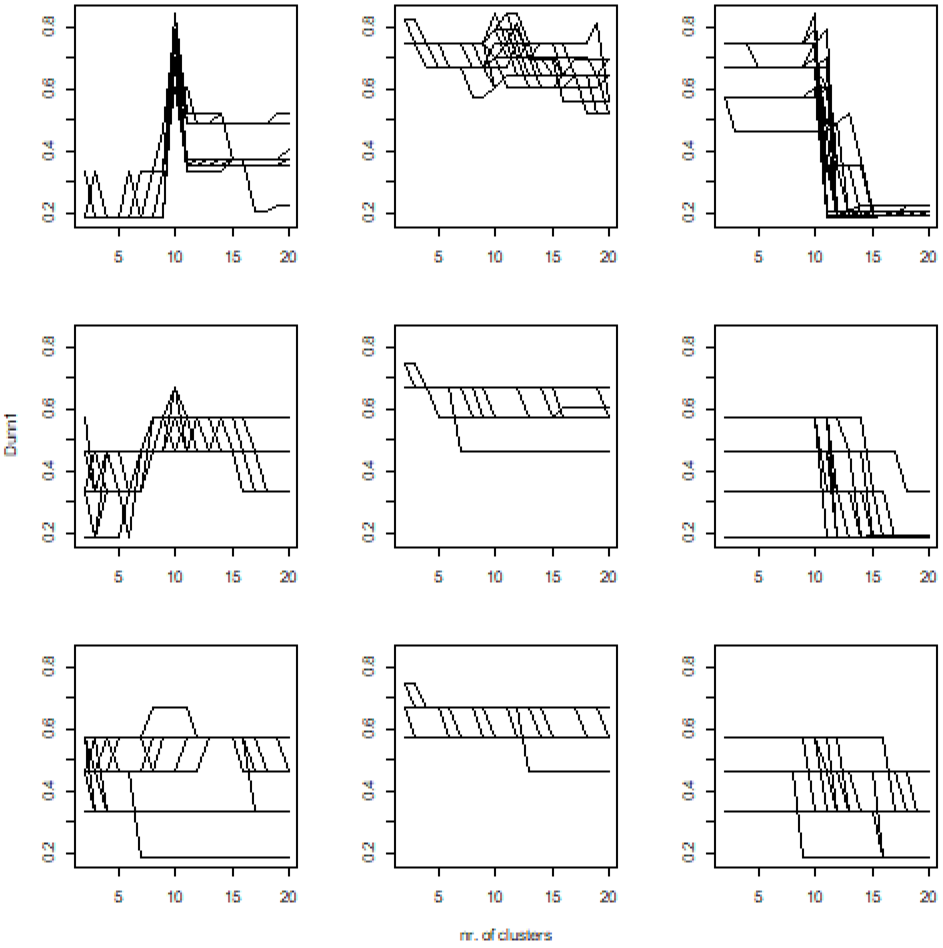


**Figure S2.10** Performance of $Dunn_{min\_max}$ index versus the number of clusters at different noise levels. Rows indicate noise level (upper is the low noise), and columns are algorithms (left = pam, medium = UPGMA, right = flexible).

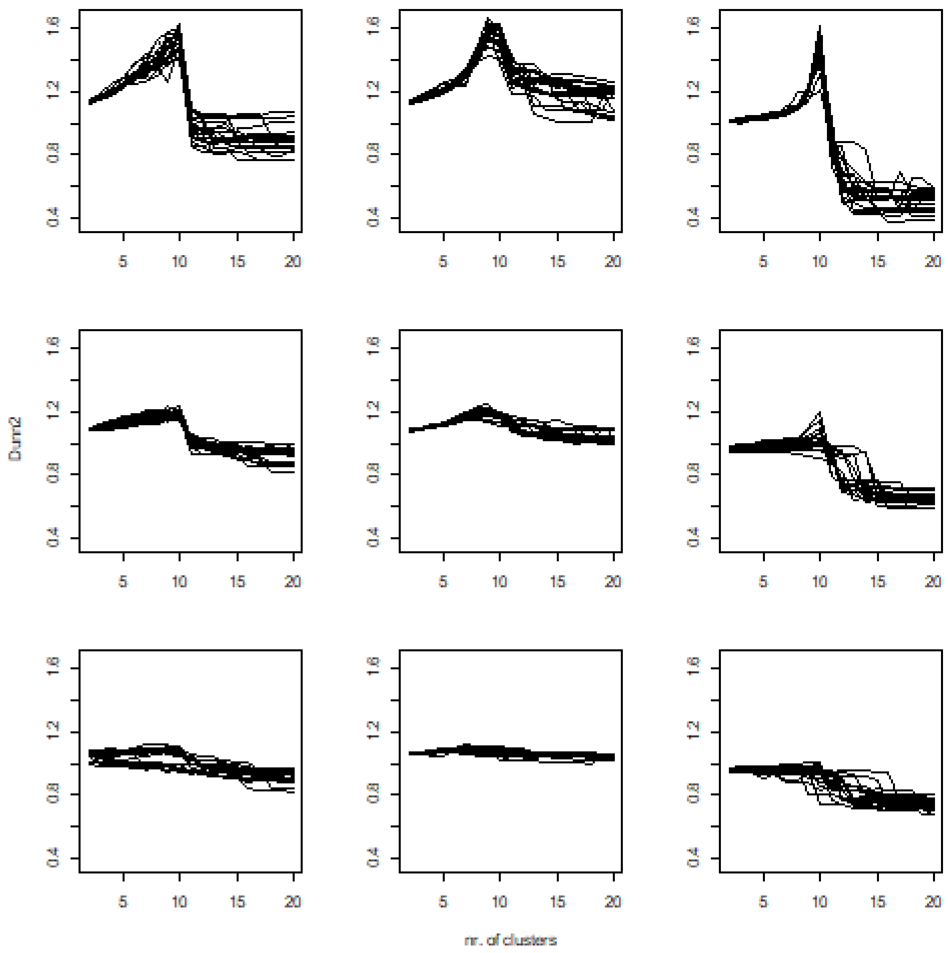


**Figure S2.11** Performance of $Dunn_{mean\_mean}$ index versus the number of clusters at different noise levels. Rows indicate noise level (upper is the low noise), and columns are algorithms (left = pam, medium = UPGMA, right = flexible).

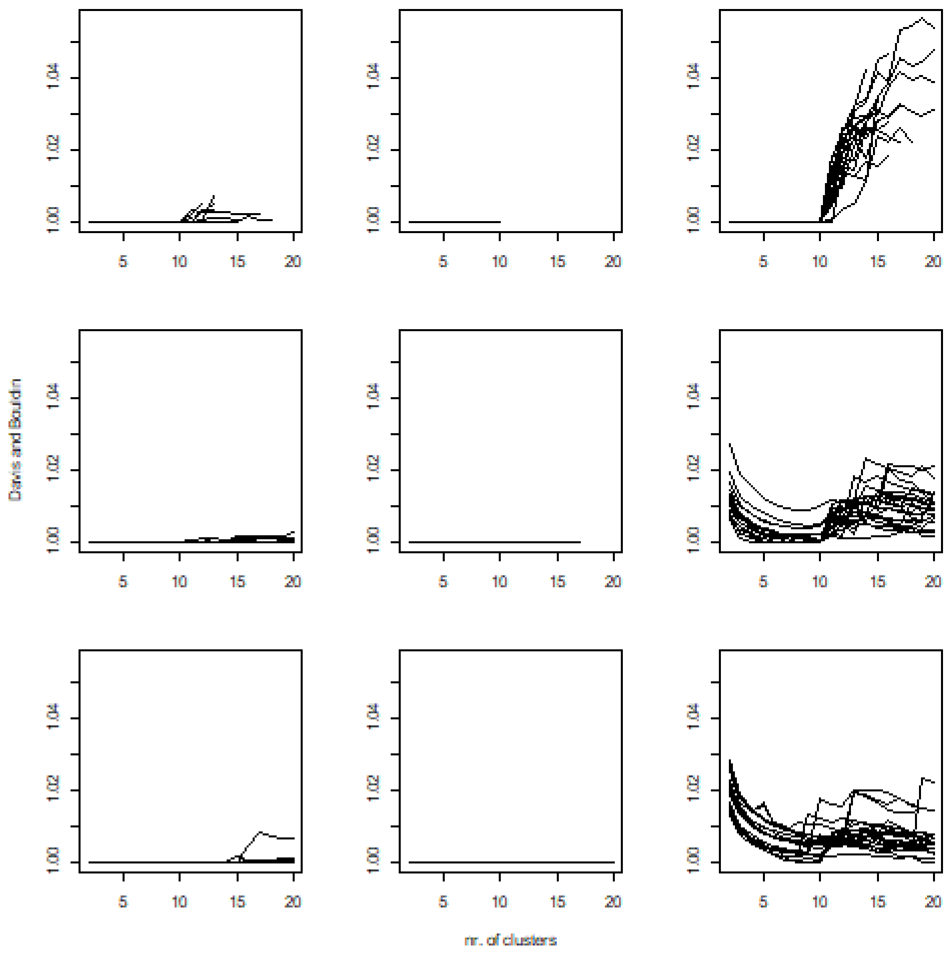


**Figure S2.12** Performance of Popma$_{unweighted}$ index versus the number of clusters at different noise levels. Rows indicate noise level (upper is the low noise), and columns are algorithms (left = pam, medium = UPGMA, right = flexible).

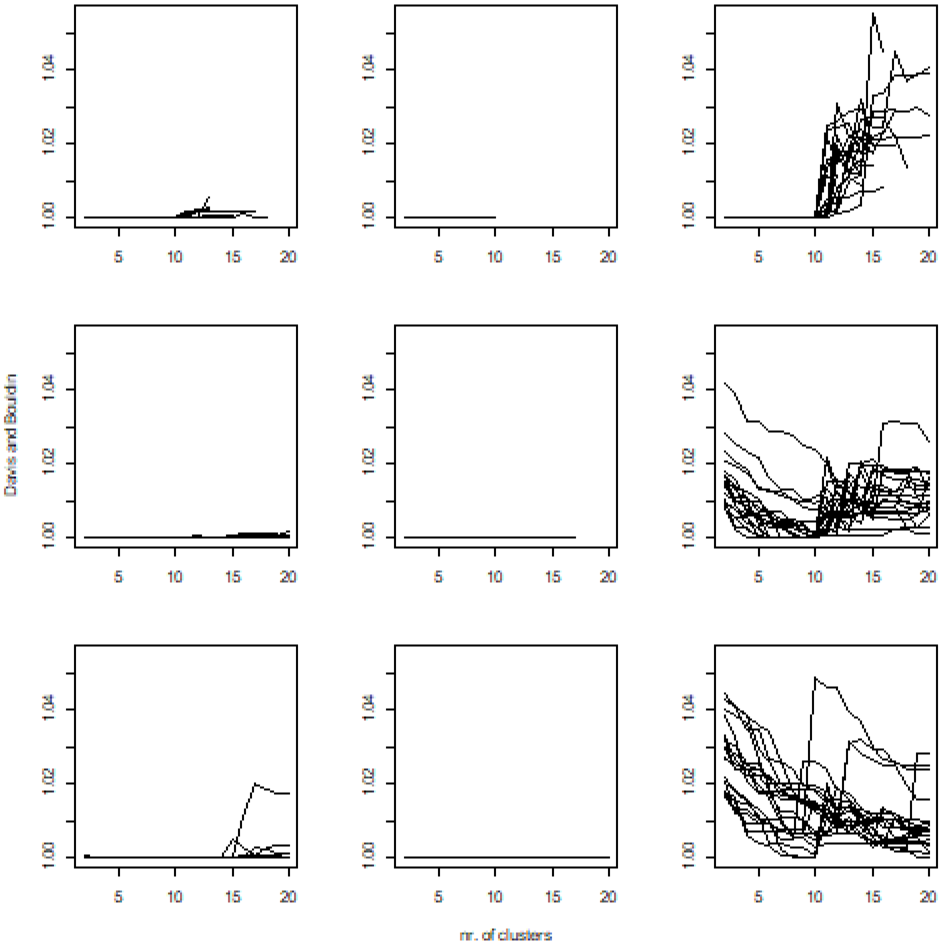


**Figure S2.13** Performance of $Popma_{weighted}$ index versus the number of clusters at different noise levels. Rows indicate noise level (upper is the low noise), and columns are algorithms (left = pam, medium = UPGMA, right = flexible).

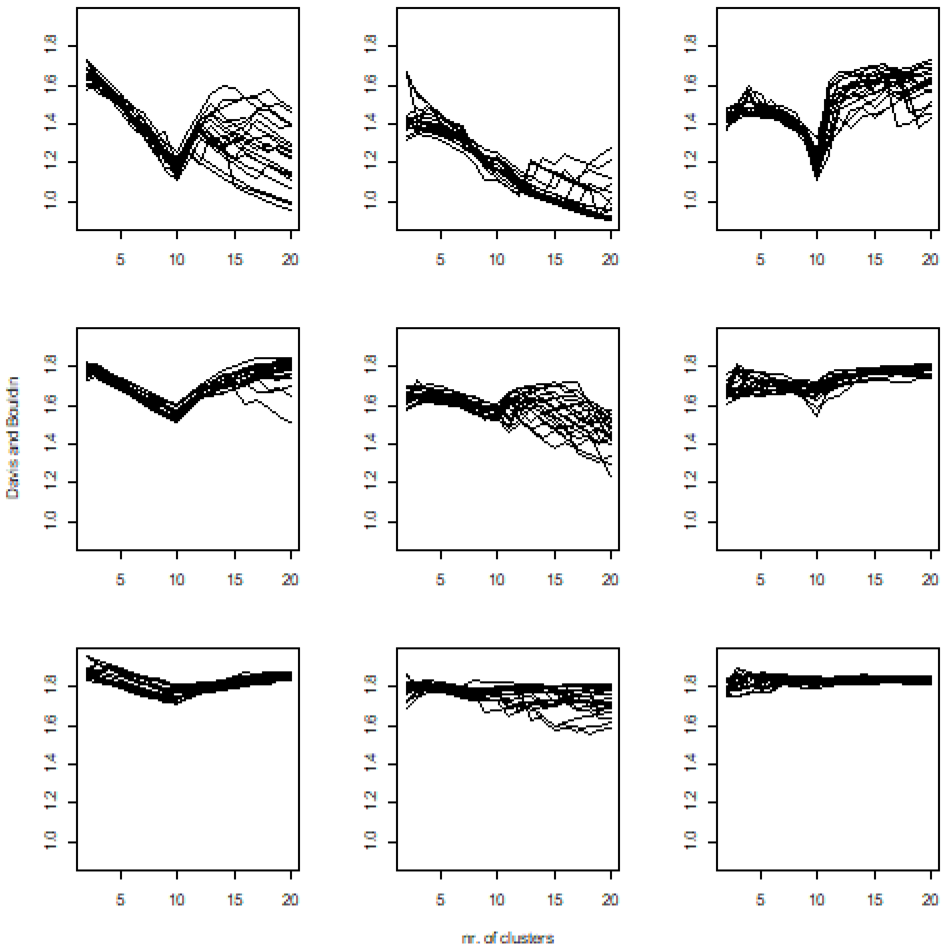


**Figure S2.14** Performance of Davis & Bouldin index versus the number of clusters at different noise levels. Rows indicate noise level (upper is the low noise), and columns are algorithms (left = pam, medium = UPGMA, right = flexible).

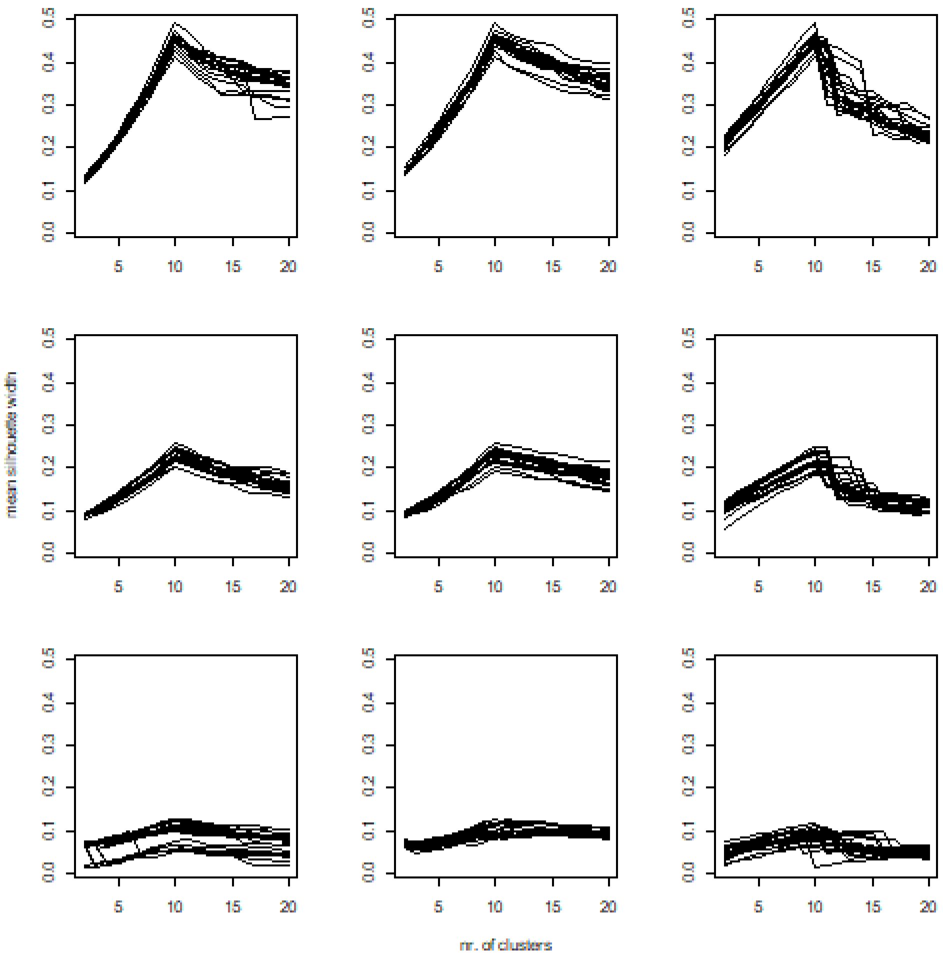


**Figure S2.14** Performance of mean silhouette width index versus the number of clusters at different noise levels. Rows indicate noise level (upper is the low noise), and columns are algorithms (left = pam, medium = UPGMA, right = flexible).

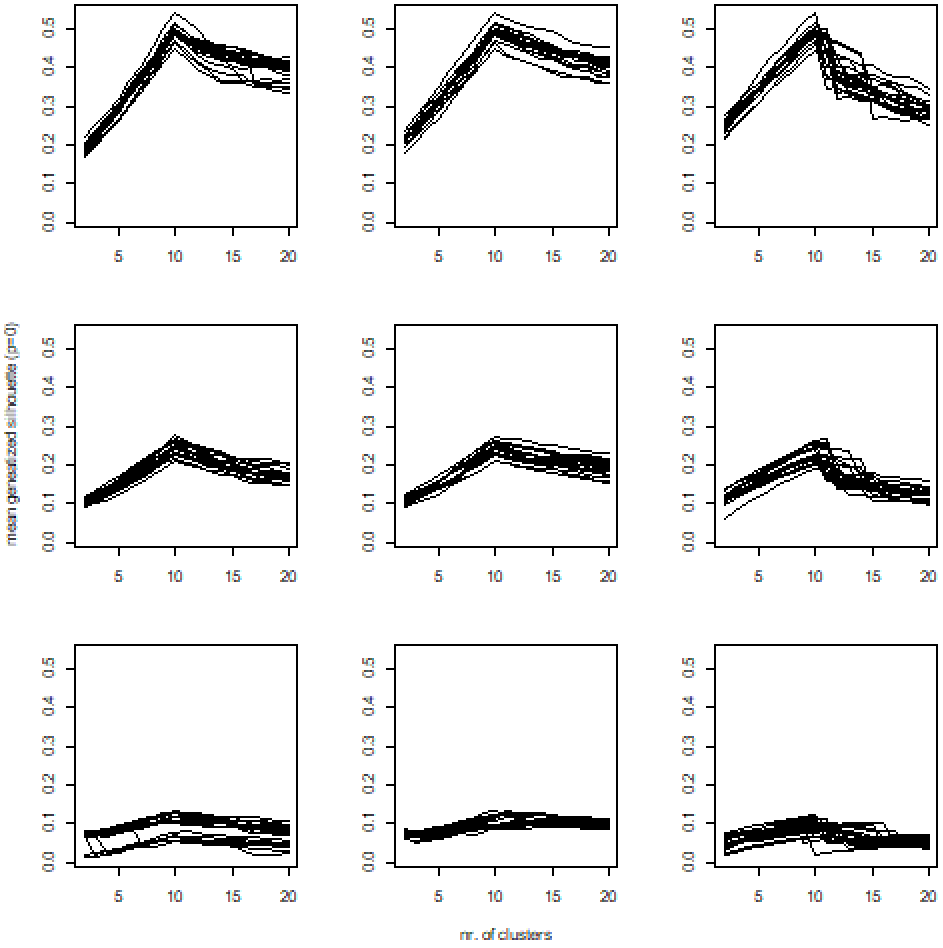


**Figure S2.15** Performance of gensil(0) index versus number of clusters at different noise levels. Rows indicate noise level (upper is the low noise), and columns are algorithms (left = pam, medium = UPGMA, right = flexible).

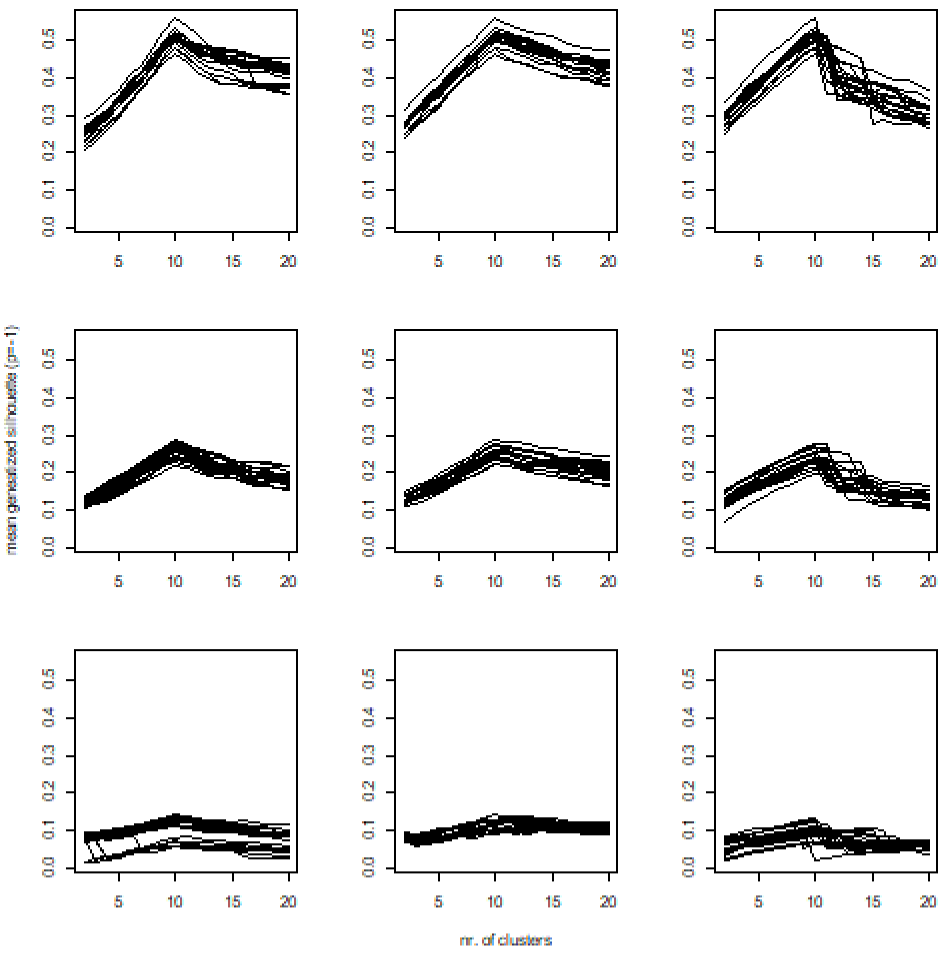


**Figure S2.16** Performance of gensil (-1) index versus the number of clusters at different noise levels. Rows indicate noise level (upper is the low noise), and columns are algorithms (left = pam, medium = UPGMA, right = flexible).

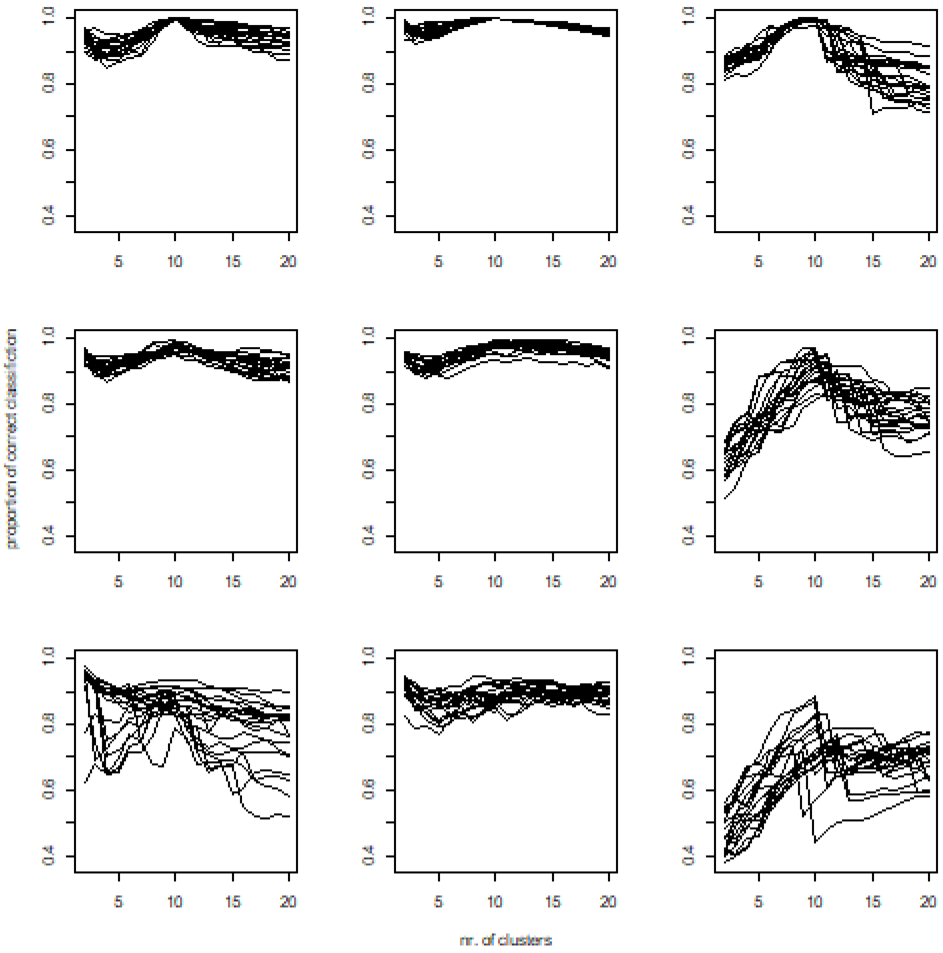


**Figure S2.17** Performance of mean proportion of correct classification index versus the number of clusters at different noise levels. Rows indicate noise level (upper is the low noise), and columns are algorithms (left = pam, medium = UPGMA, right = flexible).

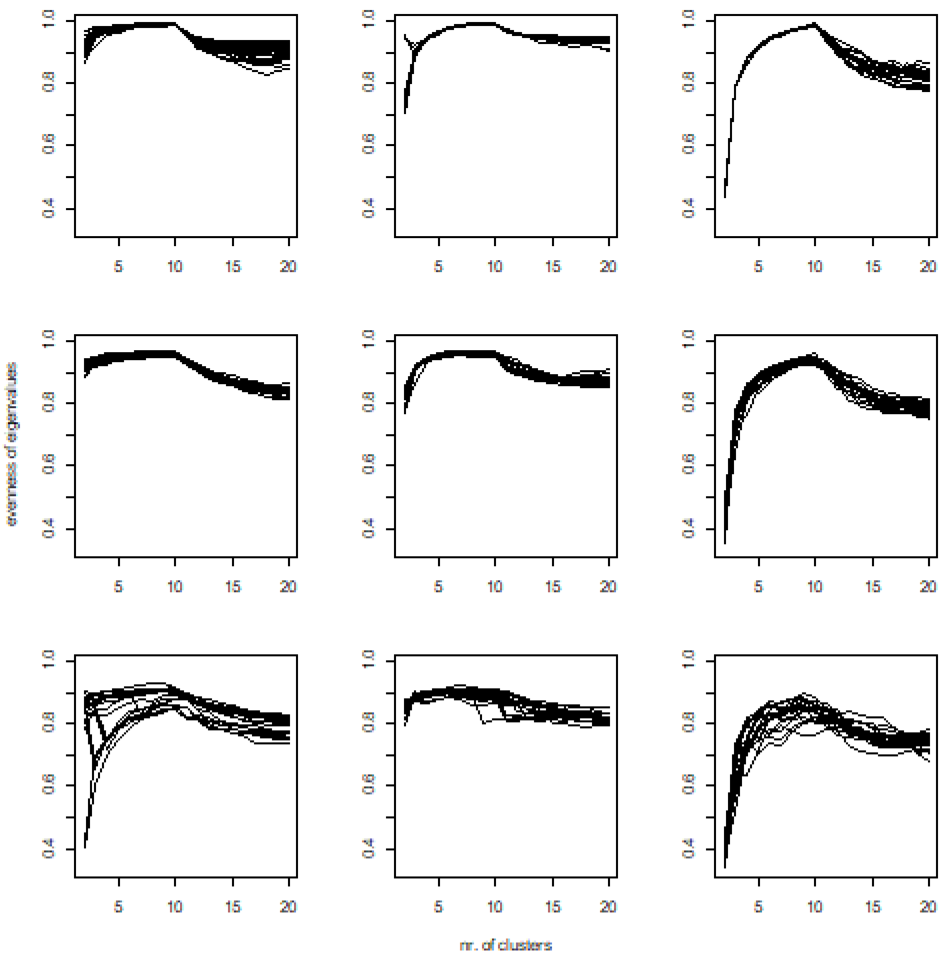


**Figure S2.18** Performance of evenness of eigenvalues index versus the number of clusters at different noise levels. Rows indicate noise level (upper is the low noise), and columns are algorithms (left = pam, medium = UPGMA, right = flexible).

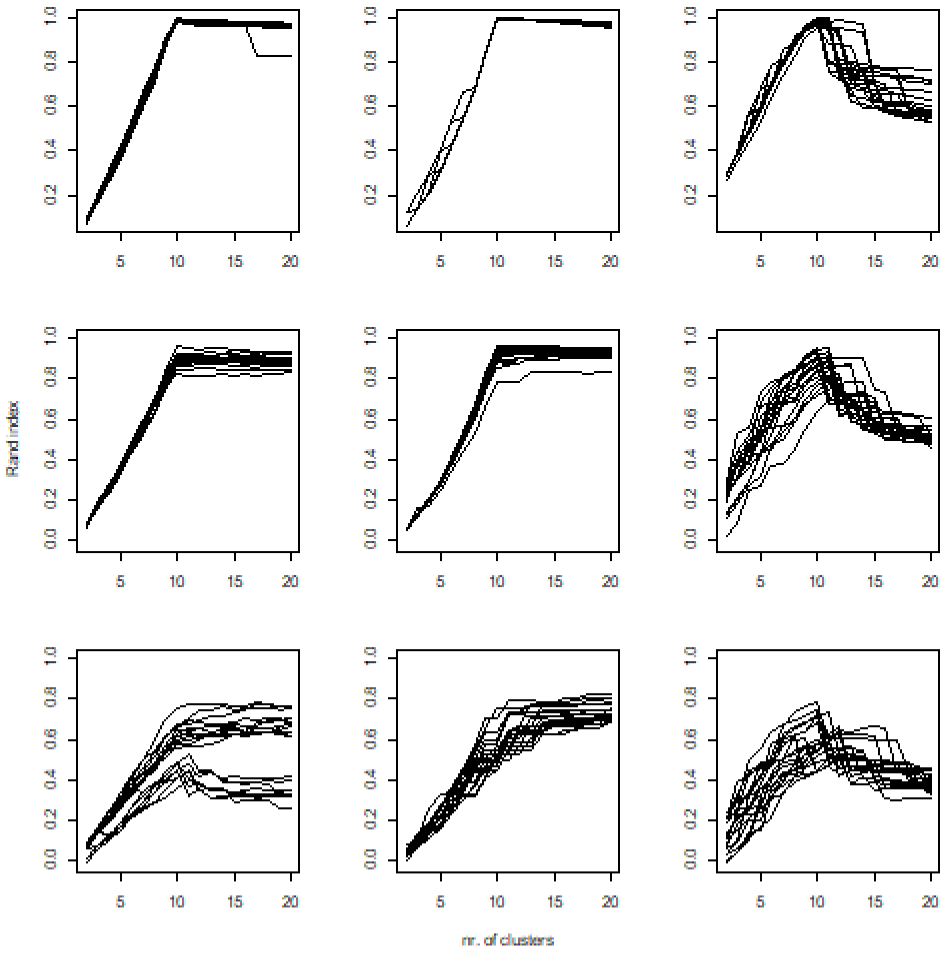


**Figure S2.19** Performance of Rand index versus the number of clusters at different noise levels. Rows indicate noise level (upper is the low noise), and columns are algorithms (left = pam, medium = UPGMA, right = flexible).
