## Appendix 1 for "Quantitative evaluation of internal clustering validation indices using binary datasets"

**Appendix 1.** Comparison of performance of the internal geometric indices for equal cluster sizes

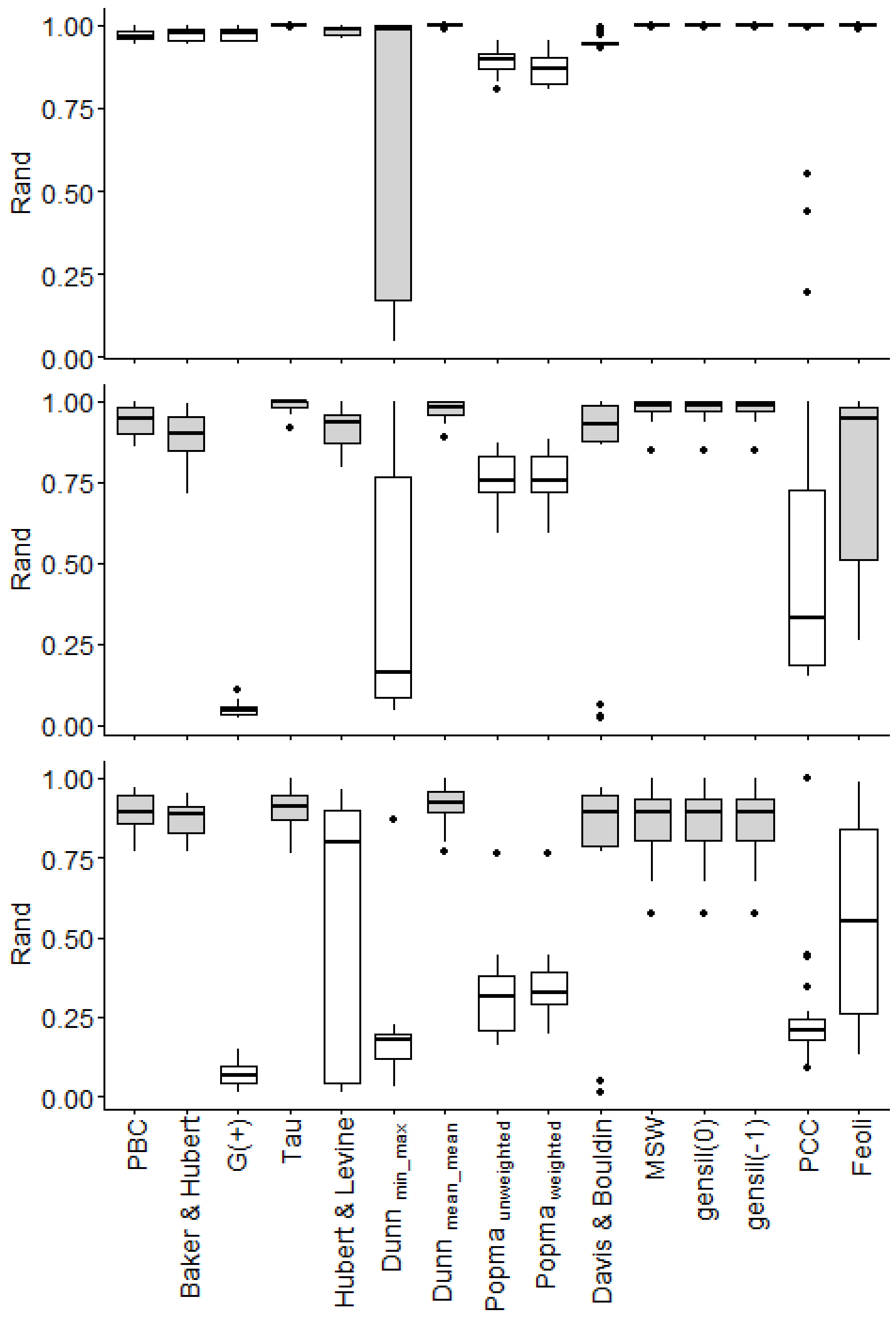


**Figure S1.1**-Performance of different geometric indices for each noise level (upper = low noise, bottom=high noise). (Gray bars indicate indices not significantly different from the best) Abbreviations: PBC = point-biserial correlation; MSW = mean silhouette widths; PCC = proportion of correct classification.

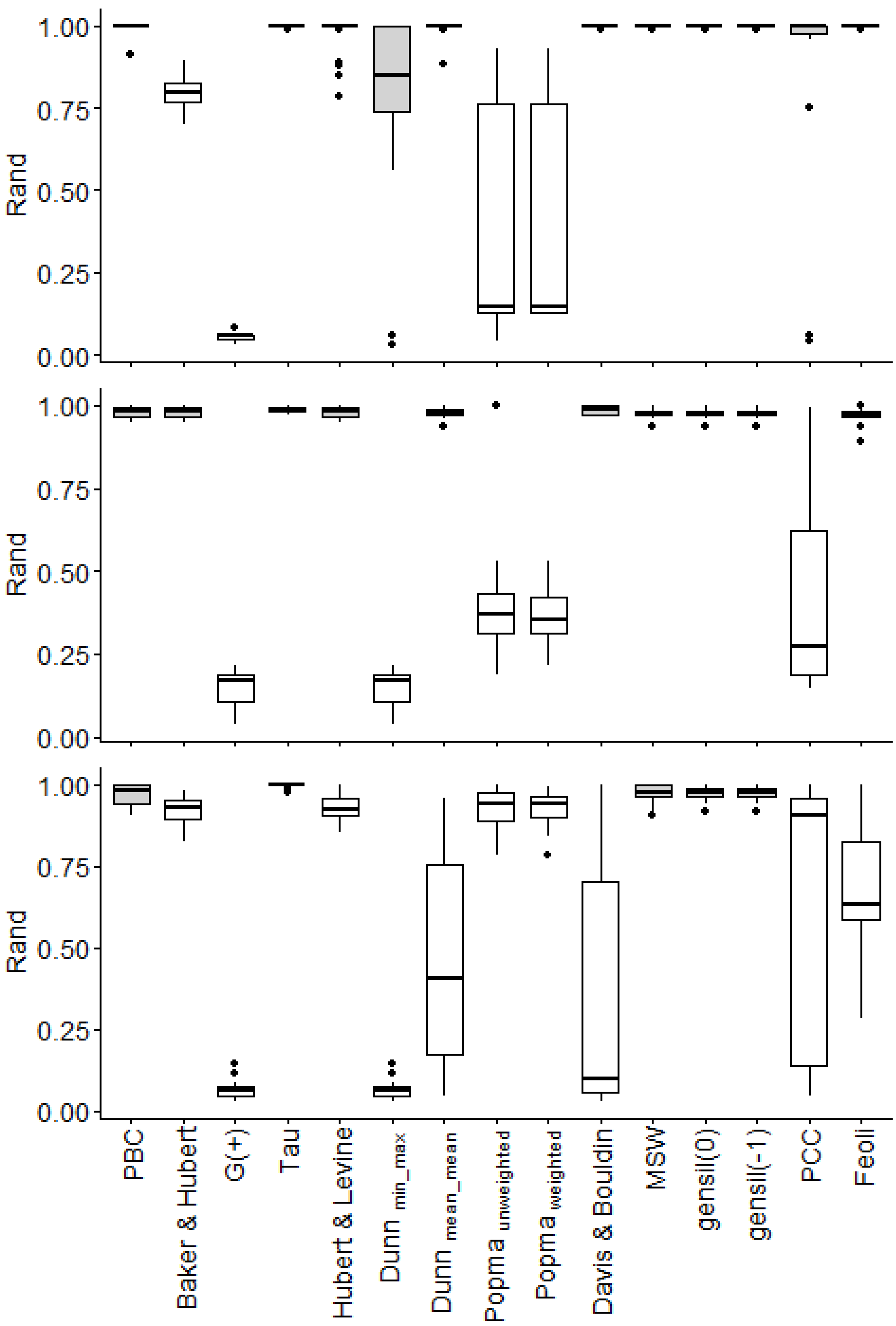


**Figure S1.2-**Performance of each clustering algorithm separately at the medium level of noise (upper=pam, medium = UPGMA, lower = flexible). (Gray bars indicate indices not significantly different from the best) Abbreviations: PBC = point-biserial correlation; MSW = mean silhouette widths; PCC = proportion of correct classification.

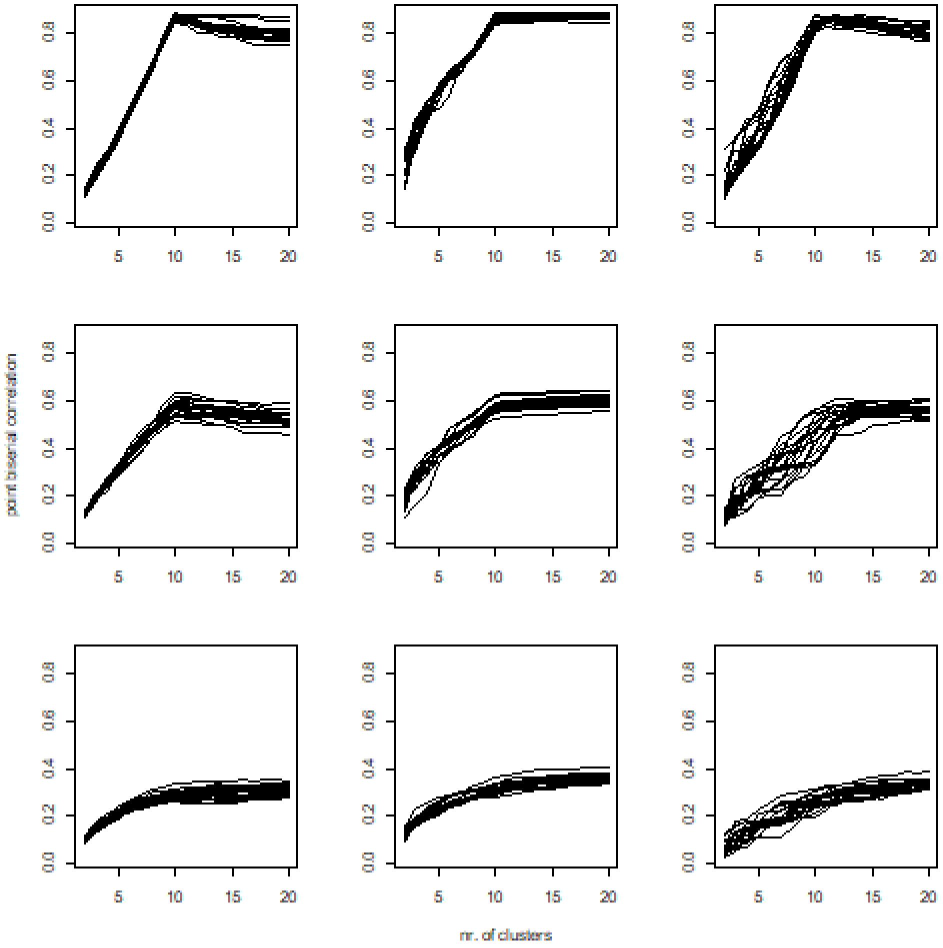


**Figure S1.3-**Performance of point. biserial index versus the number of clusters at different noise levels. Rows indicate noise level (upper is the low noise), and columns are algorithms (left = pam, medium = UPGMA, right = flexible).

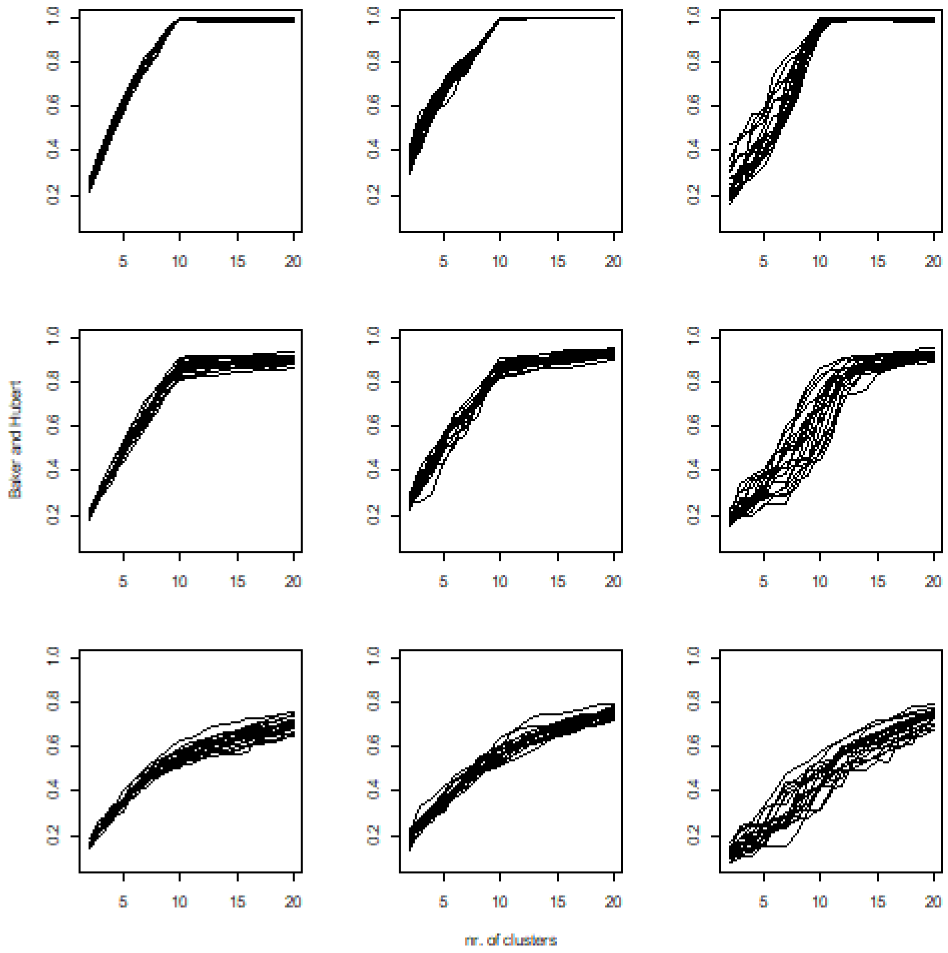


**Figure S1.4-**Performance of Baker & Hubert index versus the number of clusters at different noise levels. Rows indicate noise level (upper is the low noise), and columns are algorithms (left = pam, medium = UPGMA, right = flexible).

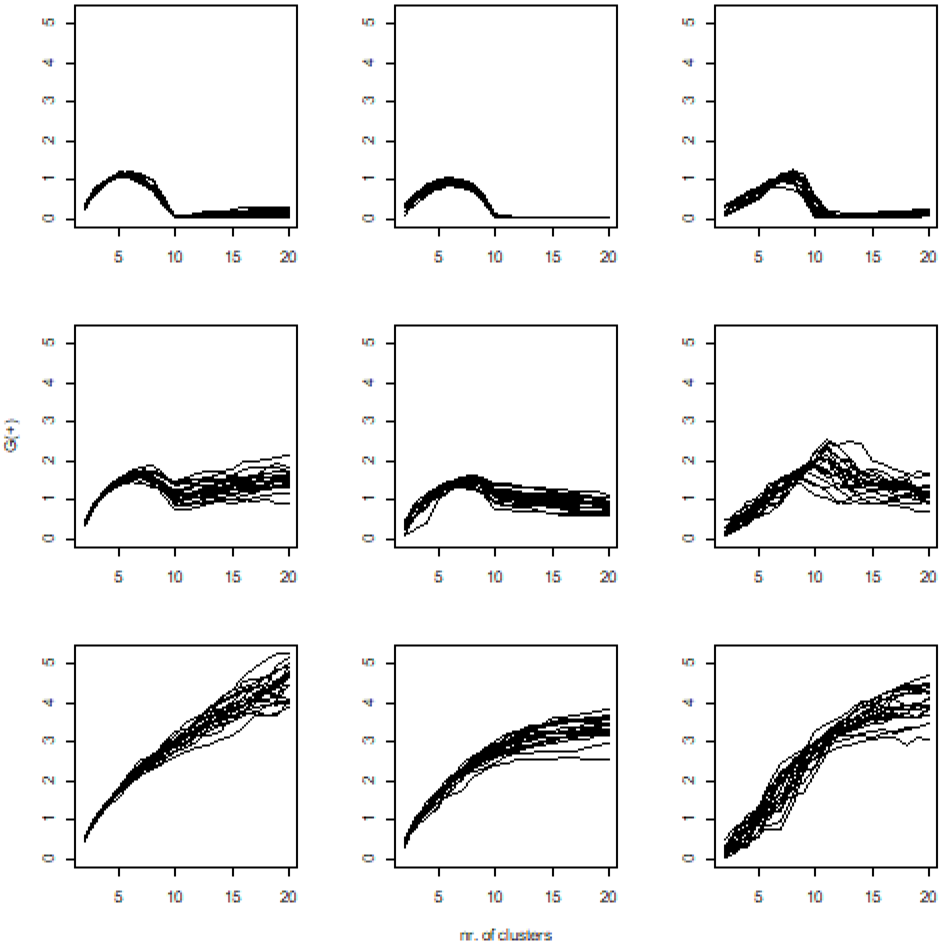


**Figure S1.5-**Performance of G (+) index versus the number of clusters at different noise levels. Rows indicate noise levels (upper is the low noise), and columns are algorithms (left = pam, medium = UPGMA, right = flexible).

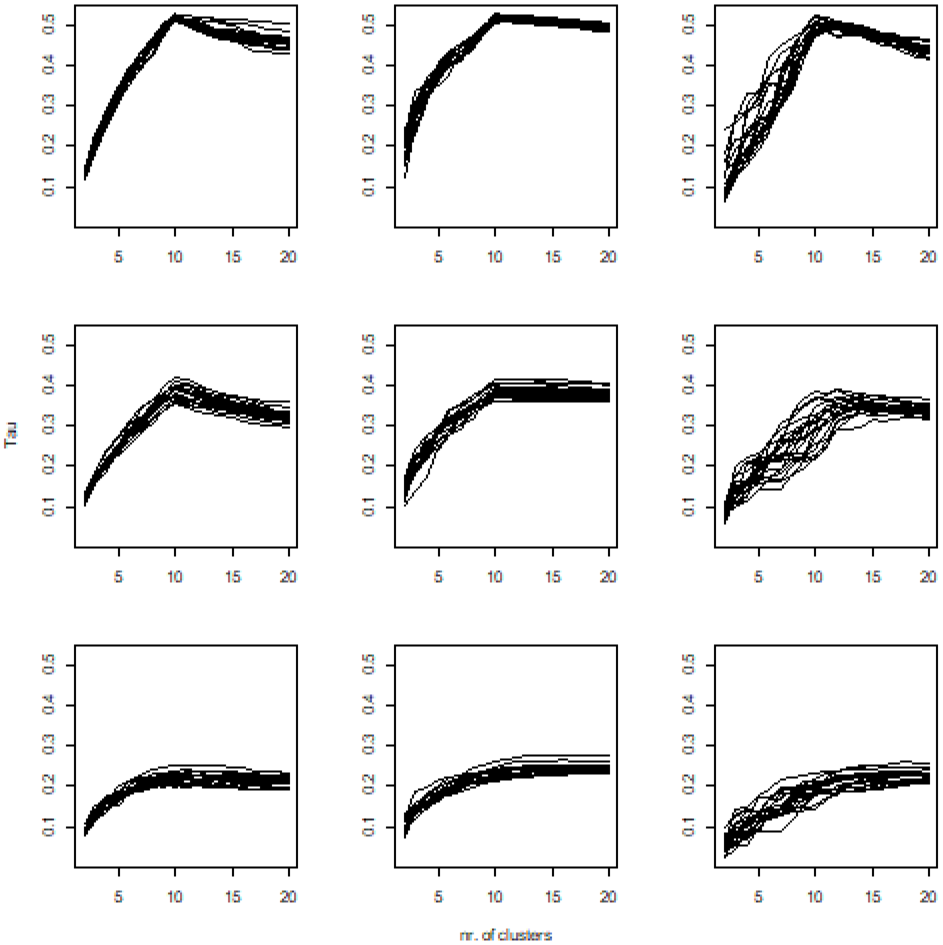


**Figure S1.6**-Performance of Tau index versus the number of clusters at different noise levels. Rows indicate noise level (upper is the low noise), and columns are algorithms (left = pam, medium = UPGMA, right = flexible).

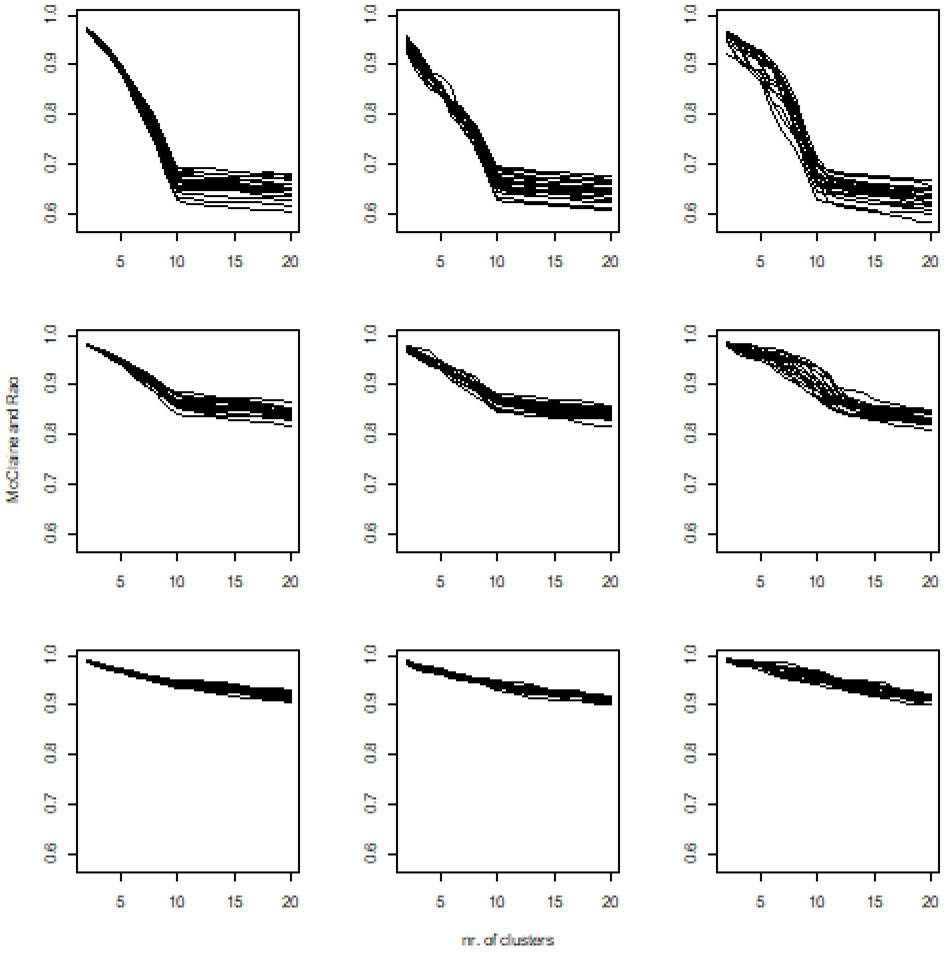


**Figure S1.7-**Performance of McClaine & Rao index versus the number of clusters at different noise levels. Rows indicate noise level (upper is the low noise), and columns are algorithms (left = pam, medium = UPGMA, right = flexible).

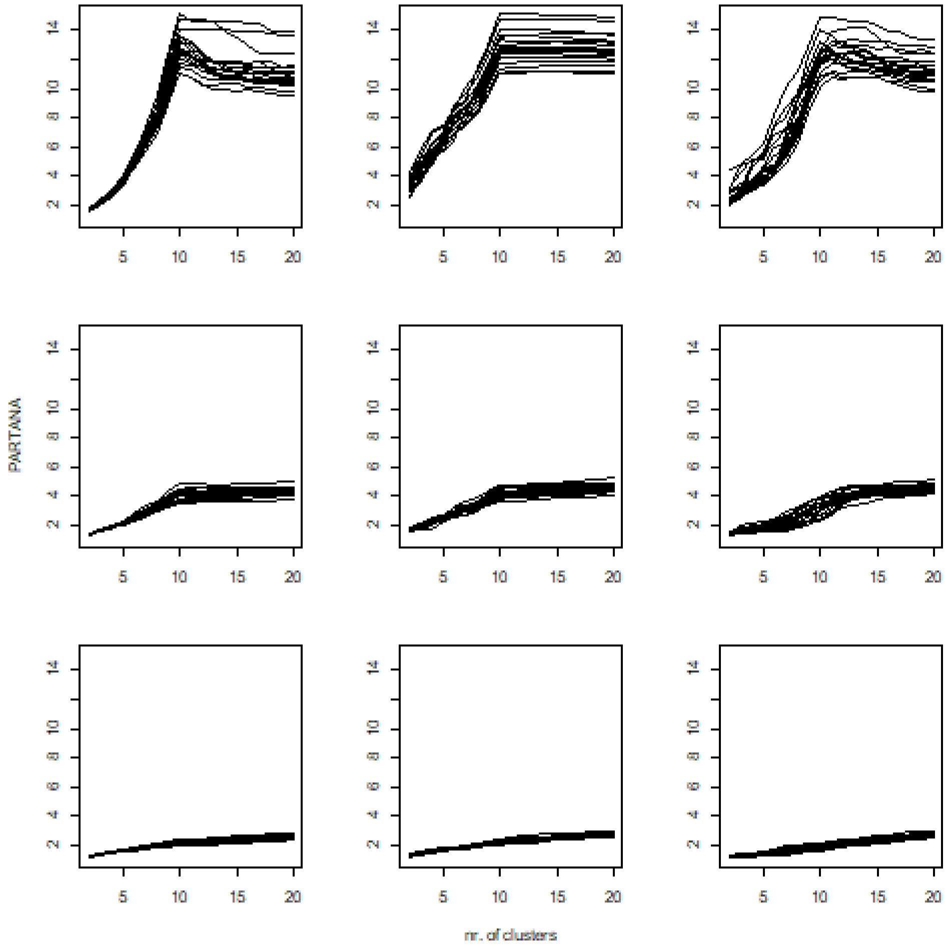


**Figure S1.8**-Performance of PARTANA index versus the number of clusters at different noise levels. Rows indicate noise level (upper is the low noise), and columns are algorithms (left = pam, medium = UPGMA, right = flexible).

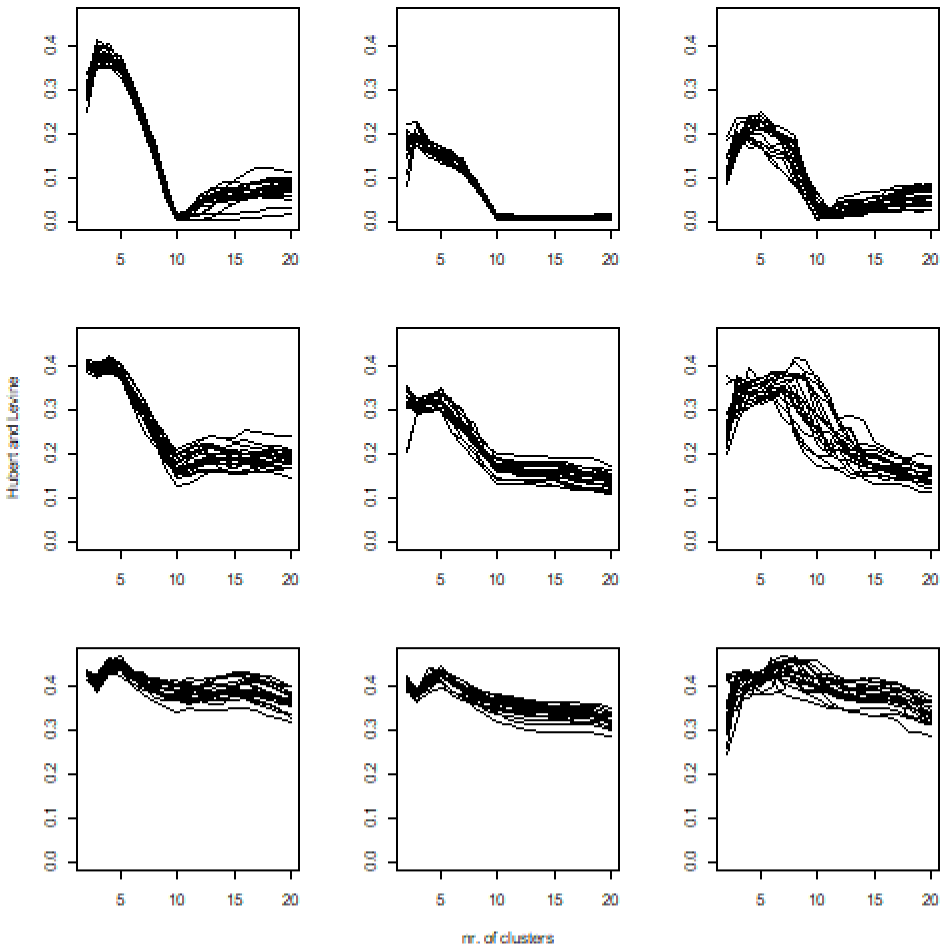


**Figure S1.9-**Performance of Hubert & Levine index versus the number of clusters at different noise levels. Rows indicate noise level (upper is the low noise), and columns are algorithms (left = pam, medium = UPGMA, right = flexible).

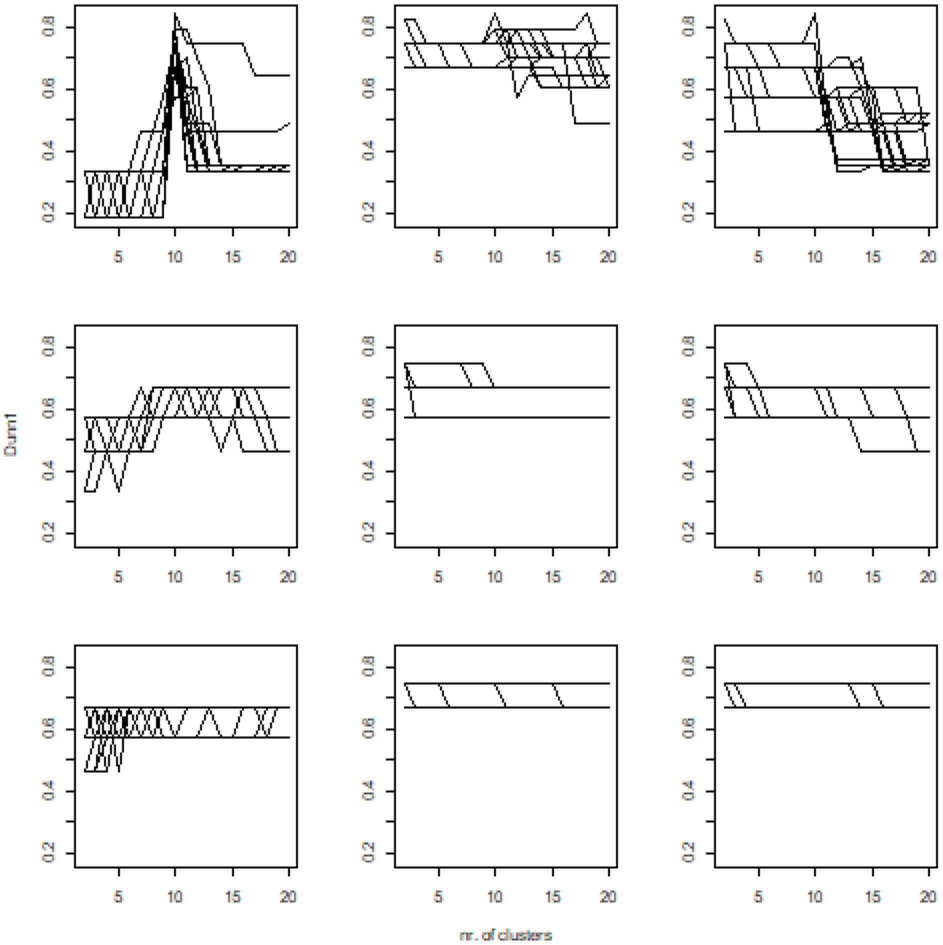


**Figure S1.10-**Performance of Dunn1index versus the number of clusters at different noise levels. Rows indicate noise level (upper is the low noise), and columns are algorithms (left = pam, medium = UPGMA, right = flexible).

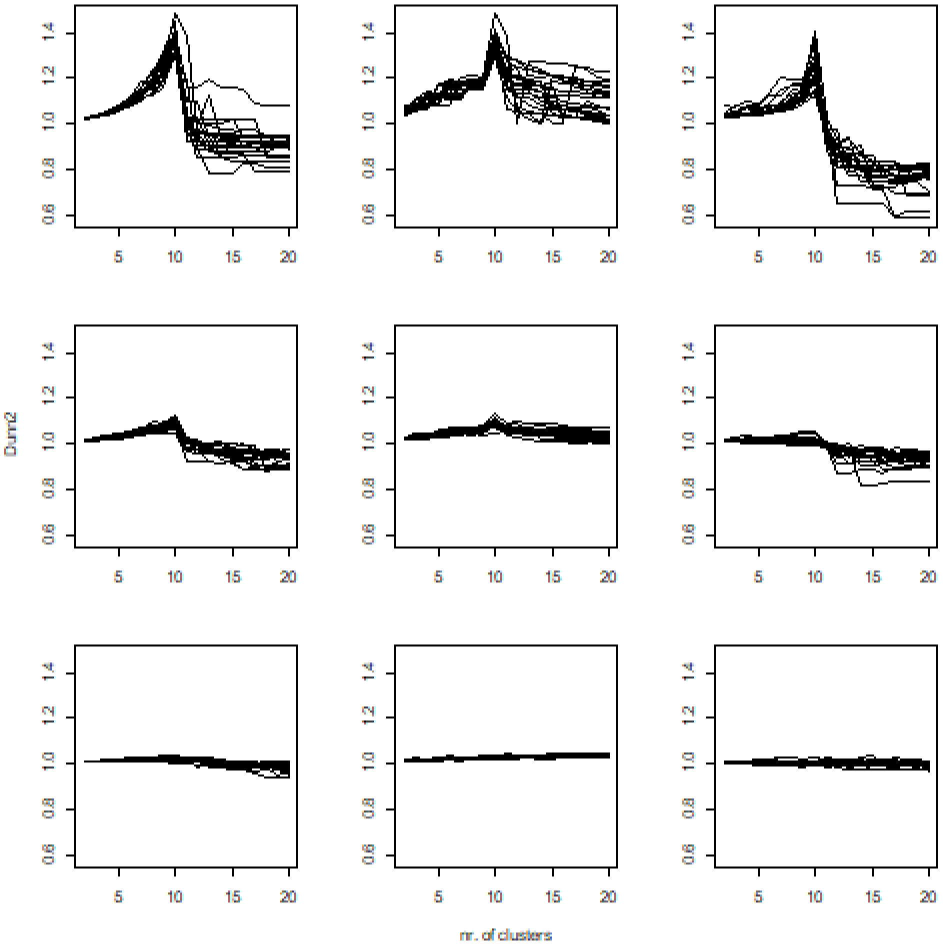


**Figure S1.11-**Performance of Dunn2 index versus the number of clusters at different noise levels. Rows indicate noise level (upper is the low noise), and columns are algorithms (left = pam, medium = UPGMA, right = flexible).

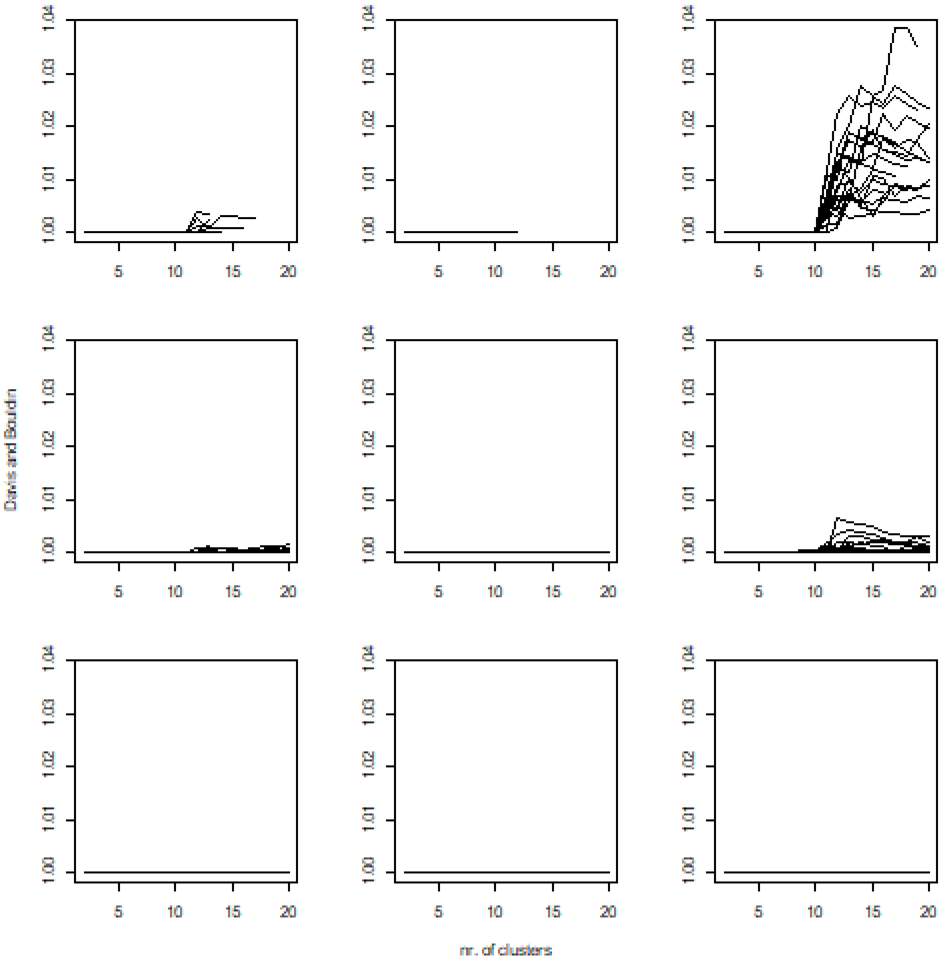


**Figure S1.12** Performance of Popma.unweighted index versus the number of clusters at different noise levels. Rows indicate noise level (upper is the low noise), and columns are algorithms (left = pam, medium = UPGMA, right = flexible).

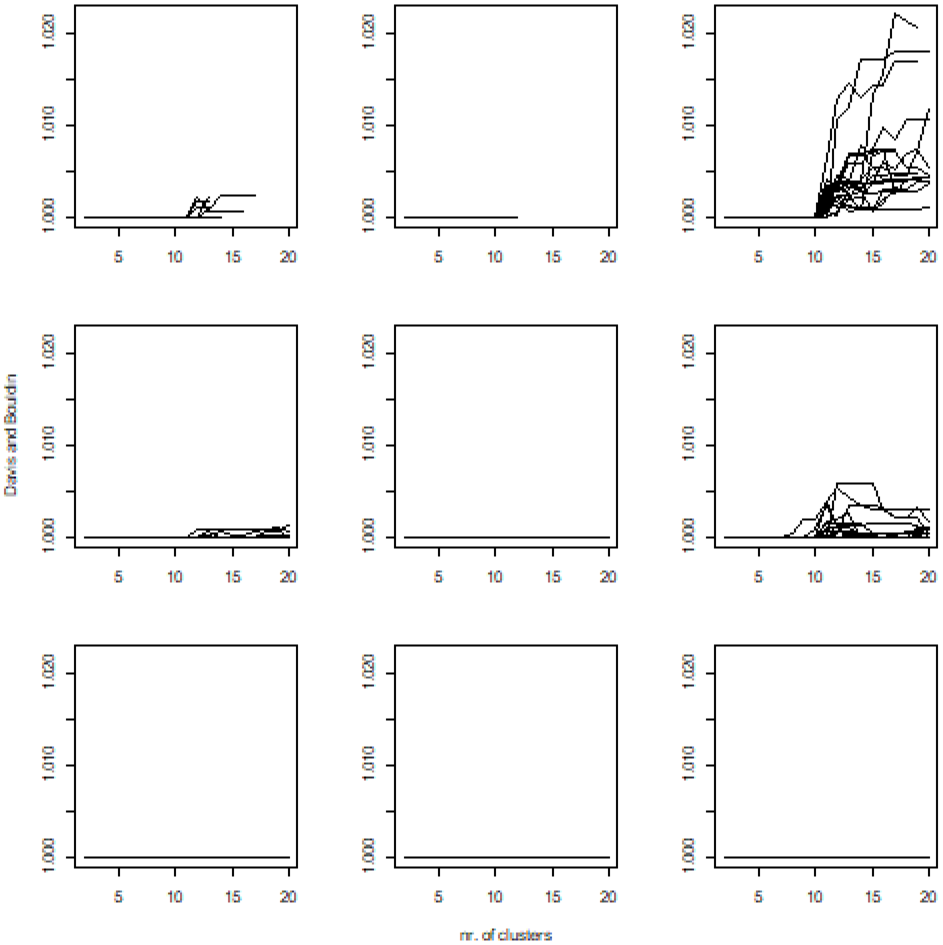


**Figure S1.13** Performance of Popma. weighted index versus the number of clusters at different noise levels. Rows indicate noise level (upper is the low noise), and columns are algorithms (left = pam, medium = UPGMA, right = flexible).

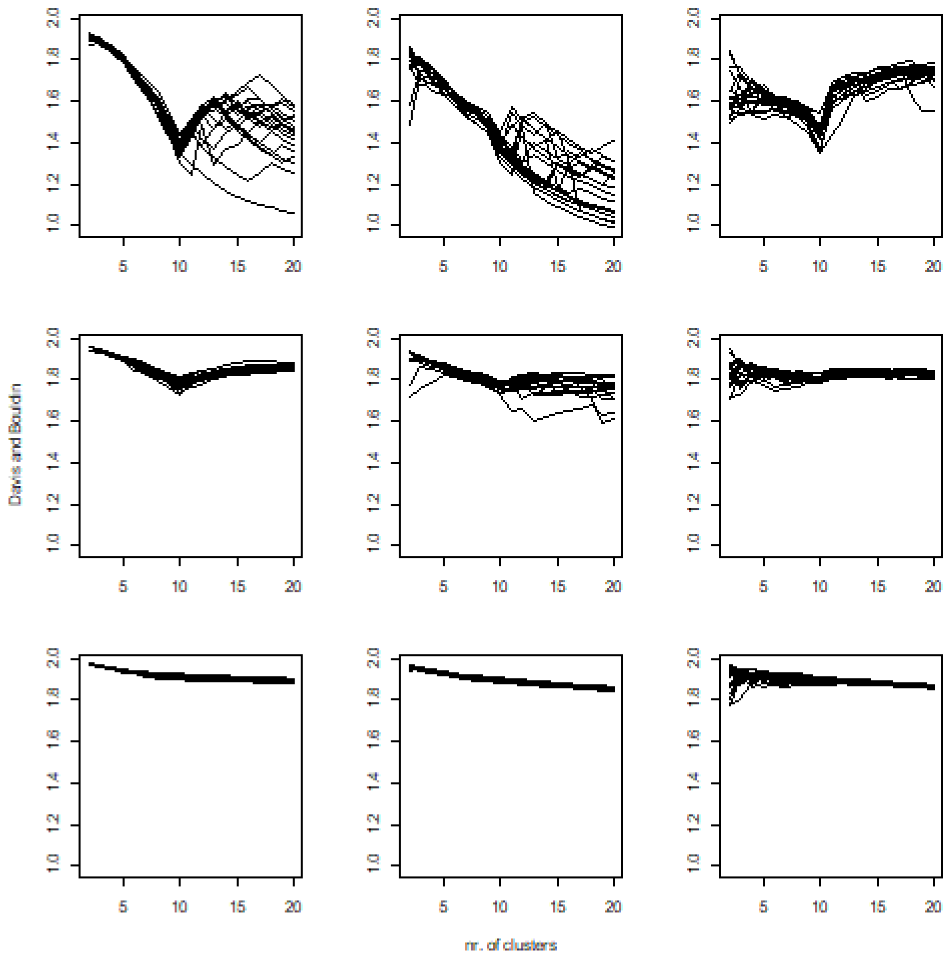


**Figure S1.14** Performance of Davis & Bouldin index versus the number of clusters at different noise levels. Rows indicate noise level (upper is the low noise), and columns are algorithms (left = pam, medium = UPGMA, right = flexible).

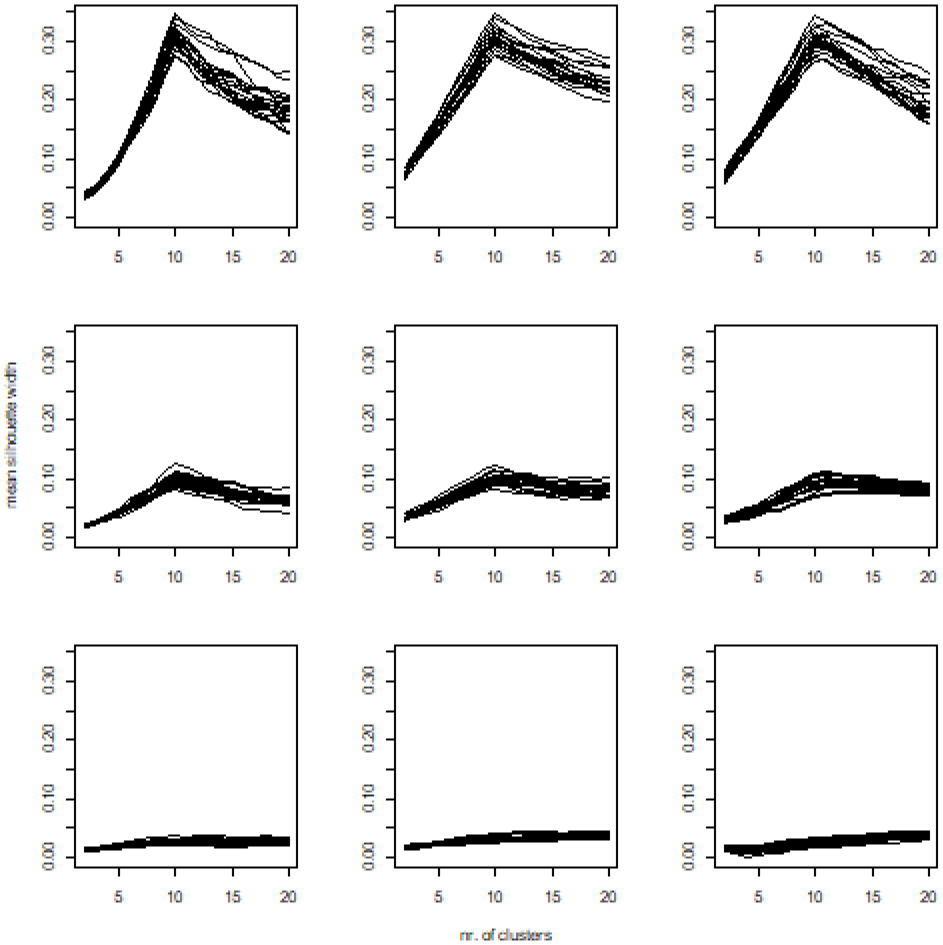


**Figure S1.15** Performance of mean silhouette width index versus the number of clusters at different noise levels. Rows indicate noise level (upper is the low noise), and columns are algorithms (left = pam, medium = UPGMA, right = flexible).

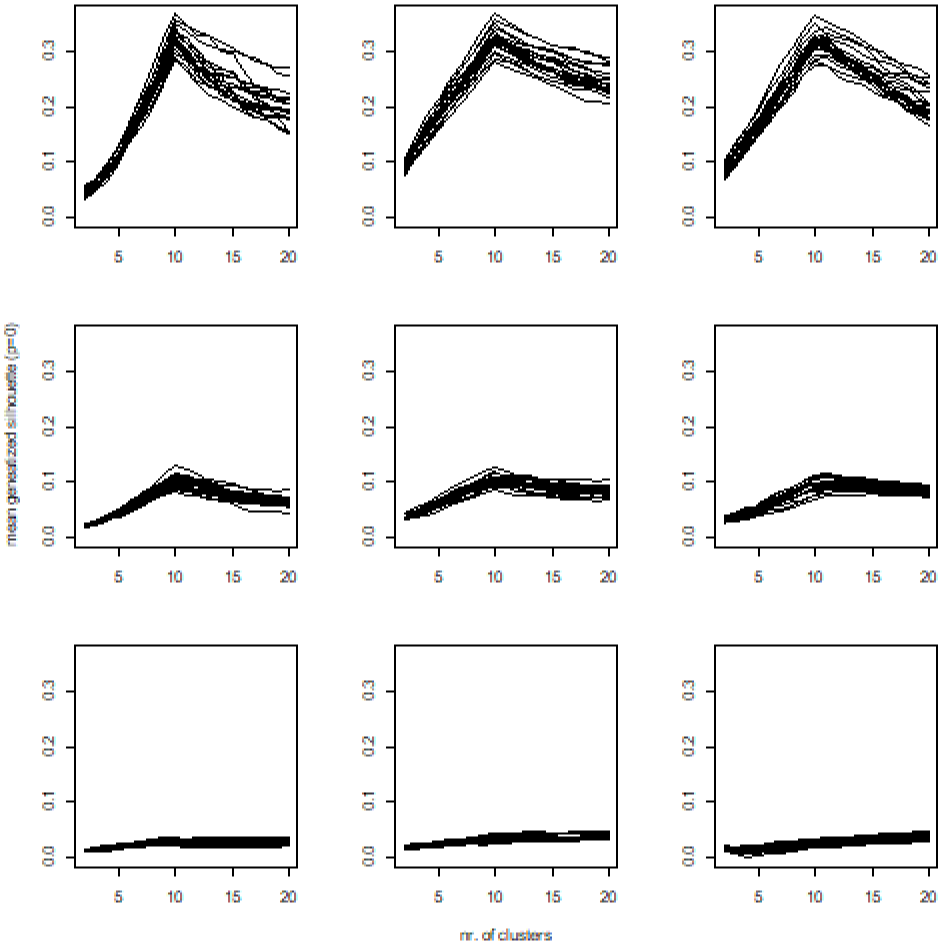


**Figure S1.16** Performance of mean gen. silhouette width (p=0) index versus the number of clusters at different noise levels. Rows indicate noise level (upper is the low noise), and columns are algorithms (left = pam, medium = UPGMA, right = flexible).

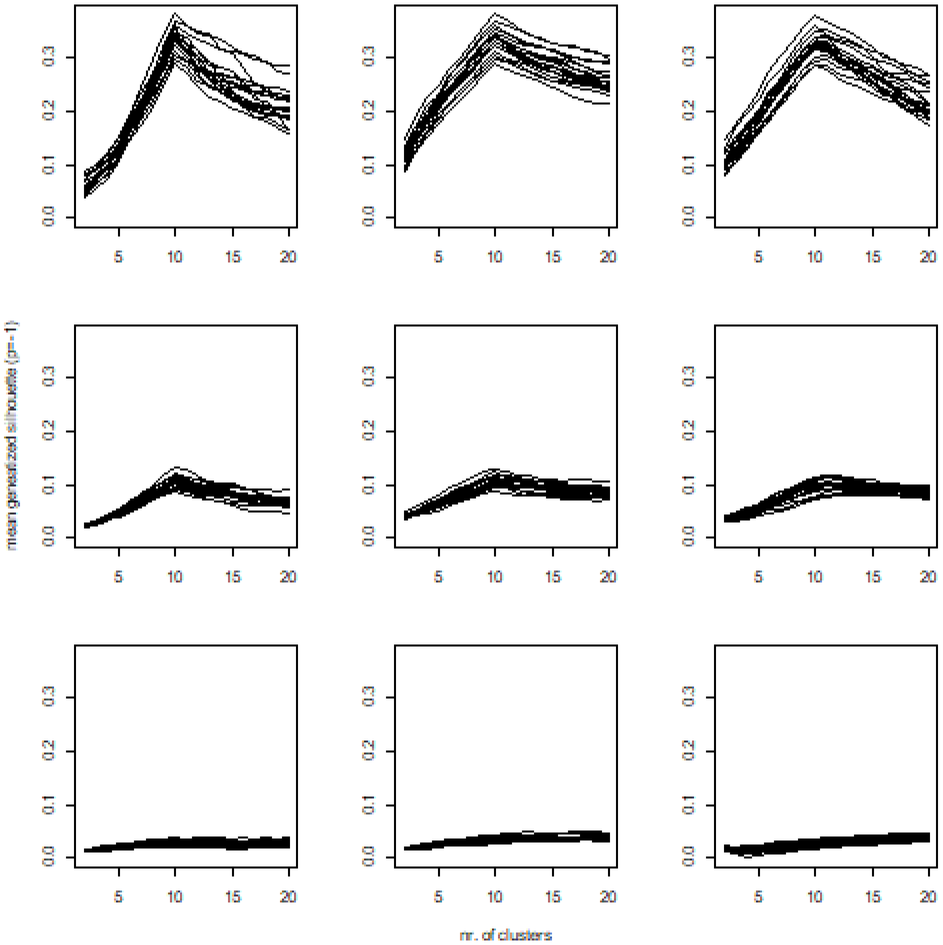


**Figure S1.17** Performance of mean gen. silhouette width (p=1) index versus the number of clusters at different noise levels. Rows indicate noise level (upper is the low noise), and columns are algorithms (left = pam, medium = UPGMA, right = flexible).

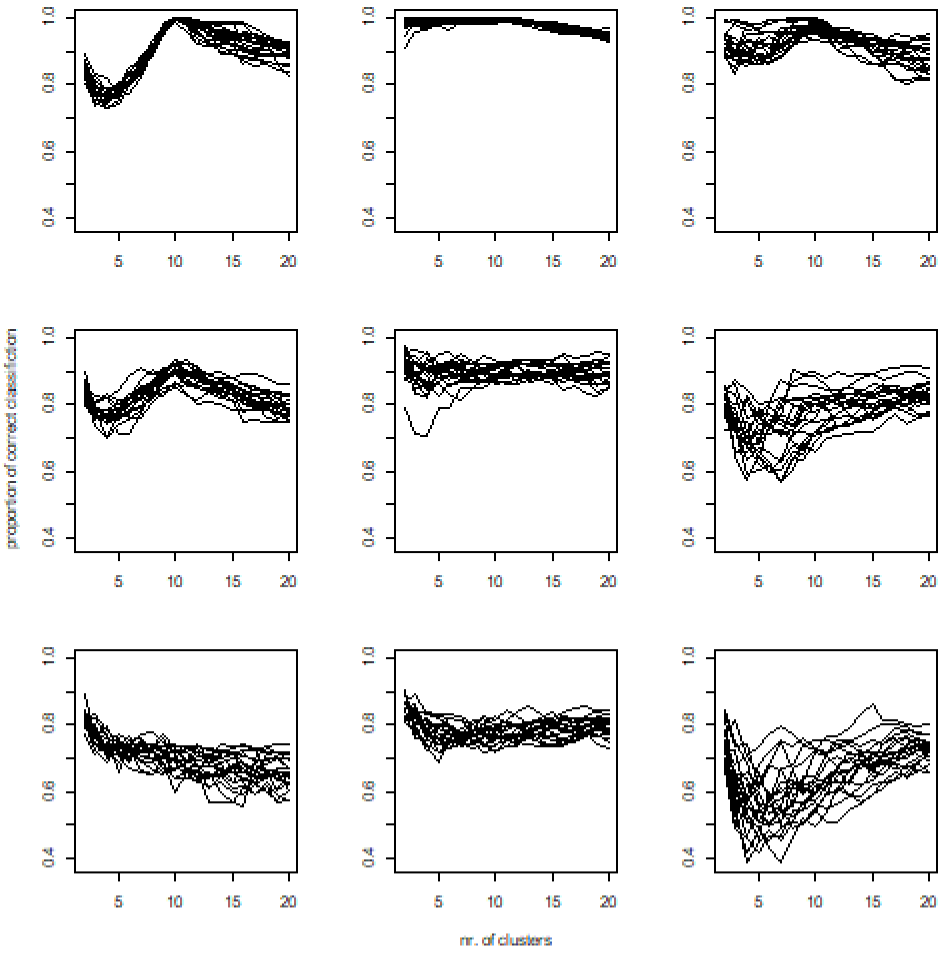


**Figure S1.18** Performance of mean prop. correct classification index versus the number of clusters at different noise levels. Rows indicate noise level (upper is the low noise), and columns are algorithms (left = pam, medium = UPGMA, right = flexible).

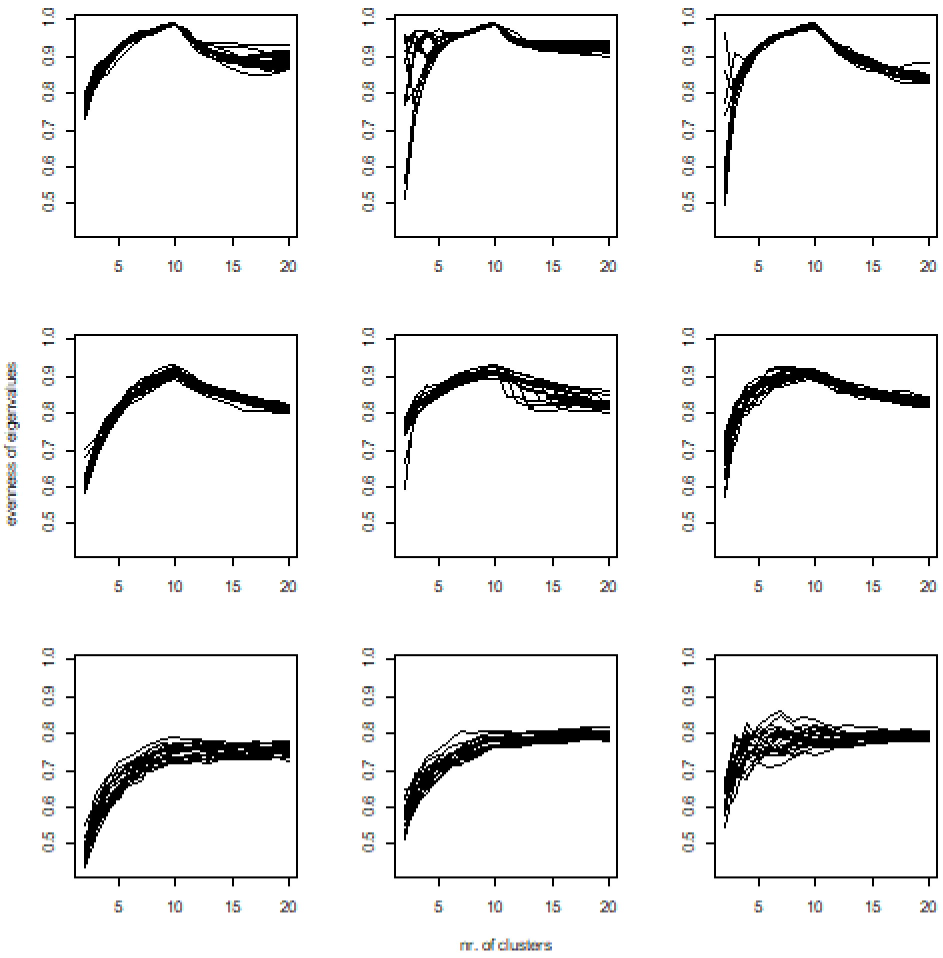


**Figure S1.19** Performance of evenness of eigenvalues index versus the number of clusters at different noise levels. Rows indicate noise level (upper is the low noise), and columns are algorithms (left = pam, medium = UPGMA, right = flexible).

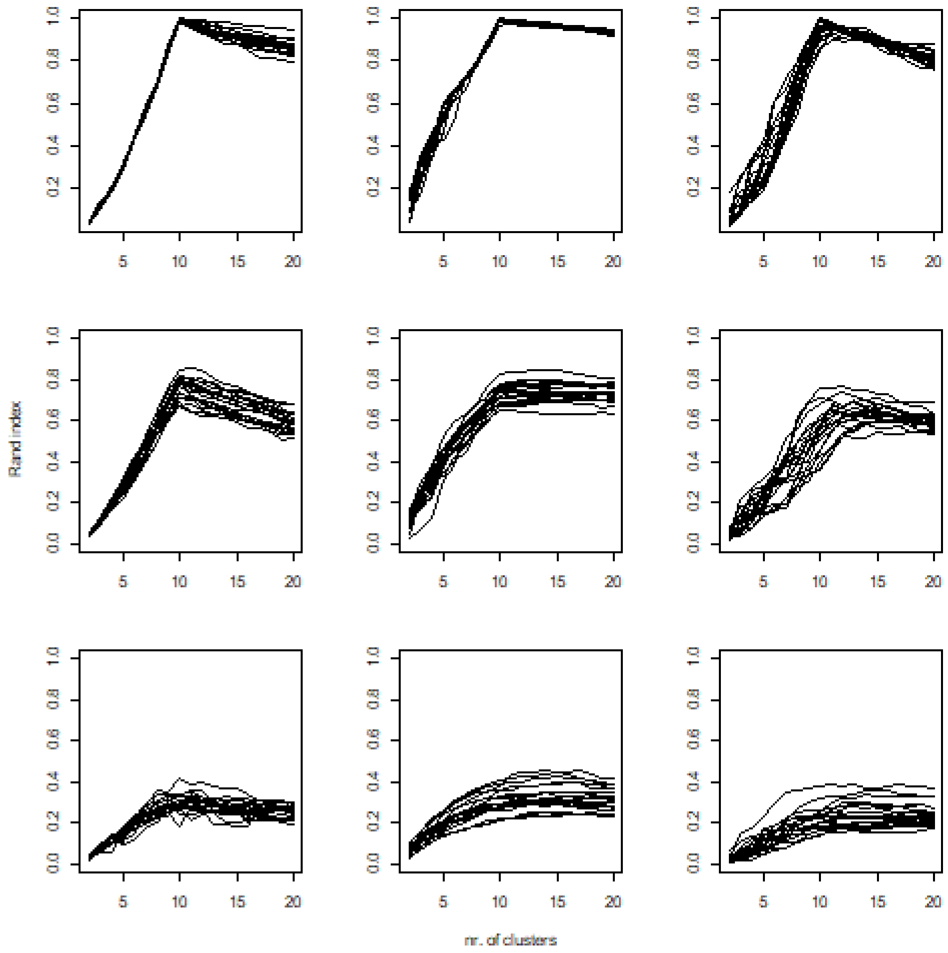


**Figure S1.20** Performance of Rand index versus the number of clusters at different noise levels. Rows indicate noise level (upper is the low noise), and columns are algorithms (left = pam, medium = UPGMA, right = flexible).
